# Unidirectional response to bidirectional selection on body size II Quantitative genetics

**DOI:** 10.1101/2020.01.23.916361

**Authors:** Arnaud Le Rouzic, Clémentine Renneville, Alexis Millot, Simon Agostini, David Carmignac, Éric Édeline

## Abstract

Anticipating the genetic and phenotypic changes induced by natural or artificial selection requires reliable estimates of trait evolvabilities (genetic variances and covariances). However, whether or not multivariate quantitative genetics models are able to predict precisely the evolution of traits of interest, especially fitness-related, life-history traits, remains an open empirical question. Here, we assessed to what extent the response to bivariate artificial selection on both body size and maturity in the medaka *Oryzias latipes*, a model fish species, fits the theoretical predictions. Three lines (Large, Small, and Control lines) were differentially selected for body length at 75 days of age, conditional on maturity. As maturity and body size were phenotypically correlated, this selection procedure generated a bi-dimensional selection pattern on two life history traits. After removal of non-heritable trends and noise with a random effect (’animal’) model, the observed selection response did not match the expected bidirectional response. For body size, Large and Control lines responded along selection gradients (larger body size and stasis, respectively), but, surprisingly, the Small did not evolve a smaller body length, and remained identical to the Control line throughout the experiment. The magnitude of the empirical response was smaller than the theoretical prediction in both selected directions. For maturity, the response was opposite to the expectation (the Large line evolved late maturity compared to the Control line, while the Small line evolved early maturity, while the opposite pattern was predicted due to the strong positive genetic correlation between both traits). The mismatch between predicted and observed response was substantial and could not be explained by usual sources of uncertainties (including sampling effects, genetic drift, and error in G matrix estimates).

## 1 Introduction

Quantitative genetics offer simple and practical models to understand the evolution of complex traits in populations (Falconer and McKay, 1996; Lynch and Walsh, 1997). In practice, these models are used both to analyze past selection response (identifying the factors involved in phenotypic change), and to predict the potential response to selection in a population. The rate of phenotypic change per generation is estimated by multiplying a measurement of the standing genetic (heritable) variation by a measurement of the strength of selection. In the simplest univariate model (the “breeder’s equation”, Lush, 1937), the genetic variation can be quantified by the heritability *h*^2^ (proportion of the phenotypic variance that is heritable between parents and offspring). This setting is convenient when a unique trait is under selection, such as in some selective breeding programs, but becomes rapidly limited when the selection pressure is more complex and targets multiple traits at once. Multivariate models propose a different setting, and quantify evolvability through the “G” matrix of additive genetic (co)variances across traits, and selection through a vector of selection gradients *β* (Lande, 1979; Lande and Arnold, 1983; Blows, 2007; McGuigan, 2006). This approach offers efficient tools to explore theoretically and estimate empirically the properties of multivariate evolution and genetic constraints in complex and integrated biological systems (Cheverud, 1984; Hansen and Houle, 2008; Houle et al., 2017).

Although the data are heterogeneous (various organisms and different kinds of traits) and experimental results lack consistency, the general pattern that seems to emerge from artificial selection experiments is that quantitative genetics may predict short-term direct responses (phenotypic change of a single selected trait) convincingly (Sheridan, 1988; Walsh and Lynch, 2018, p. 606), correlated responses (genetic change in a trait that is genetically correlated to a selected trait, without being the target of selection) at least qualitatively (Gromko, 1995), while the response to multivariate selection (in which the selection gradient affects several traits) may be complex and inconsistent in some cases (Roff, 1997, p. 188, Roff, 2007 for review). In uncontrolled environments, such as in wild populations, even univariate predictions may fail (Merilä et al., 2001). Whether such unconvincing predictions are due to experimental issues, unrealistic assumptions, or flaws in the multivariate quantitative genetics theory, largely remains to be determined. There are indeed many reasons why an experimental selection response could deviate from the theoretical prediction. Some of these mechanisms, such as genetic drift, are compatible with the mainstream theoretical framework, although rarely quantified and accounted for in the analysis of selection responses (Lynch, 1988; Hadfield et al., 2010). Other mechanisms, including environmental trends, genetic × environment interactions, directional epistasis, or scaling effects, are absent from the basic textbook models; including them in specific models is, however, generally possible (Martinez et al., 2000; Le Rouzic et al., 2011; Walsh and Lynch, 2018). More problematic for the standard theory are criticisms concentrating on additive genetic correlations being too crude to capture the complexity of trait associations. This includes claims that understanding the evolution of high-level quantitative traits (such as developmental or morphological traits) cannot be achieved without considering proximal physiological mechanisms (Davidowitz et al., 2016), that the linear assumptions may not hold on complex genotype-phenotype maps (Milocco and Salazar-Ciudad, 2020), or that predictions rely on a poor understanding of the nature and stability of genetic correlations, which could be extremely labile (Gutteling et al., 2007).

We investigated the phenotypic consequences of artificial selection on the medaka fish (*Oryzias latipes*) for a broad set of morphological, physiological, and life history traits, among which two were under direct selection. Wild-caught fish were submitted to 6 generations of truncation selection on fish length at 75 days. The experimental procedure generated three populations; a Large line, in which only large fish were bred, a Small line, in which only small fish were bred, mimicking harvest-like selection regime, and a Control line, in which fish were bred independently from their size. As the experimental design discarded *de facto* immature fish from the breeding pool, all three lines were thus also affected by a selection pressure for early maturity, a trait that was phenotypically correlated with size. Selection was thus essentially bivariate, in divergent directions across lines for body size and in the same direction (but with different intensities) for maturity. After 6 generations of artificial selection, all lines evolved, but phenotypic response did not follow the selection differentials. Fish body size evolved only in the Large line, but not in the Small line, which remained statistically indistinguishable from the Control line (Figure 1). Conversely, the frequency of mature fish did not increase in spite of a positive selection differential in all three lines. These results confirm that anticipating qualitatively and quantitatively the consequences of multivariate selection on fish morphology and physiology cannot be based on the fitness function, but also needs to account for a deeper understanding of the functional and evolutionary relationships between selected traits. The physiological and life history trait changes associated with the selection response are detailed in a companion paper Renneville et al. (2020). Here, we will investigate to what extent multivariate quantitative genetic models, which include explicit genetic covariance components, could explain and predict such a counter-intuitive selection response.

**Figure 1:**
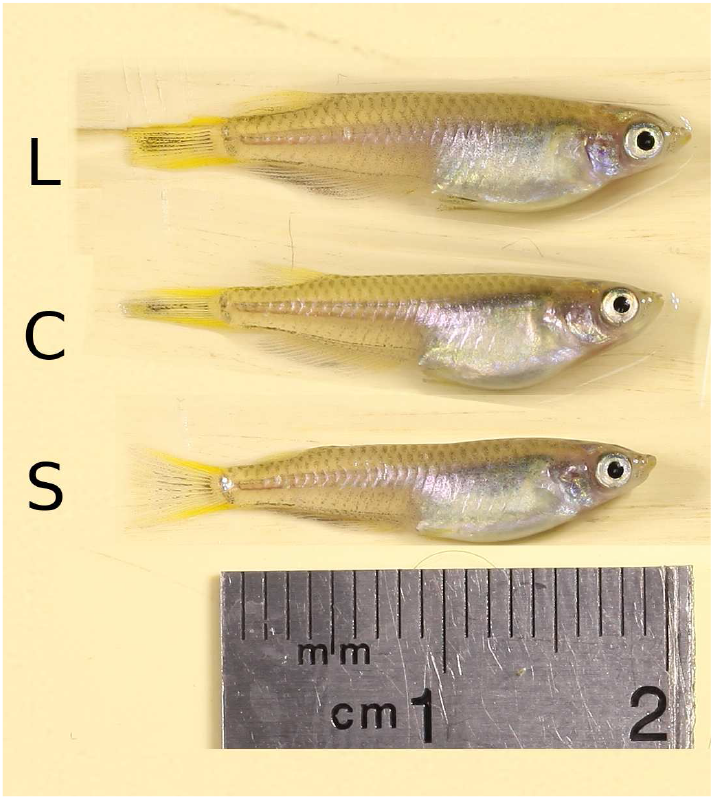
Illustration of the selection response on body size in the medaka *Oryzias latipes* after five episodes of artificial selection. This photomontage displays on the same scale individuals from the F6 generation that are very close to the average length of the Large line (20.95 mm, top), Control line (19.80 mm, middle), and Small line (19.63 mm, bottom).

## 2 Materials and Methods

### 2.1 Biological material and experimental procedure

The initial population was derived from 100 wild adult medaka (*Oryzias latipes*) sampled in June 2011 in Kiyosu (Aichi Prefecture), Japan. Wild-caught fish were stored in outdoor tanks for a year, and about 60 of them were brought to the lab and mated in groups of 3 to 6 individuals to produce a *F*_−1_ generation. From this point, fish were kept as 15-individual full-sib “families” in 3L aquariums under controlled lab conditions (27^*o*^C, 14:10 day light cycles, *ad libitum* feeding). This setting made it possible to record the pedigree in the full experiment. After two generations of random mating (54 and 56 pairs, respectively), the F_1_ mature individuals were split into three breeding groups of 15 males and 15 females (one pair per aquarium) (Large, Small, and Control), and artificial selection was further performed for 6 generations, up to generation F_7_ (Figure 2). The selection procedure involved two steps: an among-family selection step at day 60, and a within-family selection step at day 75. Body size was first measured at 60 days from the pictures of individual fish (length from snout to the base of the caudal fin), and 10 out of ≃15 families were pre-selected in each line based on their average length, after having eliminated low density or high mortality tanks. At about 75 days (in practice 76.7 ± 4.4), pairs were formed by selecting the two largest (respectively, smallest and random) males and females in each family. Immature fish were discarded from the breeding pool. Selection on maturity was necessary to (i) ensure the synchronization of all three lines, (ii) avoid selecting individuals that would never reach the reproductive stage, and (iii) limit sex identification mistakes when making pairs, as sex determination in immature fish requires molecular techniques. More detailed experimental procedures are provided in Renneville et al. (2020).

**Figure 2:**
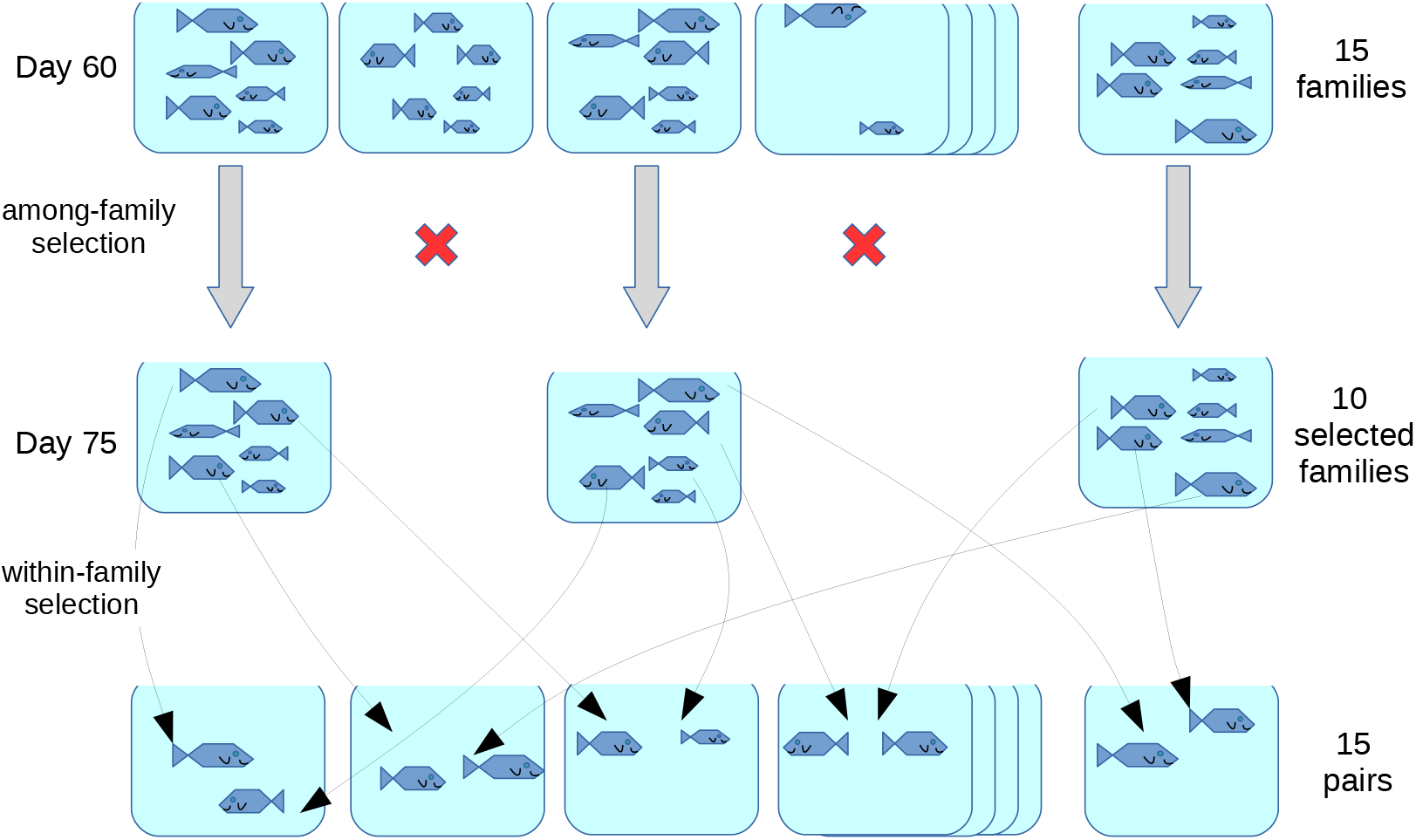
Illustration of the selection protocol. From generation F_1_, three selected lines (Large, Control, and Small lines) were kept independently, with 15 full-sib families per line (in practice: mean ± std. dev. = 15.4 ± 2.7 families per selected line). The density was normalized at 15 fish per tank (in practice: 13.9 ± 5.8). Fish were pictured, measured, and sex-determined at 60 and 75 days. Selection happened in two stages, among families at 60 days, and within families at 75 days. At 60 days, 10 families out of 15 were pre-selected based on two criteria: density (tanks with low fish counts were discarded), and average size. Families discarded from the breeding pool were kept and measured at 75 days, and were thus considered when computing the population mean. At 75 days, breeders were picked within pre-selected families based on two criteria: maturity (immature fish, which sex could not be determined, were never selected) and size (large fish in the Large line, small fish in the Small line, and random fish in the Control line). Fifteen pairs of fish were formed in each line, in a pattern that minimizes inbreeding. The offspring of each pair then constitutes the next generation.

The unavoidable increase in the inbreeding coefficient across generations was limited by a specific procedure. Every generation in all three lines, twenty theoretical pairs of fish (two males and two females from each of the 10 families pre-selected at 60 days) were determined by a computer resampling procedure (selection of the pairing pattern minimizing the median inbreeding coefficient), and this theoretical pairing pattern was followed as close as possible when fish were selected after 75 days. By generation F_7_, assuming no inbreeding in the F_1_ population, the mean inbreeding coefficients were *F* = 0.11 in the Large line, *F* = 0.091 in the Control line, and *F* = 0.085 in the Small line. As the inbreeding coefficient is expected to increase by a factor 1 − 1*/*2*N_e_* every generation, inbreeding population size estimates were about *N_e_* ≃ 27, *N_e_* ≃ 33, and *N_e_* ≃ 35 in Large, Control, and Small lines, respectively. This procedure thus made it possible to maintain an inbreeding effective population size around 30 in all three lines.

### 2.2 Data analysis

The dataset consists in the measurement of body size (Standard Length, further abbreviated Sdl, in mm) at 75 days, the sex, and the maturity (Mat) status for each of the *n* = 5285 fish of the experiment. The father and the mother of each fish was recorded, except for generation F_0_. All the data analysis was performed with R version 4.0 (R core team, 2020). Inbreeding and coancestry coefficients were calculated with the package kinship2 (Therneau and Sinwell, 2015). An archive containing datasets and scripts to reproduce tables and figures is provided as a supplementary file.

#### Selection differentials and gradients

Effective selection differentials were calculated as the difference between the average phenotype of the breeders (weighted by the actual number of surviving offspring) and the average phenotype of the population (Walsh and Lynch, 2018, p. 487). Since all individuals (breeders and non-breeders) were phenotyped, differentials were not affected by sampling, and were considered to be known with certainty (i.e. measurement error was neglected). As the selection procedure involved both among- and within-family selection, two selection differentials were calculated: the among-family differential *S_a_* measures how much the pre-selected families diverge from the population mean, and the within-family differential *S_w_* measures the difference between selected individuals and the average of their respective families. According to Walsh and Lynch (2018, p. 736), the expected response to among (full-sib) family selection is *R_a_* = *h*^2^*S_a_/*2*t*, while the within-family selection response is *R_w_* = *h*^2^*S_w_/*2(1 − *t*), where *h*^2^ is the narrow-sense heritability, and *t* is the phenotypic correlation between sibs (i.e. the proportion of the total phenotypic variance that can be attributed to the family structure, computed as the correlation between all pairs of individuals belonging to the same family). In our experiment, *t* = 0.23 ± 0.14 (mean ± std. dev. among selected lines and generations). The strength of selection was summarized by a composite selection differential *S* = *S_a_/*2*t* + *S_w_/*2(1 − *t*), which is expected to predict the selection response when multiplied by the heritability. Selection gradients, on which the multivariate selection response theory is based, were obtained by multipling composite selection differentials by the inverse of the phenotypic variance-covariance matrix (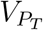 being the phenotypic variance for trait *T*, and 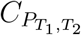 the phenotypic correlation between traits *T*_1_ and *T*_2_):

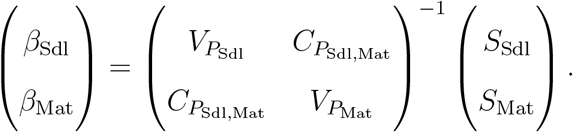

Phenotypic variances and covariances (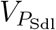, 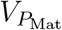, 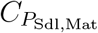 and *t*) were calculated for each line at each generation.

#### Realized heritability

Realized heritability for body size was roughly estimated by regressing the cumulated response to selection (relative to the Control line) on the cumulated selection differential. We implemented the iterated general least square (GLS) regression suggested in Walsh and Lynch (2018, p. 598) to account for the autocorrelation structure in the cumulated selection response *R*. If *S* is the cumulated vector of selection differentials, the variance structure of the regression is a matrix *V* with elements *V_i,j_* = *h*^2^*V_P_* (1*/N* + *i/N_e_*) + (*i* = *j*)*V_P_ /N*, where *N* stands for the size of the population and *N_e_* is an estimate of the effective population size (considered to be *N_e_* ≃ 30 from the inbreeding population size estimate). The (*i* = *j*) term is present only on the diagonal of the *V* matrix. The slope of the GLS regression being *h*^2^ = (*S*^⊺^*V* ^−1^*S*)^−1^*S*^⊺^*V* ^−1^*R* (⊺ stands for matrix transposition), the procedure had to be repeated until convergence since *V* depends on *h*^2^. The error variance in *h*^2^ was then calculated as 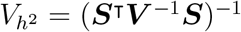.

#### Animal model

Genetic and environmental variance components were estimated from the pedigree with a bivariate mixed-effect linear model framework (’animal’ model) (Lynch and Walsh, 1997; Sorensen and Gianola, 2007; Thompson, 2008), which general setting was as follows. As we were considering two traits, the phenotype of an individual *i* (1 ≤ *i* ≤ *n*) is bivariate 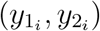, each trait following the classical infinitesimal model in quantitative genetics, e.g. 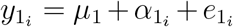, where *μ*_1_ is the grand mean of trait 1 at the first generation (model intercept), *α*_1_i is the additive genetic (breeding) value of individual *i* for trait 1, and *e*_1_i is an environmental (non-heritable residual) deviation. The variance-covariance matrix of all 2*n* breeding values 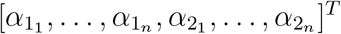 is the Kronecker product G ⊗ A, where G is the 2 × 2 additive variance-covariance matrix between both traits (additive genetic variances on the diagonal, additive genetic covariance off-diagonal), and A, a *n*×*n* square and symmetric matrix, is the genetic relationship matrix (which elements are twice the coefficient of coancestry of each pair of individuals, calculated from the pedigree). In a similar way, the variance-covariance matrix of the 2*n* residuals is E ⊗ I_*n*_, where E is the 2 × 2 environmental variance-covariance matrix between both traits, and I_*n*_ is the identity matrix of size *n*.

This theoretical setting had to be slightly modified to fit our dataset. The second trait (maturity) is the stochastic realization of an underlying probability *p_m_* of maturing before 75 days. Phenotypic values were thus considered to be on the probit scale (which fits with the assumption that maturity is a threshold character). In such a model, the mean and the variance are not independent, and the residual variance cannot be estimated. We also considered additional random effects: an aquarium effect (351 levels) to account for the fact that fish in the same aquarium shared a common environment (in addition of sharing the same parents), and a generation effect (8 levels) to account for inter-generational environmental variation. Both additional random effects were defined by 2 × 2 variance-covariance matrices with three independent parameters (variance for Sdl, variance for Mat, and covariance). Finally, the model considered the inbreeding coefficient as a covariate for both body size and maturity.

The model was fitted in a Bayesian framework with the package MCMCglmm (Hadfield, 2010). Markov chains were run for 10^6^ iterations, with a burn-in of 10^3^, and the state of the chain was stored every 100 iteration. In practice, 20 chains were run in parallel and analyzed together with the tools from the package coda (Plummer et al., 2006). Defining good priors for covariance matrices is notoriously difficult (Gelman et al., 2006; Alvarez et al., 2014). We used inverse-Wishart priors with two degrees of freedom (*ν* = 2) for all variance components, except that the residual variance of the binomial trait *V_E_*(Mat) was fixed to 1. Such priors can be considered as informative compared to the improper *ν* = 0.002 suggestion (de Villemereuil, 2012), but using informative priors was necessary to limit convergence and stationarity issues while remaining denser around zero compared to the *ν* = 3 possibility (uniform marginal distribution for correlations).

In practice, we ran the model twice with a slightly different setting. The “full” model (including all effects and all data) was expected to describe the genetic architecture of both traits during the selection response across all three lines, while a “predictive” model was run to compare theoretical and realized selection response. In order to make prediction independent from observation, the predictive model was run only on data from generations F_0_ and F_1_ (before the divergence among the three lines), and on the Control line from generation F_2_ to F_7_. In addition, inbreeding effects were removed from the predictive model, as the prediction algorithm was not compatible with fixed effects that are not evenly distributed through time.

#### Evolution of breeding values

When partitioning the different variance components, the GLMM procedure makes it possible to estimate the posterior distribution for individual breeding values. In order to avoid potential issues when estimating population genetic parameters from the mean breeding values (Hadfield et al., 2010), we computed the evolution of breeding values for each iteration of the chain, and reported the median (as well as the 95% support interval) genetic trend by averaging breeding values per line and per generation. This trend estimates the evolution of the genetic composition of the population with the influence of confounding factors (generation, aquarium, and inbreeding effects) removed.

We also assessed whether the reported genetic trends could be explained by genetic drift by applying the simulation procedure described in Walsh and Lynch (2018, p. 707) and Hadfield et al. (2010). We ran a simulation for each iteration of the MCMC chain; genetic values of the founders of the pedigree (individuals without identified parents) were drawn in a bivariate normal distribution which variance-covariance was a G matrix from the posterior distribution, and the full pedigree was then filled recursively by sampling breeding values in a bivariate normal distribution centered on the mean parental breeding values, and of variance-covariance G/2.

Since all fish from the same tank share both parents, we used Aquarium effects to catch non-additive (but possible heritable) parental contributions to the offspring phenotype. We computed the heritable maternal effect *m* (Kirkpatrick and Lande, 1989) by regressing the aquarium effect on the mother’s breeding value each iteration of the chain for both traits, and analyzed the distribution of *m* values as a posterior from the model.

#### Evolutionary predictions

The multivariate prediction of the selection response was obtained by applying the Lande & Arnold equation Δ*μ* = G*β* (Lande and Arnold, 1983) on the scale at which the assumptions of the infinitesimal model were the most reasonable (trait scale for body size, and latent probit scale for maturity). In practice, we used the framework proposed by de Villemereuil et al. (2016) implemented in the package QGglmm for R. G matrix estimates were derived from the “predictive” model described above. Selection response was predicted by applying the QGmvpred function recursively over generations, assuming a linear fitness function on the data scale (with gradients *β*_Sdl_ and *β*_Mat_). Every generation, a random bivariate normal deviation of variance G*/N_e_* was added to the population mean to simulate genetic drift (with *Ne* = 25, as approximately estimated from the drift variance in the simulated pedigrees). The predicted response on the latent scale was translated on the data scale for direct comparison with the phenotypic trends by applying the QGmvmean function.

## 3 Results

### 3.1 Selection gradients

Selection gradients were constant and repeatable throughout the experiment (Figure 3 A). The mean realized gradient on Sdl in the Large line was *β*_Sdl_ = 0.56 mm^−1^± s.d. 0.06 (i.e. in average, being 1 mm larger increased relative fitness by 56%), *β*_Sdl_ = −0.33 mm^−1^±0.19 in the Small line, and *β*_Sdl_ = −0.03 mm^−1^ ± 0.14 in the Control line. Although the experimental procedure was identical in all three lines regarding maturity (only mature fish were kept for breeding), the fact that both traits were phenotypically correlated generated different selection gradients. Selection gradient on maturity was positive in the Small line (*β*_Mat_ = 2.35 ± 1.26 expressed in inverse maturity probability, i.e. an increase in 10% in maturity probability raises the relative fitness by 23%), more moderate in Control (*β*_Mat_ = 1.12 ± 0.93) and negative in the Large (*β*_Mat_ = −1.71±1.49) lines. The negative gradient in the Large line in spite of the selection of mature individuals is a consequence of the phenotypic correlation between size and maturity probability (large fish are enriched with individuals that should not have been mature if average sized). When normalized by the phenotypic standard deviation to make them unitless (and thus comparable across traits), gradients were *β*(*σ*)_Sdl_ = 1.39 and *β*(*σ*)_Mat_ = −0.55 in the Large, indicating that selection on body size was dominating. In contrast, in the Small line, normalized gradients were of the same magnitude (*β*(*σ*)_Sdl_ = −0.83 vs. *β*(*σ*)_Mat_ = 0.76). In sum, selection was bivariate and not completely symmetric among selected lines lines; there was no gradient on body size in the Control line but a slight positive gradient on maturity.

**Figure 3:**
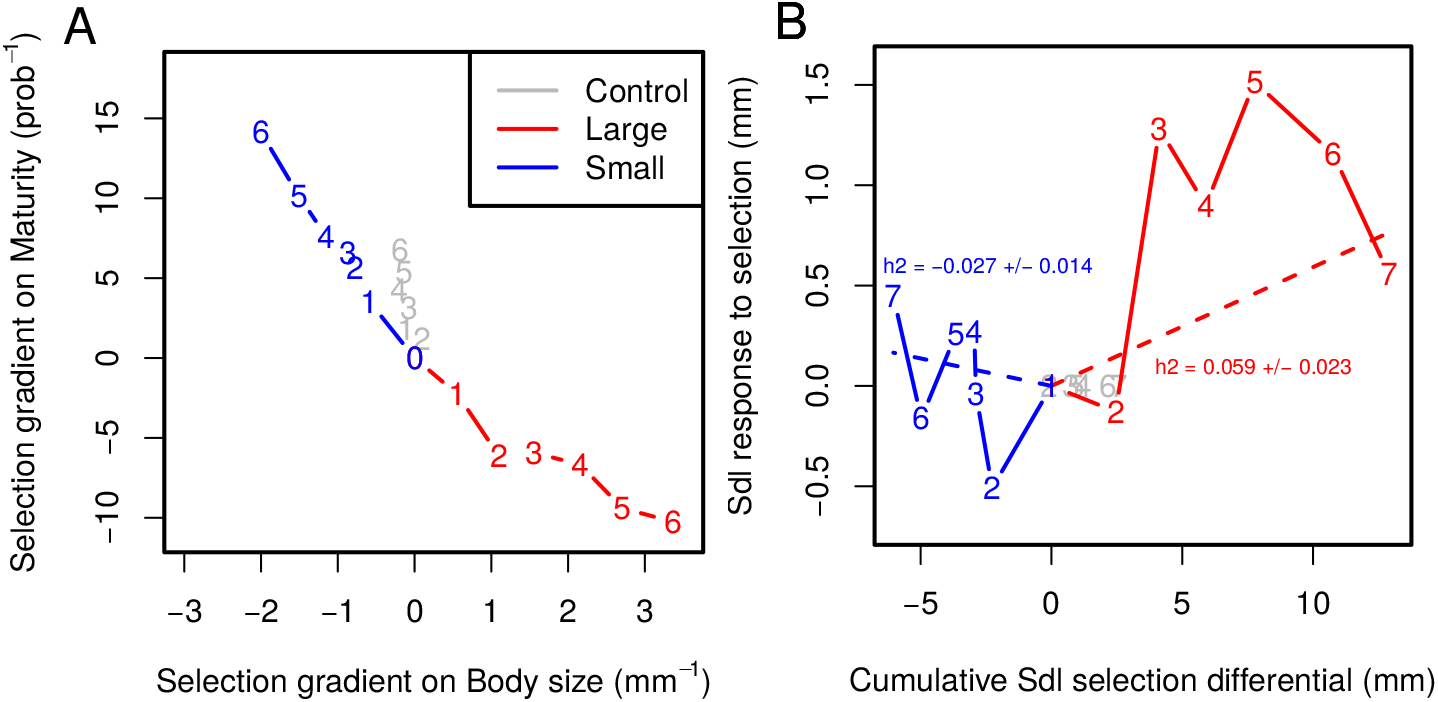
A: Bi-dimensional cumulative selection gradients in all three lines. Gradients were calculated from selection differentials that take into account both within- and among-family selection. B: Cumulative selection differentials vs. cumulative selection response for Body size in Large and Small lines, centered on the Control line. The regression coefficient (calculated by GLS procedure independently for both lines) is an estimate of heritability *h*^2^(± std. err.).

### 3.2 Phenotypic response to selection

The phenotypic response to selection for fish length and maturity is presented in Figure 4. For both traits, time series were characterized by substantial generation-specific effects. Artificial selection has generated a ≃ 1mm difference in body size between the Large and the Control lines (about a 6% increase in length, equivalent to a ≃ 18% increase in mass, illustrated in Figure 1). Virtually all the phenotypic difference was built in two generations of selection, and there was no more phenotypic progress from generations F_3_ to F_6_. The difference between Large and Control lines dropped to less than 1mm in the last generation, but indirect evidence suggest that this was not a stable genetic effect (e.g. the phenotypic difference between Large and Small lines was maintained in fish from later generations, Diaz-Pauli et al. (2019)). Surprisingly, there was no significant difference between the Control and the Small line, i.e. the Small line did not respond to selection on size. Realized heritabilities on body size were positive in the Large line (*h*^2^ = 0.059± s.e. 0.023), and virtually zero in the Small line (*h*^2^ = −0.027 ± 0.014) (Figure 3 B).

**Figure 4:**
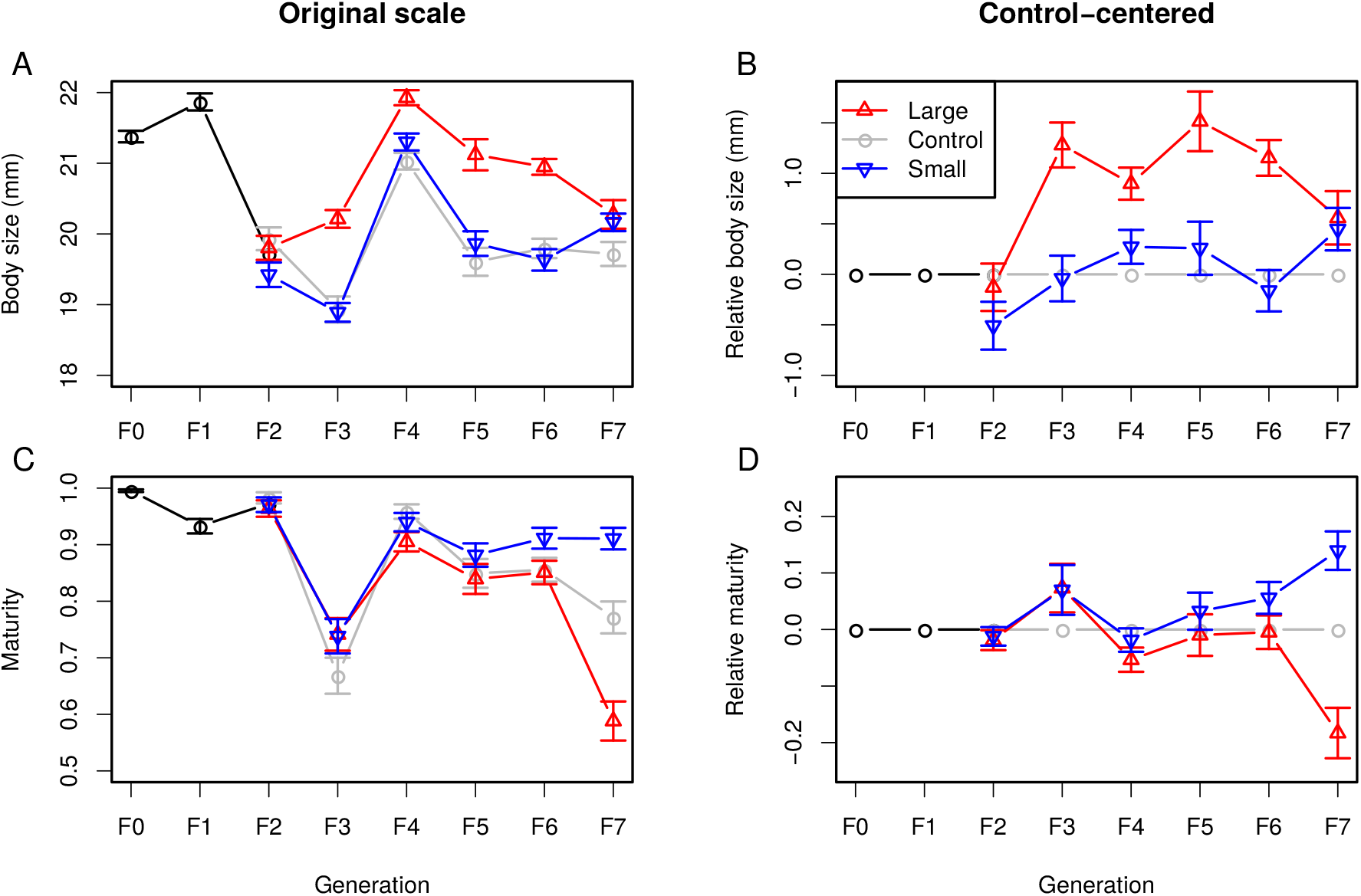
Response to selection for body length (A and B), and maturity (C and D) for all three lines (red: Large, blue: Small, gray: Control). A and C: phenotypic means, B and D: control-centered responses. Generations F_0_ and F_1_ (common to all lines) are drawn in black. Error bars stand for standard errors of the means.

The evolution in maturity was characterized by an irregular decrease, especially in the Control and Large lines. As for fish length, maturity was largely affected by generation-specific effects, especially in F_3_, when maturity dropped from 95% to 75% in all three lines before increasing again to 90% in F_4_. Overall, the general pattern for the bivariate selection response was featured by (i) for body size, a modest selection response in the Large line, but not in the Small line, and (ii) for maturity, a modest divergence from the Control line in two opposite directions (earlier maturation in the Small line, later maturation in the Large line).

### 3.3 Genetic response to selection

A mixed-effect “animal” model was fit to the data, including three (co)variance components: genetic (additive) *G*, aquarium/family (*A*), which is a batch effect that may include heritable maternal effects, and macro-environmental (generation) (*F*), in addition to the residual variance *E*. Variance components were not independent; for instance, aquarium effects and genetic effects were partly confounded, as fish sharing the same aquarium were full sibs.

Table 1 reports the variance components as the median and 95% support interval of the posterior distribution from the MCMC Bayesian analysis. The table also displays heritabilities (*h*^2^ = *G/*(*G* + *A* + *E*)) for both traits as well as correlations (*r*) for all variance components. Model stationarity was unproblematic, but posteriors displayed a substantial amount of autocorrelation (Appendix 1), making it necessary to run long MCMC chains to compensate the poor mixing. The additive genetic variance for body size was *V_G_*(Sdl) = 0.79 mm^2^, and heritability was *h*^2^=0.14, which is more than the realized heritability (although not statistically different). The residual covariance between both traits was substantial, corresponding to a correlation *r_E_* = 0.94 between residual body size and residual probit maturity probability. The genetic correlation was positive, but lower than for the residuals (*r_G_* = 0.62). The posterior distribution of the variance components is illustrated in Figure 5. The effect of inbreeding was negative for both traits (with some statistical support for maturity). Heritable maternal effects were small, slightly negative, and not statistically different from zero.

**Table 1:** Posterior median and 95% support interval from the MCMCglmm model fit, for the full model (left) and the predictive model (right). *V* and *C* stand for variance and covariances, respectively, and subscripts indicate different random effects: additive genetic effects *G*, aquarium effects *A*, residual effects *E*, and generation effects *F*. *f* and *m* correspond to inbreeding and maternal effects, respectively. Sdl is assumed to be Gaussian, and Mat is binomial, on a probit scale (its residual variance *V_E_*(Mat) is fixed to 1 instead of being estimated). Bold-faced variables were the direct output of the model, other variables were estimated by a combination of those (see Methods).

|  |  | Full |  |  | Predictive |  |
| --- | --- | --- | --- | --- | --- | --- |
|  | 2.5% | median | 97.5% | 2.5% | median | 97.5% |
| $\mathbf{V_G}(\text{Sdl})$ | 0.41 | 0.86 | 1.44 | 1.00 | 1.82 | 2.71 |
| $h^2(\text{Sdl})$ | 0.07 | 0.15 | 0.25 | 0.17 | 0.30 | 0.43 |
| $\mathbf{V_G}(\text{Mat})$ | 0.21 | 0.51 | 1.14 | 0.56 | 1.48 | 3.97 |
| $h^2(\text{Mat})$ | 0.12 | 0.25 | 0.43 | 0.25 | 0.46 | 0.67 |
| $\mathbf{C_G}(\text{Sdl, Mat})$ | 0.10 | 0.43 | 0.94 | 0.60 | 1.42 | 2.78 |
| $r_G(\text{Sdl, Mat})$ | 0.31 | 0.65 | 0.81 | 0.72 | 0.87 | 0.94 |
| $\mathbf{V_E}(\text{Sdl})$ | 3.01 | 3.34 | 3.63 | 2.45 | 2.99 | 3.53 |
| $\mathbf{C_E}(\text{Sdl, Mat})$ | 1.61 | 1.72 | 1.81 | 1.36 | 1.57 | 1.75 |
| $r_E(\text{Sdl, Mat})$ | 0.92 | 0.94 | 0.96 | 0.84 | 0.91 | 0.94 |
| $\mathbf{V_A}(\text{Sdl})$ | 1.14 | 1.40 | 1.70 | 0.88 | 1.22 | 1.67 |
| $\mathbf{V_A}(\text{Mat})$ | 0.34 | 0.50 | 0.73 | 0.35 | 0.72 | 1.43 |
| $\mathbf{C_A}(\text{Sdl, Mat})$ | 0.37 | 0.54 | 0.75 | -0.06 | 0.25 | 0.62 |
| $r_A(\text{Sdl, Mat})$ | 0.50 | 0.65 | 0.77 | -0.07 | 0.27 | 0.54 |
| $\mathbf{V_F}(\text{Sdl})$ | 0.41 | 1.05 | 3.64 | 0.57 | 1.62 | 6.05 |
| $\mathbf{V_F}(\text{Mat})$ | 0.60 | 1.53 | 5.46 | 1.08 | 3.26 | 12.88 |
| $\mathbf{C_F}(\text{Sdl, Mat})$ | -0.39 | 0.50 | 2.69 | 0.15 | 1.53 | 6.90 |
| $r_F(\text{Sdl, Mat})$ | -0.29 | 0.45 | 0.85 | 0.09 | 0.72 | 0.94 |
| $\mathbf{f_{Sdl}}$ | -15.74 | -6.78 | 2.31 | | | |
| $\mathbf{f_{Mat}}$ | -18.07 | -10.21 | -2.73 | | | |
| $m_{\text{Sdl}}$ | -0.19 | -0.04 | 0.11 | -0.19 | 0.00 | 0.18 |
| $m_{\text{Mat}}$ | -0.16 | -0.04 | 0.09 | -0.18 | -0.02 | 0.14 |

**Figure 5:**
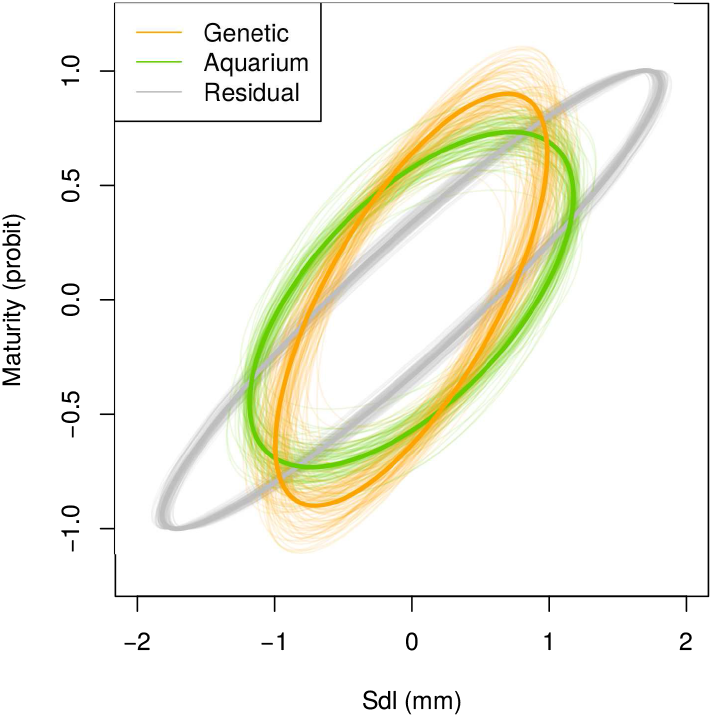
Graphical representation of the posterior distribution of the (co)variance matrix for genetic, aquarium, and residual effects (full model). Thin lines represent individual iterations of the MCMC chain, while the thick line is the mean posterior. The binomial nature of the maturity trait fixes the residual variance to 1.

The generation effect variance was relatively high, especially for Maturity. This generation effect not only captures inter-generational fluctuations, but it also corrects for a general trend in the time series for both traits. Figure 6 displays the genetic (average of breeding values posterior distributions) and non-genetic (from the generation effects) trends from the best model.

**Figure 6:**
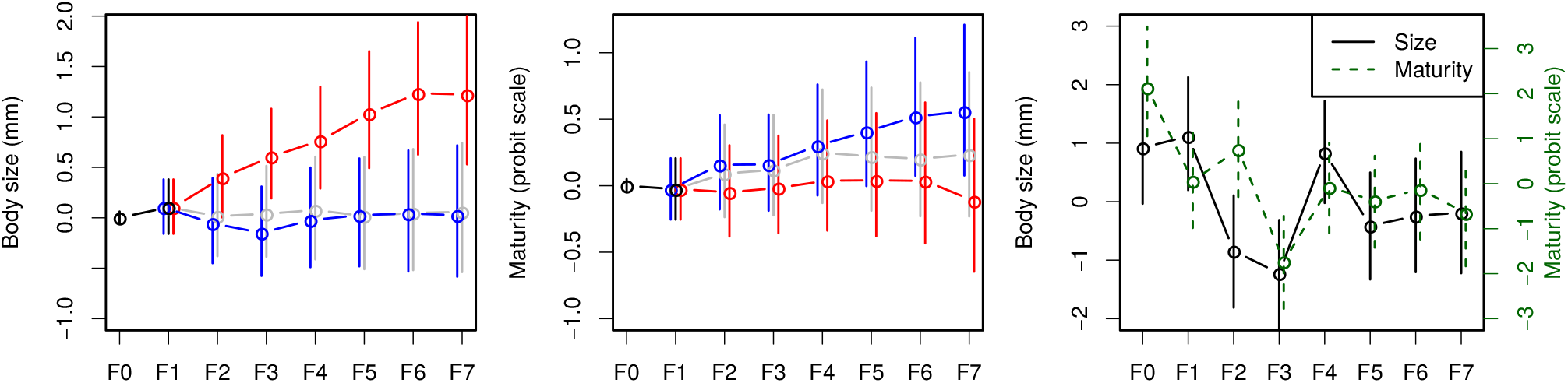
Estimated dynamics of genetic (left and center) and generation (right) effects from the full model. The figure represents the median and 95% support interval over MCMC replicates.

The strength of genetic drift during the selection response was quantified by simulating breeding values based on the estimated **G** matrix in the real pedigree. Figure 7 shows that the 95% support interval of the distribution of average breeding values under genetic drift encompasses the median trend of breeding values in all lines and traits except for the body size of the Large line, which exceeds the drift interval for the last 3 generations. As the variance across replicates is expected to increase by a factor *V_A_/N_e_* each generation, this procedure made it possible to estimate a drift effective population size in each line, which was *N_e_* = 23, *N_e_* = 20, and *N_e_* = 25 in Control, Large, and Small lines (averaged over both traits), respectively.

**Figure 7:**
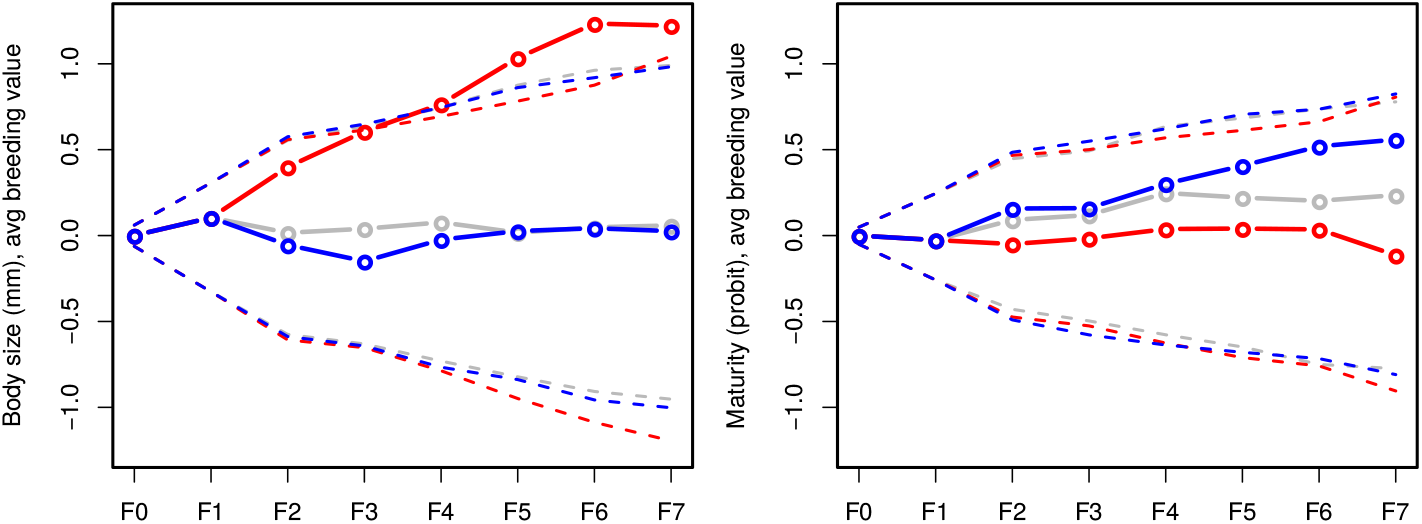
Estimation of the genetic space that is reachable by genetic drift along the selection response (95% support interval: hyphenated lines) (simulated pedigrees with **G** matrices sampled from the posterior distribution) vs. estimated genetic trends (plain lines).

### 3.4 Selection response prediction

In addition to providing the theoretical framework to design statistical models for the estimation of variance components, quantitative genetics also aims at predicting the selection response from the genetic architecture of phenotypic traits. In practice, testing the predictive power of such models requires a specific protocol, as genetic variance components need to be estimated from the starting population (from e.g. an experimental cross design) and compared to the realized selection response. Although we do not have access to a direct measurement of additive variance components in the starting population here, we estimated the **G** matrix by fitting the animal model on the Control line individuals (including F_0_ and F_1_ generations), and compared the predicted selection response to the observed response from the selected lines (no overlap between both datasets). Even when considering uncertainties due to the estimation procedure as well as genetic drift, there was no overlap between predicted and observed evolution for body size (insufficient realized selection response, Figure 8A), while the prediction was even not in the correct direction for maturity (Figure 8B). For both traits, the mismatch was largely supported statistically for the Large line, and on the edge of the 95% support interval for the Small line. The bivariate representation of the predicted vs. observed selection response (Figure 8C) shows that the gradient were oriented close to the line of most evolutionary resistance of the **G** matrix, the predicted response was close to the line of least evolutionary resistance (main axis of the **G** matrix), while the observed response was tiny and inconsistent.

**Figure 8:**
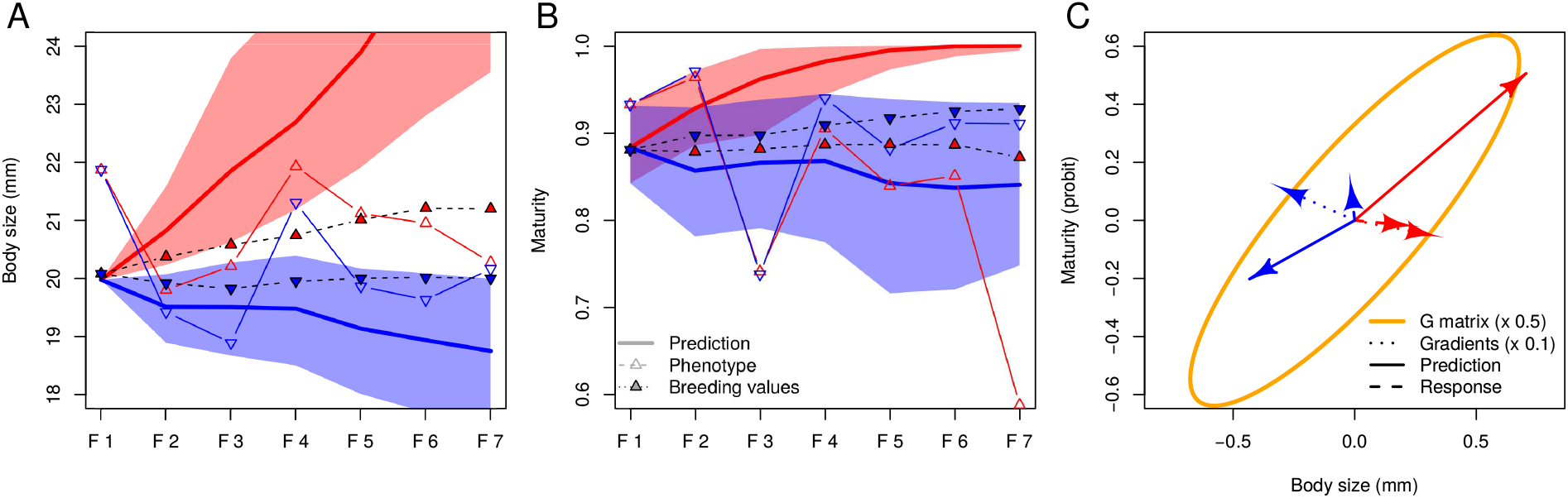
Predicted vs. realized selection responses. The expected selection response was simulated by applying the Lande – Arnold equation Δ*z* = G*β* recursively over six generations, using the selection gradients *β* estimated in Figure 3 and G matrices from the posterior distribution of the animal model applied on data from the Control line (so that the realized selection response was not part of the G matrix estimate). Genetic drift was simulated by applying a random deviation of variance G*/N_e_* with *N_e_* = 25 every generation. A and B: predicted time series for body size and maturity, respectively. Plain lines: predicted responses (Large line in red, Small line in blue); open symbols: actual phenotypic response; hyphenated lines and filled symbols: median genetic response estimated from Figure 6. Shaded areas represent the 95% support interval. C: bivariate average gradients (dotted lines), average predicted responses (plain lines) and average genetic responses (hyphenated lines).

## 4 Discussion

Artificial selection has long been proven to be an efficient way to simulate evolutionary processes in controlled conditions (Hill and Caballero, 1992; Conner, 2003). Here, we applied a classical truncation selection scheme, with substantial improvements compared to classical mass breeding experiments: (i) we kept track of the pedigree during the whole experiment, (ii) crosses were optimized to limit inbreeding, which kept the effective inbreeding population size above *N_e_* = 27 in all three lines, (iii) we recorded fecundity and mortality rates in all families, making it possible to evaluate the potential strength of natural selection, (iv) we raised a control line in the same conditions as selected lines, which helps distinguishing non-genetic and genetic trends, and (v) we selected explicitly on both body size and maturity, and considered both life history traits simultaneously in our analysis. The main drawback of this approach is an increased cost and human power involved, which necessarily limits the size of the experiment in terms of replicates (three lines in total) and duration (almost 3 years, for 8 generations and 6 episodes of selection, which may not be conclusive for low-evolvability traits). Nevertheless, in spite of such logistic limitations, our *>* 5000 fish pedigree displayed sufficient statistical power to (i) evidence a mismatch between predicted and observed selection response, and (ii) discard genetic drift as the driver of this mismatch. In sum, the size of the experiment might be too limited to fully understand and generalize how life-history traits respond to complex multivariate selection, but is sufficient to conclude that the observed response does not follow quantitative genetics predictions.

### 4.1 Departure from theoretical predictions

#### Non-genetic effects

In spite of the tight control over environmental conditions (constant food, lighting, water quality and temperature), the data analysis highlighted a substantial amount of generation-specific effects that obscured the genetic selection response. Generations F_0_, F_1_, and F_4_ appeared to be substantially “better” (larger body size and higher maturity frequency) than e.g. F_3_ and F_7_. The fact that “good” generations were closer to the beginning of the experiment tends to generate an overall decreasing trend, which was difficult to interpret.

A candidate explanation relies on an increase in inbreeding, which is unavoidable in such an experiment. Inbreeding coefficients were included as covariates in the model, and caught part of the trend for both traits (the negative effect of inbreeding on maturity — but not on body size — being statistically supported, Appendix 2). However, a causal link between inbreeding and measured traits remains dubious. The optimized pairing protocol limited the increase in inbreeding below 10% from generations F_0_ to F_7_, which is unlikely to generate inbreeding depression. Furthermore, we found no correlation between phenotypic traits and inbreeding coefficient within generations. Finally, the residual environmental trend on body size and maturity remained negative even after accounting for the effect of inbreeding in the model, suggesting an inbreeding-independent phenomenon.

#### Evolutionary trends

One of the most unexpected result of this experiment was the lack of response to selection on body size in the Small line, in spite of a substantial and consistent selection gradient. This lack of phenotypic response was confirmed by the breeding value predictions from the animal model.

The inaccurate predictions could be due, to some extent, to inconsistent variance components in the experiment. This possibility was addressed by fitting the model on different generations or different selected lines (Appendix 3), which should provide compatible posterior distributions of the assumptions of the infinitesimal model were holding. Excluding the Control line decreases substantially the estimated additive variance for both traits, which is consistent with the mismatch between the observed selection responses and the evolution predicted from the Control line. Walsh and Lynch, 2018, p. 611 proposed a list of 13 possible explanations for asymmetric selection responses, which we tried to address as thoroughly as possible (Table 2).

**Table 2:**
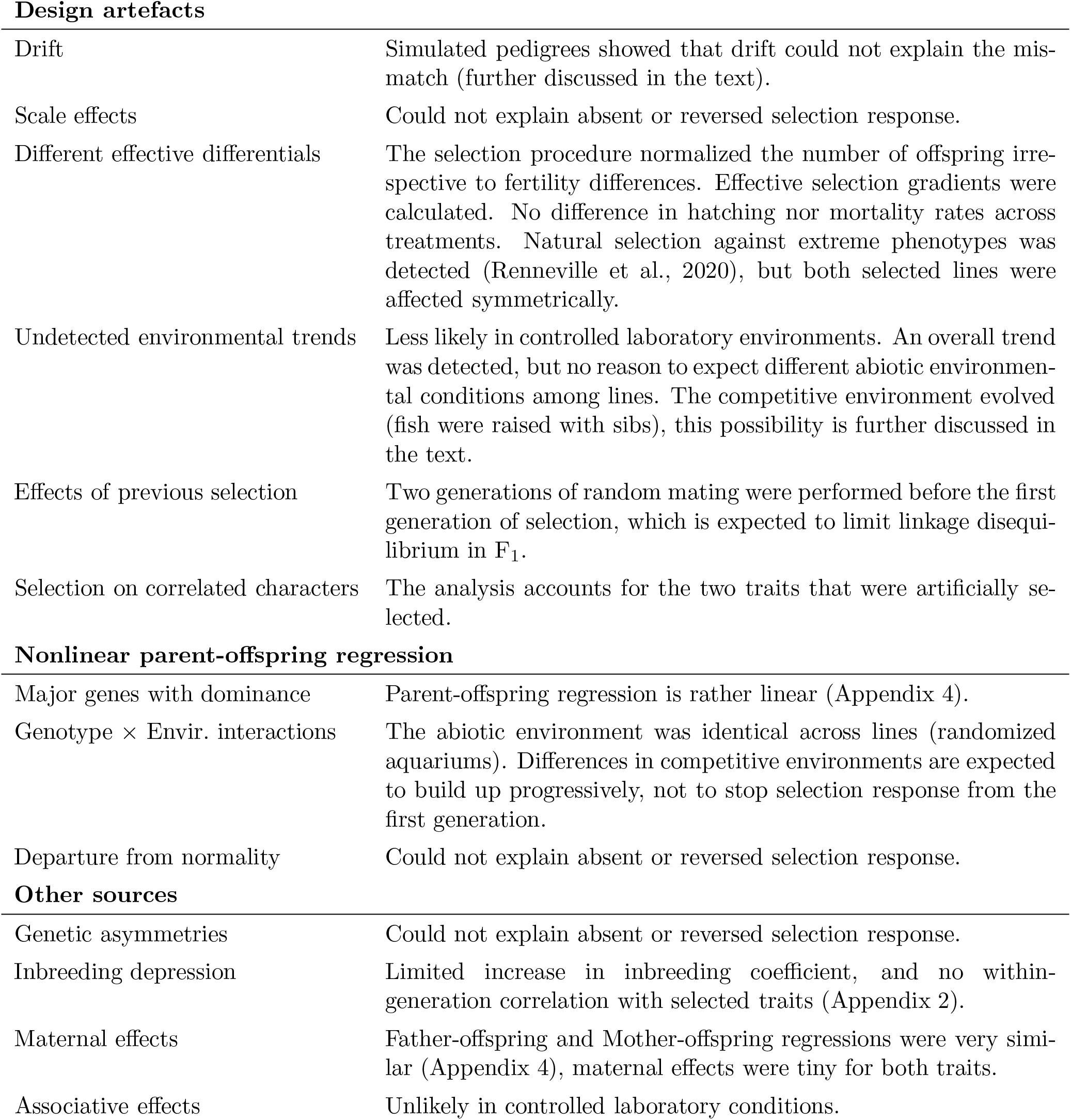
Why do quantitative genetics predictions fail? Thirteen possible explanations for asymmetric responses, as proposed by Walsh and Lynch (2018).

|  |  |
| --- | --- |
| Drift | Simulated pedigrees showed that drift could not explain the mismatch (further discussed in the text). |
| Scale effects | Could not explain absent or reversed selection response. |
| Different effective differentials | The selection procedure normalized the number of offspring irrespective to fertility differences. Effective selection gradients were calculated. No difference in hatching nor mortality rates across treatments. Natural selection against extreme phenotypes was detected (Renneville et al., 2020), but both selected lines were affected symmetrically. |
| Undetected environmental trends | Less likely in controlled laboratory environments. An overall trend was detected, but no reason to expect different abiotic environmental conditions among lines. The competitive environment evolved (fish were raised with sibs), this possibility is further discussed in the text. |
| Effects of previous selection | Two generations of random mating were performed before the first generation of selection, which is expected to limit linkage disequilibrium in $F_1$. |
| Selection on correlated characters | The analysis accounts for the two traits that were artificially selected. |
| **Nonlinear parent-offspring regression** |  |
| Major genes with dominance | Parent-offspring regression is rather linear (Appendix 4). |
| Genotype × Envir. interactions | The abiotic environment was identical across lines (randomized aquariums). Differences in competitive environments are expected to build up progressively, not to stop selection response from the first generation. |
| Departure from normality | Could not explain absent or reversed selection response. |

|  |  |
| --- | --- |
| Genetic asymmetries | Could not explain absent or reversed selection response. |
| Inbreeding depression | Limited increase in inbreeding coefficient, and no within-generation correlation with selected traits (Appendix 2). |
| Maternal effects | Father-offspring and Mother-offspring regressions were very similar (Appendix 4), maternal effects were tiny for both traits. |
| Associative effects | Unlikely in controlled laboratory conditions. |

None of these explanations, taken individually, was particularly convincing. The potential for genetic drift to generate unexpected evolutionary patterns is substantial, and ruling out the influence of drift in laboratory experiments in notoriously difficult (Lynch, 1988), especially with unreplicated selection lines. Nevertheless, we were able to exclude genetic drift as a major explanatory factor of the observed response: effective population sizes were rather large for such an experiment (*N_e_* ≃ 25), and the support intervals predicted for genetic drift from simulated pedigrees (Figure 7) were smaller than the the predicted response mismatch(Figure 8).

The fact that Large and Small lines were set in different competitive environments appears as an appealing explanation for the asymmetric response. Indeed, in order to keep track of the pedigree, fish were raised in the same tank as their full-sibs. If small or large body size was partly correlated to any competitive behavioral trait, the environment was varying during the experiment, as fish from the Large line were competing with better competitors every generation. This mechanism could have biased the selection response estimates, and explain the lower-than-expected body-size response to selection. However, this possibility is dismissed by indirect evidence from an independent phenotyping experiment in which fish from the present selection experiment at later generations were raised in individual tanks (Diaz-Pauli et al., 2019). In absence of competition, the genetic difference between the Large and Small lines was not larger than we found here, indicating that removing competition did not magnify the phenotypic effects of selection on medaka body size.

Finally, natural selection against a small body size appears unlikely due to several consistent observations: (i) the Control line was affected by a general phenotypic (but not genetic) trend towards smaller body size, not larger; (ii) differences in fertility and mortality rates were limited by the experimental procedures, as the number of progeny per fish pair was normalized whenever possible; (iii) the difference between non-weighted selection differentials and effective differentials was reasonably small, (iv) natural selection on body length could be indirectly estimated (Renneville et al., 2020), and appeared to be stabilizing (decrease in the number of offspring for both small and large fish).

Selection response on maturity followed an even more unexpected pattern. In the Large line, selection on body size was dominating, leading to an expected increase in maturity due to the strong genetic correlation in spite of a weak negative selection gradient on maturity. In the Small line, selection on both traits was of the same magnitude, and the expected trend for maturity was slightly decreasing. Yet, both phenotypic and genetic trends were opposite to the prediction. Here again, the mismatch was large enough to discard genetic drift as the only mechanism.

### 4.2 Consistency with previous results

Due to their close relationship with fitness, life history traits are often suspected to behave differently from other (morphological, physiological, behavioral) characters. Their heritability tends to be lower (Price and Schluter, 1991; Roff, 1997) (probably due to a larger residual variance rather than a low genetic variance, Houle (1992)), and the correlation pattern among fitness-related traits have long been a matter of debate (Lande, 1982; Reznick, 1985; Houle, 1991), with few strong theoretical predictions about the sign of genetic covariances among life history traits. Meta-analyses support the idea that correlations between life history traits range between −1 and 1 depending on traits and organisms, being slightly positive on average, but lower than between other kinds of traits (Roff, 1996). The fact that phenotypic and genetic correlations generally match is well-established empirically (Cheverud, 1988; Kruuk et al., 2008), although the underlying reasons are unclear. Our results featuring a substantial positive genetic correlation between growth and maturity, associated with a strong residual correlation, are thus not unexpected.

Less expected was the lack of response to directional selection in the Small line. Fish artificially-selected for large or small size generally respond to selection in both directions (Diaz-Pauli et al., 2014), symmetrically (as in the Atlantic silverside, Conover & Munch, 2002, or in zebrafish, Amaral and Johnston, 2012) or slightly asymmetrically with a slower response in the Small line (in guppy, van Wijk et al., 2013). In the only experiment in which maturity was probably selected together with size (zebra fish, Uusi-Keikkilä et al., 2015), the response was complex and asymmetric (no size change and later age maturation in the Large line, smaller adult size and maturation at a smaller size — but not age — in the Small line).

### 4.3 Consequences on the response of life history traits to selection pressure

One of the most appealing applications of quantitative genetics outside of their original plant and animal breeding field is related to the prediction of the evolutionary consequences of human activity and/or environmental change on natural populations (Shaw, 2019). For instance, size-selective harvesting induces direct selection pressures on body size, and reduces life expectancy, which generates complex selection pressures on correlated life history traits (including growth rate, fertility, survival, and age at maturity) (Heino et al., 2015). Long-term evolutionary trends towards smaller body size, earlier maturity, and as a consequence, lower fecundity are frequent in highly-harvested species (Trippel, 1995; Law, 2000). It is therefore increasingly recognized that fisheries management programs should account for evolutionary change in life history traits (Kuparinen and Merilä, 2007; Fenberg and Roy, 2008; Laugen et al., 2014).

Generalizing results obtained from model species in laboratory to wild species of interest is not straightforward, as differences in environment may condition trait means, trait variances, and genetic correlations (Gutteling et al., 2007; Postma et al, 2007). The development of the ‘animal’ statistical model makes it possible to evaluate genetic components from observations in unmanipulated wild populations (Kruuk, 2004; Wilson et al., 2010). However, this approach remains particularly sensitive to e.g. gene-by-environment interactions, and the causal factors of observed trends may be difficult to identify formally (Postma and Charmantier, 2007; Walsh and Lynch, 2018). In contrast, controlled experiments (typically, complex breeding schemes) can only be carried out in laboratory conditions, and experimental approaches are often the only way to study key questions in population management, even when studying complex marine ecosystems (Suquet et al., 2005).

Accounting for evolutionary response management strategies in wild populations generally relies on standard models in ecology and quantitative genetics, which assume that evolution can be reliably predicted when genetic trait variances and covariances are known (Diaz-Pauli et al., 2014), which is generally not the case. Here, we show that such standard expectations may not be fulfilled, which questions the possibility to apply general recipes. Our results support the idea that bivariate selection response is hardly predictable even in a controlled environment, which questions the robustness of fishery management genetic models. Although we lack a clear explanation about why some heritable characters may not evolve when selected together, this phenomenon may decrease our confidence in the estimates of phenotypic trajectories for populations under anthropic pressure.

## Supporting information

Data files and analysis scripts

## Data accessibility statement

The data and the R scripts used for the analysis are fully available and provided as supplementary material.

## Author contributions

All co-authors participated to the data collection. ALR, CR, and EE cured and analyzed the data, ALR and EE wrote the manuscript. All co-authors agreed on the final version of the manuscript.

## Acknowledgements

We thank Christophe Pélabon for insightful discussions and for his invitation to the Center of Advanced Study (CAS) in Oslo. Many thanks to Pierre de Villemereuil for his availability and kind help for the use of the QGglmm package. We thank both reviewers for their insighful and benevolent suggestions.

## Appendix 1 Model convergence

## Autocorrelation and effective size

|  | Lag 1 | Lag 5 | Lag 10 | Lag 50 | Lag 100 | Lag 500 | Eff. Size |
| --- | --- | --- | --- | --- | --- | --- | --- |
| V[G](Sdl) | 0.99 | 0.96 | 0.93 | 0.72 | 0.52 | -0.08 | 1312.64 |
| V[G](Mat) | 0.99 | 0.97 | 0.94 | 0.79 | 0.64 | 0.10 | 466.69 |
| C[G](Sdl, Mat) | 0.99 | 0.97 | 0.94 | 0.75 | 0.59 | 0.02 | 679.69 |
| V[E](Sdl) | 0.95 | 0.85 | 0.79 | 0.55 | 0.40 | -0.06 | 1461.35 |
| C[E](Sdl, Mat) | 0.96 | 0.88 | 0.82 | 0.57 | 0.42 | -0.02 | 1526.90 |

Autocorrelation for the random effect parameters was assessed with the autocorr.diag() function from package coda. The effective size (sample size adjusted for autocorrelation) was evaluated with the function effectiveSize() from the same package.

## Stationarity

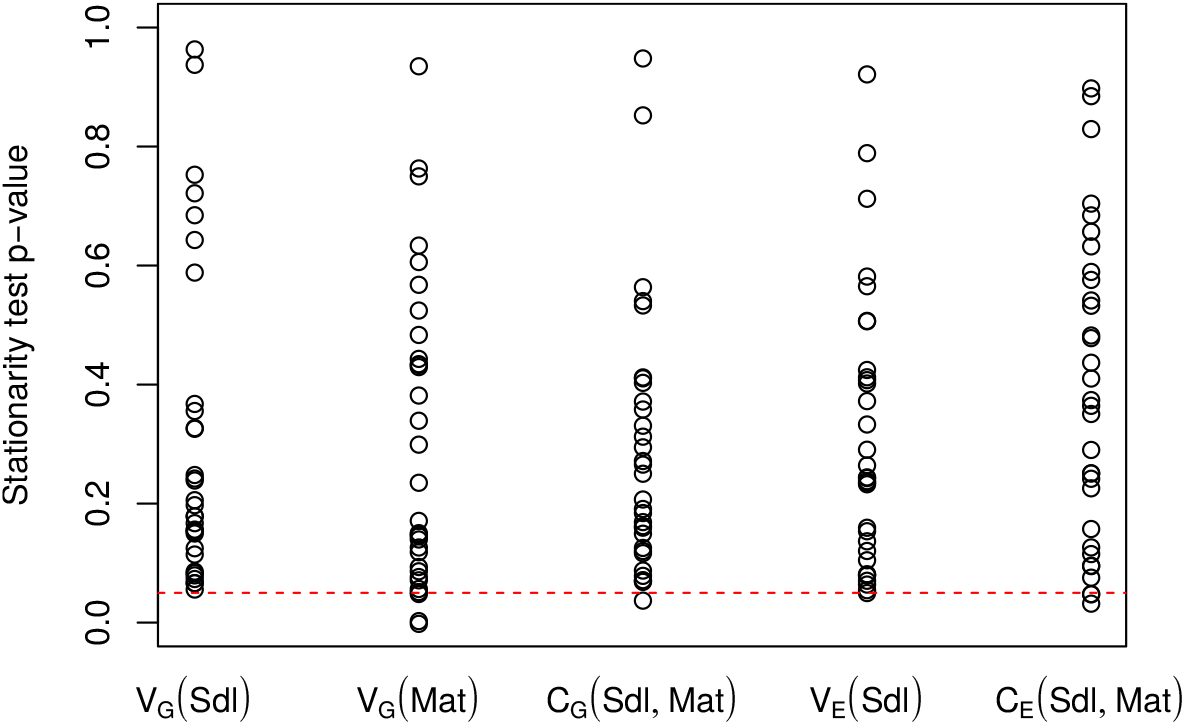

The vast majority of MCMC chains passed the Heidelberg stationarity test implemented in the heidel.diag() function from the coda package (null hypothesis *H*_0_: the chain is stationary at least over its last half, the dashed line illustrates the 5% threshold).

## Appendix 2 Inbreeding

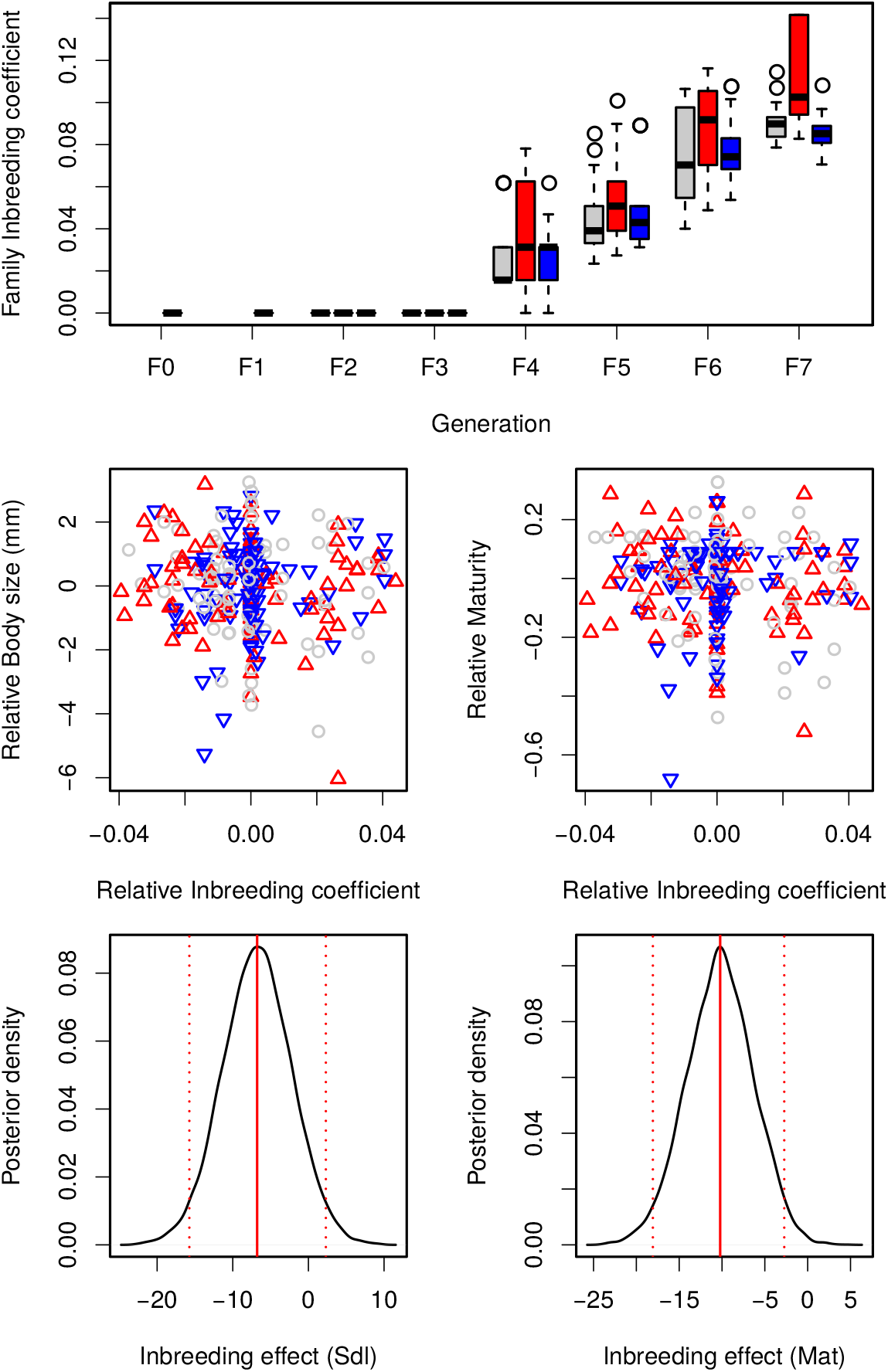

Top: Distribution of the inbreeding coefficients (calculated from the full pedigree) across families in the course of the experiment (assuming no inbreeding in F_0_). Middle: relationship between the inbreeding coefficient of families (normalized by the average of the line each generation) to phenotypic traits (centered on the line and generation mean). None of these regressions were statistically significant. Bottom: posterior distribution of the inbreeding effects on Sdl and maturity (vertical lines indicate 2.5%, 50%, and 97.5% quantiles).

## Appendix 3 Model fitting on partial datasets

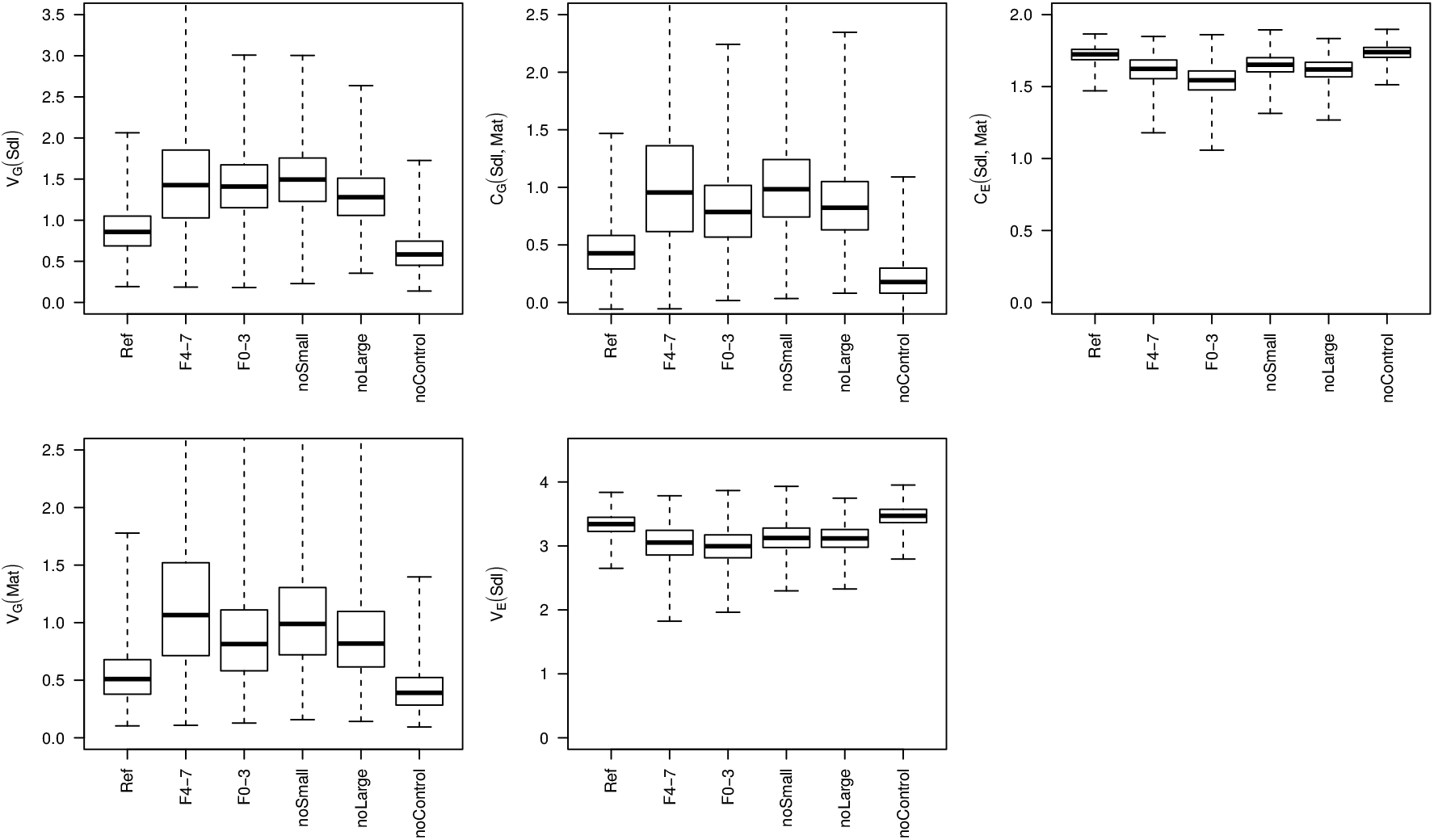

The animal model estimates variance components in the starting population (F_0_) accounting for drift and selection in subsequent generations. As a consequence, if the assumptions of the infinitesimal model hold, fitting the model on partial datasets should not affect the estimates (while the posterior distribution is expected to be wider due to the decrease in information). We split the dataset according to (i) generations (fitting the model on generations F_0_ to F_3_, and from F_4_ to F_7_), and (ii) to the selected line (Large, Small, and Control lines), fitting the model excluding sequentially each line. In the figure, “Ref” stands for the posterior when including all the data, and boxplots represent the full range of the posterior distributions and their quartiles. The estimates for genetic variances and covariances increased for most sub-datasets, and residual variances and covariances decrease accordingly. The most straightforward explanation is that the parameters estimated from the full dataset result from a compromise between early/late generations and selection lines, and that the goodness of fit of the model increased when fitted on partial data. Note that most posterior distributions largely overlap (no posterior distributions differ significantly from the reference), suggesting that the estimated parameters remain meaningful.

## Appendix 4 Parent-offspring regression for body size

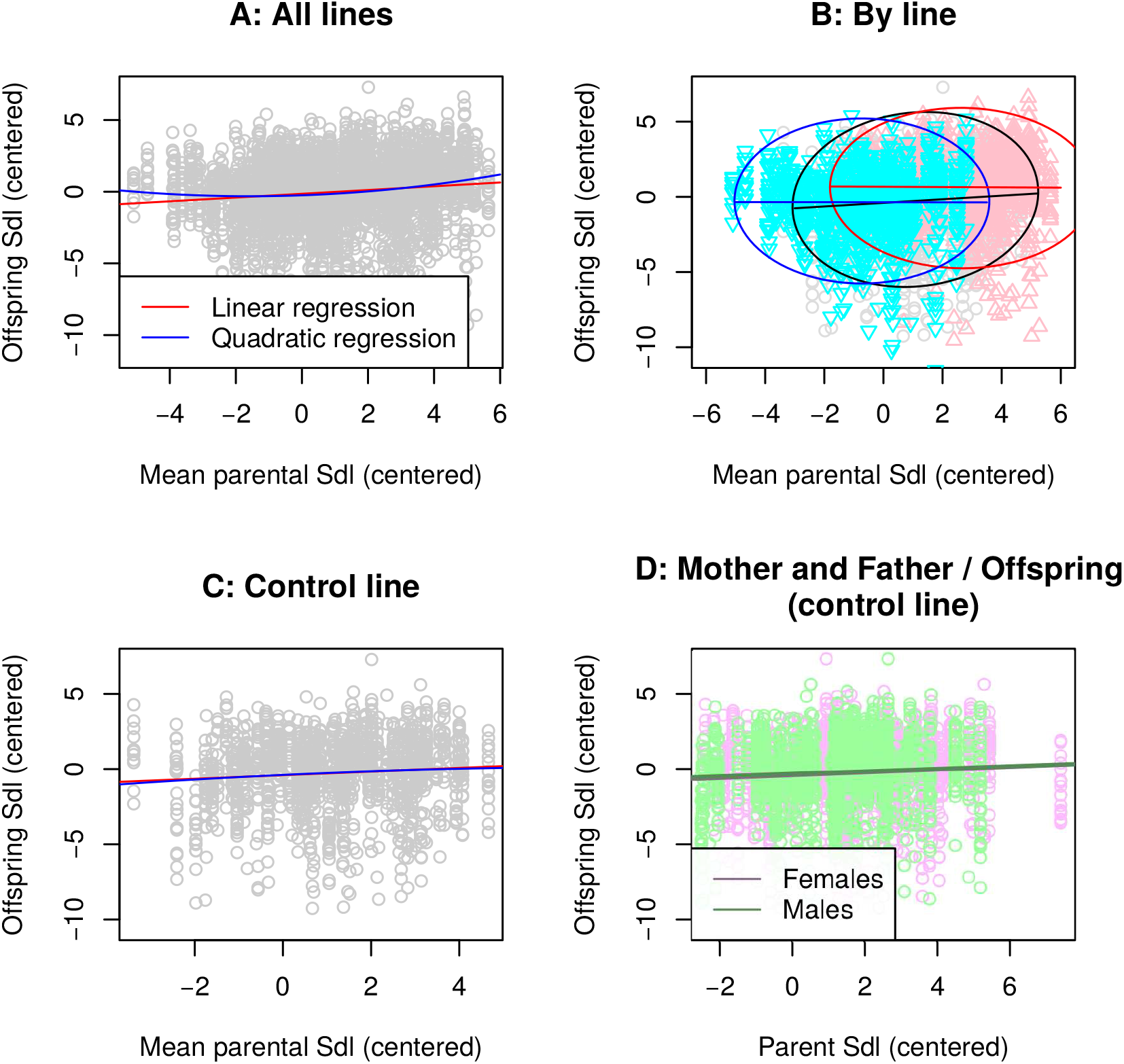

The mid-parent-offspring regression coefficient estimates trait heritability. In addition, the shape of the parent-offspring relationship is indicative of potential deviations from the infinitesimal model assumptions. Non-linear parent-offspring relationships may indicate dominance, epistasis, or genetic asymmetries.

A. Taking all selected lines into account, normalizing by generation phenotypic averages to cancel out generation effects, the parent-offspring relationship appeared to be slightly non-linear (significant quadratic component: *y* = *c* + *h*^2^*x* + *k*_2_*x*, with *h*^2^ = 0.083 ± *s.e.*0.021 being an estimate of heritability (Pr(*h*^2^ = 0) = 6.57 · 10^−5^), the quadratic term being also significant (Pr(*k*_2_ = 0) = 5.81 · 10^−5^).

B. However, considering each line separately, the pattern rather reflected different linear relationships in all three lines. The Large line response to selection shifted the offspring phenotype upwards, while the Small line lack of response set the average offspring at the same level as the Control. Non-linearity in this case was the consequence, rather than the cause, of the asymmetric response.

C. When considering the Control line alone, which had the most statistical power because of the large variance in parental phenotypes, the quadratic term disappeared, supporting the fact that the parent-offspring regression was linear (*h*^2^ ≃ 0.14, Pr(*h*^2^ = 0) = 0.00698, Pr(*k*_2_ = 0) = 0.62)

D. Running mother-offspring and father-offspring regressions independently provided very similar results. Focusing on the control line sub-dataset, the mother-offspring regression lead to *h*^2^ = 0.089±0.031 (s.e.), while the father-offspring regression resulted in *h*^2^ = 0.081±0.032.

