## Supplementary material for "Unidirectional response to bidirectional selection on body size II Quantitative genetics": Data files and analysis scripts: fig-drift.R

```
#!/usr/bin/env Rscript

cache.dir <- "./cache"

n.sim <- 1000
plot.type <- c("all", "quantiles")[2]
plot.overlap <- TRUE
quantile.probs <- c(0.025, 0.975)

library(MASS)
library(parallel)
mc.cores <- min(64, detectCores() -1)

makeTransparent<-function(someColor, alpha=100)
# From StackOverflow: https://stackoverflow.com/questions/8047668/transparent-equivalent-of-given-color
{
  newColor<-col2rgb(someColor)
  apply(newColor, 2, function(curcoldata){rgb(red=curcoldata[1], green=curcoldata[2],
    blue=curcoldata[3],alpha=alpha, maxColorValue=255)})
}

pedsim <- function(G, ped, verbose=FALSE) {
	vNA <- rep(NA, nrow(ped))
	newtab <- data.frame(
		Sdl      = vNA,
		Mat      = vNA,
		Sdl.sire = vNA,
		Mat.sire = vNA,
		Sdl.dam  = vNA,
		Mat.dam  = vNA)
	
	newtab[is.na(ped$sire),c("Sdl.sire", "Mat.sire")] <- mvrnorm(sum(is.na(ped$sire)), mu=c(0,0), Sigma=G)
	newtab[is.na(ped$dam),c("Sdl.dam", "Mat.dam")] <- mvrnorm(sum(is.na(ped$dam)), mu=c(0,0), Sigma=G)
	
	loop.count <- 0
	
	while(sum(is.na(newtab$Sdl) > 0)) {
		loop.count <- loop.count + 1
		if (verbose) {
			cat("Loop ", loop.count, ", ", sum(is.na(newtab$Sdl)), " remaining.\n")
		}
		canbecalc <- which(is.na(newtab$Sdl) & (!is.na(newtab$Sdl.sire)) & (!is.na(newtab$Sdl.dam)))
		stopifnot(length(canbecalc) > 0)
		newtab[canbecalc,c("Sdl","Mat")] <- mvrnorm(length(canbecalc), mu=c(0,0), Sigma=G/2) + c(rowMeans(newtab[canbecalc,c("Sdl.sire","Sdl.dam")]), rowMeans(newtab[canbecalc,c("Mat.sire","Mat.dam")]))
		for (i in canbecalc) { # not the most efficient way
			whichfather <- which(ped$animal[i] == ped$sire)
			whichmother <- which(ped$animal[i] == ped$dam)
			if (length(whichfather) > 0) newtab[whichfather, c("Sdl.sire","Mat.sire")] <- newtab[i,c("Sdl","Mat")]
			if (length(whichmother) > 0) newtab[whichmother, c("Sdl.dam","Mat.dam")] <- newtab[i,c("Sdl","Mat")]	
		}
	}
	return(newtab[,c("Sdl","Mat")])
}

funcdir  <- "../R_functions"

col.line <- c(O="black", A="darkgray", P="blue", G="red")

source(paste(funcdir, "AnimalModel.R", sep="/"))

source("./models.R")

Gtab <- do.call(rbind, mod.all[[1]]$VCV)[,paste0("trait", c("Sdl","Sdl","Mat","Mat"), ":trait", c("Sdl","Mat","Sdl","Mat"), ".animal")]
ped <- mod.args[[1]]$pedigree
gen <- mod.args[[1]]$data$gen
line <- mod.args[[1]]$data$Line

gentrends <- read.table("genBLUPs.txt", header=TRUE)

cache.file <- paste(cache.dir, "fig-drift.rds", sep="/")

ss <- NULL
if (file.exists(cache.file)) 
	ss <- readRDS(cache.file)

if (is.null(ss) || length(ss) < n.sim)
	ss <- mclapply(sample(1:nrow(Gtab), n.sim, replace=(n.sim > nrow(Gtab))), function(i) gentab(pedsim(matrix(Gtab[i,], ncol=2), ped.all), gen=gen, line=line), mc.cores=mc.cores)

saveRDS(ss, file=cache.file)

ylabs <- c(Sdl="Body size (mm), avg breeding value", Mat="Maturity (probit), avg breeding value")

pdf("fig-drift.pdf", width=10, height=if(plot.overlap) 4 else 12)

layout(if (plot.overlap) t(1:2) else cbind(1:3,4:6))

par(mar=c(3, 4, 1, 1), cex=1/par("cex"), lwd=2)
# common scales for all figures
ylim.sdl <- c(-1.25, 1.25)
ylim.mat <- c(-1.25, 1.25)

for (tt in c("Sdl","Mat")) {
	for (ll in c("A", "G", "P")) {
		if (ll == "A" || !plot.overlap) {
			plot(NULL, xlim=c(0,7), xaxt="n", xlab="", ylim=if(tt=="Sdl") ylim.sdl else ylim.mat, ylab=ylabs[tt])
			axis(1, at=0:7, paste0("F", 0:7))
		}
		lines(0:7, gentrends[,paste(tt, ll, sep=".")], type="b", col=col.line[ll], lwd=3)
		if (plot.type=="all") 
			invisible(sapply(ss, function(x) lines(0:7, x[,paste(tt, ll, sep=".")], type="l", col=makeTransparent(col.line[ll], alpha=10))))
		if (plot.type=="quantiles") {
			dd <- sapply(ss, function(x) x[,paste(tt, ll, sep=".")])
			qq <- apply(dd, 1, quantile, prob=quantile.probs)
			invisible(apply(qq, 1, lines, x=0:7, lty=2, col=col.line[ll]))
		}
	}
}

dev.off()


## Estimation of Ne

num.gen.drift <- 7

for (tt in c("Sdl","Mat")) {
	for (ll in c("A", "G", "P")) {
		v.drift <- var(sapply(ss, function(x) x[nrow(x), paste(tt, ll, sep=".")]))
		v.a <- mean(Gtab[,paste0("trait", tt, ":trait", tt, ".animal")])
		cat("Ne estimate for", tt, "line", ll, ":", num.gen.drift*v.a/v.drift, "\n")
	}
}
```
