## Supplementary material for "Unidirectional response to bidirectional selection on body size II Quantitative genetics": Data files and analysis scripts: fig-genetictrends.R

```
#!/usr/bin/env Rscript

funcdir  <- "../R_functions"

col.line <- c(O="black", A="darkgray", P="blue", G="red")

source(paste(funcdir, "AnimalModel.R", sep="/"))

source("./models.R")

quantilelist <- function(ll, probs=c(0.025, 0.50, 0.975)) {
	require(abind)
	arr <- do.call(abind, c(lapply(ll, as.matrix), list(along=3)))
	apply(arr, c(1,2), quantile, probs=probs)
}

# Save the environmental BLUPs in a file 
envtrends <- envMCMC(mod.all[[1]]) 
write.table(quantilelist(envtrends, probs=0.50), file="envBLUPs.txt", quote=FALSE, sep="\t")

# Save the genetic trends
gentrends <- genMCMC(mod.all[[1]], mydata.all$gen, mydata.all$Line)
write.table(quantilelist(gentrends, probs=0.50), file="genBLUPs.txt", quote=FALSE, sep="\t")

grandmean.sdl <- 0 # mean(do.call("c", lapply(mod.all[[1]]$Sol, function(mm) mm[,paste0("traitSdl")])))
grandmean.mat <- 0 # mean(do.call("c", lapply(mod.all[[1]]$Sol, function(mm) mm[,paste0("traitMat")])))

# common scales for all figures
ylim.sdl <- c(-1,1.9) # c(19, 22.1)
ylim.mat <- c(-1,1.2) # c(0.8, 5)

ylim.sdl.env <- c(-2, 3)
ylim.mat.env <- c(-3,3.5)

delta.x <- 0.1

q.envtrends <- quantilelist(envtrends, probs=c(0.025, 0.50, 0.975))
q.gentrends <- quantilelist(gentrends, probs=c(0.025, 0.50, 0.975))

pdf("fig-genetictrends.pdf", width=15, height=4)
layout(t(1:3), widths=c(1, 1, 1.15))

par(mar=c(3, 4, 1, 1), cex=0.85/par("cex"), lwd=2)

plot(NULL, xlim=c(0,7), ylim=ylim.sdl, xlab="", xaxt="n", ylab="Body size (mm)")
axis(1, at=0:7, paste0("F", 0:7))

arrows(x0=1:7, y0=grandmean.sdl+q.gentrends[1,,][paste0("F",1:7),"Sdl.A"], y1=grandmean.sdl+q.gentrends[3,,][paste0("F",1:7),"Sdl.A"], col=col.line["A"], length=0)
arrows(x0=1:7+delta.x, y0=grandmean.sdl+q.gentrends[1,,][paste0("F",1:7),"Sdl.G"], y1=grandmean.sdl+q.gentrends[3,,][paste0("F",1:7),"Sdl.G"], col=col.line["G"], length=0)
arrows(x0=1:7-delta.x, y0=grandmean.sdl+q.gentrends[1,,][paste0("F",1:7),"Sdl.P"], y1=grandmean.sdl+q.gentrends[3,,][paste0("F",1:7),"Sdl.P"], col=col.line["P"], length=0)
arrows(x0=0:1, y0=grandmean.sdl+q.gentrends[1,,][c("F0","F1"),"Sdl.A"], y1=grandmean.sdl+q.gentrends[3,,][c("F0","F1"),"Sdl.A"], col=col.line["O"], length=0)

lines(1:7, grandmean.sdl+q.gentrends[2,,][paste0("F",1:7),"Sdl.A"], type="b", col=col.line["A"])
lines(1:7+delta.x, grandmean.sdl+q.gentrends[2,,][paste0("F",1:7),"Sdl.G"], type="b", col=col.line["G"])
lines(1:7-delta.x, grandmean.sdl+q.gentrends[2,,][paste0("F",1:7),"Sdl.P"], type="b", col=col.line["P"])
lines(0:1, grandmean.sdl+q.gentrends[2,,][c("F0","F1"), "Sdl.A"], type="b", col=col.line["O"])


plot(NULL, xlim=c(0,7), ylim=ylim.mat, xlab="", xaxt="n", ylab="Maturity (probit scale)")
axis(1, at=0:7, paste0("F", 0:7))

arrows(x0=1:7, y0=grandmean.mat+q.gentrends[1,,][paste0("F",1:7),"Mat.A"], y1=grandmean.sdl+q.gentrends[3,,][paste0("F",1:7),"Mat.A"], col=col.line["A"], length=0)
arrows(x0=1:7+delta.x, y0=grandmean.mat+q.gentrends[1,,][paste0("F",1:7),"Mat.G"], y1=grandmean.sdl+q.gentrends[3,,][paste0("F",1:7),"Mat.G"], col=col.line["G"], length=0)
arrows(x0=1:7-delta.x, y0=grandmean.mat+q.gentrends[1,,][paste0("F",1:7),"Mat.P"], y1=grandmean.sdl+q.gentrends[3,,][paste0("F",1:7),"Mat.P"], col=col.line["P"], length=0)
arrows(x0=0:1, y0=grandmean.mat+q.gentrends[1,,][c("F0","F1"),"Mat.A"], y1=grandmean.sdl+q.gentrends[3,,][c("F0","F1"),"Mat.A"], col=col.line["O"], length=0)

lines(1:7, grandmean.mat+q.gentrends[2,,][paste0("F",1:7),"Mat.A"], type="b", col=col.line["A"])
lines(1:7+delta.x, grandmean.mat+q.gentrends[2,,][paste0("F",1:7),"Mat.G"], type="b", col=col.line["G"])
lines(1:7-delta.x, grandmean.mat+q.gentrends[2,,][paste0("F",1:7),"Mat.P"], type="b", col=col.line["P"])
lines(0:1, grandmean.mat+q.gentrends[2,,][c("F0","F1"), "Mat.A"], type="b", col=col.line["O"])


# Environmental trends
x.x <- 0:7
xlim <- c(-0.2, 7.2)
par(mar=c(3, 4, 1, 4), lwd=2)
plot(x.x, grandmean.sdl+q.envtrends[2,,][,"Sdl"], type="b", xaxt="n", xlab="", xlim=xlim, ylim=ylim.sdl.env, ylab="Body size (mm)")
arrows(x0=x.x, y0=grandmean.sdl+q.envtrends[1,,][,"Sdl"], y1=grandmean.sdl+q.envtrends[3,,][,"Sdl"], length=0)
par(new=TRUE)
plot(x.x+delta.x, grandmean.mat+q.envtrends[2,,][,"Mat"], type="b", col="darkgreen", lty=2, axes=FALSE, xlab="", ylab="", xlim=xlim, ylim=ylim.mat.env)
arrows(x0=x.x+delta.x, y0=grandmean.mat+q.envtrends[1,,][,"Mat"], y1=grandmean.mat+q.envtrends[3,,][,"Mat"], col="darkgreen", length=0, lty=2)

axis(1, at=x.x, paste0("F", 0:7))
axis(4, col.ticks="darkgreen", col.axis="darkgreen")
mtext(side=4, line=2,  col="darkgreen", "Maturity (probit scale)", cex=par("cex"))
par(new=FALSE)
legend("topright", col=c("black","darkgreen"), lty=1:2, legend=c("Size", "Maturity"))

dev.off()
```
