## Supplementary material for "Unidirectional response to bidirectional selection on body size II Quantitative genetics": Data files and analysis scripts: fig-gradients.R

```
#!/usr/bin/env Rscript

# A: control, G: large, P: small
line.col <- c(A="darkgray", G="red", P="blue")
line.names <- c(A="Control", G="Large", P="Small")

lines <- setNames(names(line.col), names(line.col))

funcdir  <- "../R_functions"

makeTransparent<-function(someColor, alpha=100)
# From StackOverflow: https://stackoverflow.com/questions/8047668/transparent-equivalent-of-given-color
{
  newColor<-col2rgb(someColor)
  apply(newColor, 2, function(curcoldata){rgb(red=curcoldata[1], green=curcoldata[2],
    blue=curcoldata[3],alpha=alpha, maxColorValue=255)})
}

source(paste(funcdir, "Dataclean.R", sep="/"))

sdl <- cleanSdl()

seldiff.tab <- read.table("tab-seldiff.txt", header=TRUE) # from fig-seldiff.R


pdf("fig-gradients.pdf", width=4, height=4)
par(mar=c(5, 5, 2, 1), lwd=2)

plot(NULL, xlim=c(-3, 3.5), ylim=c(-11, 18), xlab=expression("Selection gradient on Body size (mm"^{-1}*")"), ylab=expression("Selection gradient on Maturity (prob"^{-1}*")"))

legend("topright", col=line.col, lty=1, legend=line.names)
mtext("A", side=3, line=0, adj=-0.2, cex=1.6)

Pmat.all <- var(cbind(sdl$Sdl, sdl$Mat), na.rm=TRUE)
Pmat.algo <- c("all", "linegen")[2]

ssgrad.table <- NULL # gradients are stored to be saved in a file
for (ll in lines) {
	ssgrad <- sapply(1:6, function(gg) {
		filter <- sdl$Generation == paste0("F",gg) & if(gg==1) TRUE else sdl$Line==ll
		Pmat <- if (Pmat.algo == "all") Pmat.all else var(cbind(sdl$Sdl[filter], sdl$Mat[filter]), na.rm=TRUE)
		dff <- seldiff.tab[paste0("F",gg),paste0(c("Sdl","MatTRUE"), ".", ll)]
		setNames(c(solve(Pmat) %*% unlist(dff)), c("Sdl","MatTRUE"))
	})
	ssgrad.table <- cbind(ssgrad.table, t(ssgrad))
	lines(c(0,cumsum(ssgrad["Sdl",])), c(0,cumsum(ssgrad["MatTRUE",])), type="b", col=line.col[ll], lty=1, pch=as.character(0:6))
}

dev.off()


# Table: gradients per line
grad.table <- ssgrad.table # This can be easily changed among tgrad, rgrad, and ssgrad
rownames(grad.table) <- paste0("F",1:6)
colnames(grad.table) <- paste0(c("Sdl","Mat"), ".", rep(lines, each=2))

write.table(round(grad.table, digits=4), file="tab-gradients.txt", sep="\t", col.names=TRUE, quote=FALSE)
```
