## Supplementary material for "Unidirectional response to bidirectional selection on body size II Quantitative genetics": Data files and analysis scripts: fig-inbreeding.R

```
#!/usr/bin/env Rscript

line.col <- c(A="gray", G="red", P="blue")
line.names <- c(A="Control", G="Large", P="Small")
line.pch <- c(A=1, G=2, P=6)

funcdir  <- "../R_functions"

maxgen <- 7

source(paste(funcdir, "Dataclean.R", sep="/"))
source(paste(funcdir, "Inbreeding.R", sep="/"))

degener <- function(ff) as.numeric(unlist(sapply(strsplit(ff, ""), "[", 2)))

sdl <- cleanSdl()
fam <- cleanFam(sdl)

sdl$inb <- Inbreeding(sdl)
fam$inb  <- sdl$inb[match(rownames(fam), as.character(sdl$Family))]

genMeanInb <- tapply(sdl$inb,  INDEX=sdl$Generation, FUN=mean, na.rm=TRUE)
linMeanInb <- tapply(sdl$inb, INDEX=list(sdl$Generation, sdl$Line), FUN=mean, na.rm=TRUE)
#linMeanInb <- linMeanInb[paste0("F", 1:7),]
fam$LinInb <- linMeanInb[cbind(fam$Gen, fam$Line)]

relinb <- fam$inb - fam$LinInb
relsdl <- fam$MeanSdl - fam$LinSdl
relmat <- fam$MeanMat - fam$LinMat

source("models.R")

mod.inb <- mod.all[[grep(names(mod.all), pattern="i$")]]

pr.inb.Sdl <- do.call(rbind, mod.inb$Sol)[,"traitSdl:inb"]
pr.inb.Mat <- do.call(rbind, mod.inb$Sol)[,"traitMat:inb"]


pdf("fig-inbreeding.pdf", width=8, height=12)

	layout(rbind(c(1, 1), c(2, 3), c(4, 5)))
	par(mar=c(5, 5, 0.1, 1), lwd=2, cex=0.8/par("cex"))
	
	bxpxx <- 1:(8*4)
	bxpsplFG <- split(fam$inb, fam$Gen)
	bxpsplFGL <- split(fam$inb, list(fam$Gen, fam$Line))
	boxplot(bxpsplFG[paste0("F", 0:1)], xlim=range(bxpxx), at=c(2.5, 6.5), ylim=range(fam$inb), xaxt="n", xlab="Generation", ylab="Family Inbreeding coefficient")
	boxplot(bxpsplFGL[paste0("F", 2:maxgen,".A")], at=4*(2:maxgen)+1, add=TRUE, col="gray", xaxt="n", yaxt="n")
	boxplot(bxpsplFGL[paste0("F", 2:maxgen,".G")], at=4*(2:maxgen)+2, add=TRUE, col="red", xaxt="n", yaxt="n")
	boxplot(bxpsplFGL[paste0("F", 2:maxgen,".P")], at=4*(2:maxgen)+3, add=TRUE, col="blue", xaxt="n", yaxt="n")
	axis(1, at=4*(0:maxgen)+2, paste0("F", 0:maxgen))
	
	
	plot(relinb, relsdl, xlab="Relative Inbreeding coefficient", ylab="Relative Body size (mm)", col=line.col[as.character(fam$Line)], pch=line.pch[as.character(fam$Line)])

	plot(relinb, relmat, xlab="Relative Inbreeding coefficient", ylab="Relative Maturity", col=line.col[as.character(fam$Line)], pch=line.pch[as.character(fam$Line)])
	
	plot(density(pr.inb.Sdl), xlab="Inbreeding effect (Sdl)", ylab="Posterior density", main="")
	abline(v=quantile(pr.inb.Sdl, c(0.025, 0.5, 0.975)), lty=c(3,1,3), col="red")
	
	plot(density(pr.inb.Mat), xlab="Inbreeding effect (Mat)", ylab="Posterior density", main="")
	abline(v=quantile(pr.inb.Mat, c(0.025, 0.5, 0.975)), lty=c(3,1,3), col="red")

dev.off()
```
