## Supplementary material for "Unidirectional response to bidirectional selection on body size II Quantitative genetics": Data files and analysis scripts: fig-matrices.R

```
#!/usr/bin/env Rscript

library(ellipse)

source("./models.R")

makeTransparent<-function(someColor, alpha=100)
# From StackOverflow: https://stackoverflow.com/questions/8047668/transparent-equivalent-of-given-color
{
  newColor<-col2rgb(someColor)
  apply(newColor, 2, function(curcoldata){rgb(red=curcoldata[1], green=curcoldata[2],
    blue=curcoldata[3],alpha=alpha, maxColorValue=255)})
}


mod <- mod.all[[1]]
xlong <- do.call(rbind, mod$VCV)[1:100,] # more lines just makes overcrowded figures

matrix.col <- c(animal="orange", aqua="chartreuse3", units="gray")

pdf("fig-matrices.pdf", width=5, height=5)
	par(mar=c(4,4,1,1))
	plot(NULL, xlim=c(-2, 2), ylim=c(-1.2,1.2), xlab="Sdl (mm)", ylab="Maturity (probit)")
	
	for (vcomp in rev(names(matrix.col))) {
		for (i in 1:nrow(xlong)) {
			mm <- matrix(xlong[i,c(paste0("traitSdl:traitSdl.", vcomp), paste0("traitMat:traitSdl.", vcomp), paste0("traitSdl:traitMat.", vcomp), paste0("traitMat:traitMat.", vcomp))], ncol=2)
			lines(ellipse(mm, t=1), col=makeTransparent(matrix.col[vcomp], 30))
		}
	}
	for (vcomp in rev(names(matrix.col))) {
		mm <- matrix(colMeans(xlong)[c(paste0("traitSdl:traitSdl.", vcomp), paste0("traitMat:traitSdl.", vcomp), paste0("traitSdl:traitMat.", vcomp), paste0("traitMat:traitMat.", vcomp))], ncol=2)
		lines(ellipse(mm, t=1), col=matrix.col[vcomp], lwd=3)
	}
	legend("topleft", lty=1, col=matrix.col, legend=c("Genetic", "Aquarium", "Residual"))

dev.off()
```
