## Supplementary material for "Unidirectional response to bidirectional selection on body size II Quantitative genetics": Data files and analysis scripts: fig-paroff.R

```
#!/usr/bin/env Rscript

line.col <- c(A="gray", G="red", P="blue")
line.names <- c(A="Control", G="Large", P="Small")
line.pch <- c(A=1, G=2, P=6)

funcdir  <- "../R_functions"

source(paste(funcdir, "Dataclean.R", sep="/"))

degener <- function(ff) as.numeric(unlist(sapply(strsplit(ff, ""), "[", 2)))

sdl <- cleanSdl()

meanSdlGen <- tapply(sdl$Sdl, sdl$Gen, mean, na.rm=TRUE)

sdl$SdlCor <- sdl$Sdl - meanSdlGen[as.character(sdl$Gen)]
sdl$FatherSdl <- ifelse(is.na(sdl$Father), NA, sdl$Sdl[match(sdl$Father, sdl$ID)])
sdl$MotherSdl <- ifelse(is.na(sdl$Mother), NA, sdl$Sdl[match(sdl$Mother, sdl$ID)])
sdl$MeanParentSdl <- 0.5*sdl$FatherSdl + 0.5*sdl$MotherSdl - meanSdlGen[paste0("F", degener(as.character(sdl$Gen))+1)]
# No need to compute FatherMat and MotherMat: they are all 1 by definition

# Dataset split by line. The rule is not straightforward, as 
# generations F0 and F1 should be considered as "controls"
sdlC <- sdl[sdl$Line=="A" | sdl$Gen %in% c("F0","F1"),]
sdlG <- sdl[sdl$Line=="G" & !(sdl$Gen %in% c("F0","F1")),]
sdlP <- sdl[sdl$Line=="P" & !(sdl$Gen %in% c("F0","F1")),]

meanSdlCGen <- tapply(sdlC$Sdl, sdlC$Gen, mean, na.rm=TRUE)
sdlC$sdlCor <- sdlC$Sdl - meanSdlCGen[as.character(sdlC$Gen)]

# Full dataset, centered on generation mean
reg1 <- lm(SdlCor ~ MeanParentSdl, data=sdl)
reg2 <-  lm(SdlCor ~ MeanParentSdl + I(MeanParentSdl^2), data=sdl)

mpartest <- data.frame(MeanParentSdl=seq(-6, 6, by=0.1))

pdf("fig-paroff.pdf", width=6, height=6)
	layout(rbind(1:2, 3:4))
	par(cex=0.8)

	plot(sdl$MeanParentSdl, sdl$SdlCor, xlab="Mean parental Sdl (centered)", ylab="Offspring Sdl (centered)", main="A: All lines", col="gray")
	lines(mpartest$MeanParentSdl, predict(reg1, mpartest), col="red")
	lines(mpartest$MeanParentSdl, predict(reg2, mpartest), col="blue")
	
	legend("bottomleft", lty=1, bg=rgb(255,255,255,alpha=230,max=255), col=c("red","blue"), legend=c("Linear regression", "Quadratic regression"))
	
	library(ellipse)
	
	plot(sdlC$MeanParentSdl, sdlC$SdlCor, xlab="Mean parental Sdl (centered)", ylab="Offspring Sdl (centered)", col="lightgray", xlim=c(-6,6), main="B: By line")
	points(sdlG$MeanParentSdl, sdlG$SdlCor, col="pink", pch=line.pch["G"])
	points(sdlP$MeanParentSdl, sdlP$SdlCor, col="cyan", pch=line.pch["P"])
		
	ellipsC <- ellipse(var(cbind(sdlC$MeanParentSdl, sdlC$SdlCor), na.rm=TRUE), centre=colMeans(cbind(sdlC$MeanParentSdl, sdlC$SdlCor), na.rm=TRUE))
	ellipsG <- ellipse(var(cbind(sdlG$MeanParentSdl, sdlG$SdlCor), na.rm=TRUE), centre=colMeans(cbind(sdlG$MeanParentSdl, sdlG$SdlCor), na.rm=TRUE))
	ellipsP <- ellipse(var(cbind(sdlP$MeanParentSdl, sdlP$SdlCor), na.rm=TRUE), centre=colMeans(cbind(sdlP$MeanParentSdl, sdlP$SdlCor), na.rm=TRUE))
	
	lines(ellipsC, col="black")
	lines(ellipsG, col="red")
	lines(ellipsP, col="blue")
	
	rangeC <- which(mpartest$MeanParentSdl > min(ellipsC[,1]) & mpartest$MeanParentSdl <= max(ellipsC[,1]))
	rangeG <- which(mpartest$MeanParentSdl > min(ellipsG[,1]) & mpartest$MeanParentSdl <= max(ellipsG[,1]))
	rangeP <- which(mpartest$MeanParentSdl > min(ellipsP[,1]) & mpartest$MeanParentSdl <= max(ellipsP[,1]))
	
	lines(mpartest$MeanParentSdl[rangeC], predict(lm(SdlCor ~ MeanParentSdl, data=sdlC), mpartest)[rangeC], col="black")
	lines(mpartest$MeanParentSdl[rangeG], predict(lm(SdlCor ~ MeanParentSdl, data=sdlG), mpartest)[rangeG], col="red")
	lines(mpartest$MeanParentSdl[rangeP], predict(lm(SdlCor ~ MeanParentSdl, data=sdlP), mpartest)[rangeP], col="blue")
	
	
	# Control line only
	reg1C <- lm(SdlCor ~ MeanParentSdl, data=sdlC)
	reg2C <-  lm(SdlCor ~ MeanParentSdl + I(MeanParentSdl^2), data=sdlC)
	
	plot(sdlC$MeanParentSdl, sdlC$SdlCor, xlab="Mean parental Sdl (centered)", ylab="Offspring Sdl (centered)", main="C: Control line", col="gray")
	lines(mpartest$MeanParentSdl, predict(reg1C, mpartest), col="red")
	lines(mpartest$MeanParentSdl, predict(reg2C, mpartest), col="blue")
	
	# Mother/Offspring and Father/Offspring
	moSdl <- sdlC$MotherSdl- meanSdlCGen[paste0("F", degener(as.character(sdlC$Gen))+1)]
	faSdl <- sdlC$FatherSdl- meanSdlCGen[paste0("F", degener(as.character(sdlC$Gen))+1)]
	plot(moSdl, sdlC$SdlCor, xlab="Parent Sdl (centered)", ylab="Offspring Sdl (centered)", main="D: Mother and Father / Offspring \n(control line)", type="p", col="plum1")
	points(faSdl, sdlC$SdlCor, col="palegreen1")
	abline(lm(sdlC$SdlCor ~ moSdl), col="plum4", lty=1, lwd=2)
	abline(lm(sdlC$SdlCor ~ faSdl), col="palegreen4", lty=1, lwd=2)
	
	

	legend("bottomleft", lty=1, col=c("plum4", "palegreen4"), legend=c("Females", "Males"), bg=rgb(255,255,255,alpha=230,max=255))
	
	
	
dev.off()
```
