## Supplementary material for "Unidirectional response to bidirectional selection on body size II Quantitative genetics": Data files and analysis scripts: fig-partialdata.R

```
#!/usr/bin/env Rscript

source("./models.R")

best.args <- mod.args[[names(mod.all)[1]]]
mod.best <- mod.all[[1]]

gensel <- list('F4-7'=c("F0","F1","F2","F3"), 'F0-3'=c("F4","F5","F6","F7"))

# Without specific generations
gofgen <- sapply(names(gensel), function(gi) {
	gen <- gensel[[gi]]
	tokeep <- !(best.args$data$gen %in% gen)
	
	aa <- best.args
	aa$data <-best.args$data[tokeep,]
	loadORrun(paste0(names(mod.all)[1], "-", gi), aa)
}, simplify=FALSE, USE.NAMES=TRUE)

genline <- list(noSmall="P",noLarge="G",noControl="A")

# Without a specific selection line
gofline <- sapply(names(genline), function(li) {
	line <- genline[[li]]
	tokeep <- best.args$data$gen %in% c("F0","F1") | best.args$data$Line != line
	
	aa <- best.args
	aa$data <- best.args$data[tokeep,]
	loadORrun(paste0(names(mod.all)[1], "-", li), aa)
}, simplify=FALSE, USE.NAMES=TRUE)

sensitiv.unit <- function(ref, totest, component, ylab=component, stars=TRUE, ...) {
	boxplot(lapply(c(list(Ref=ref), totest), function(mm) do.call("c", lapply(mm$VCV, function(mmm) mmm[,component]))), ylab=ylab, range=0, las=2, ...)
	if (stars)
		text(x=2:length(totest), y=0, sapply(totest, function(mm) { diffs <- do.call("c", lapply(mm$VCV, function(mmm) mmm[,component])) - do.call("c", lapply(ref$VCV, function(mmm) mmm[,component])); if (mean(diffs < 0) < 0.025 || mean(diffs>0) < 0.025) "*" else ""}), cex=2)
	
}

sensitiv.all <- function(ref, totest, ...) {
	layout(cbind(1:2, 3:4, 5:6))
	par(mar=c(6, 5, 0.5, 0.5))
	sensitiv.unit(ref, totest, "traitSdl:traitSdl.animal", ylab=expression(V[G](Sdl)), ylim=c(0, 3.5), ...)
	sensitiv.unit(ref, totest, "traitMat:traitMat.animal", ylab=expression(V[G](Mat)), ylim=c(0, 2.5), ...)
	sensitiv.unit(ref, totest, "traitSdl:traitMat.animal", ylab=expression(C[G](Sdl,Mat)), ylim=c(0, 2.5), ...)
	sensitiv.unit(ref, totest, "traitSdl:traitSdl.units", ylab=expression(V[E](Sdl)), ylim=c(0, 4.5), ...)
	sensitiv.unit(ref, totest, "traitSdl:traitMat.units", ylab=expression(C[E](Sdl,Mat)), ylim=c(0, 2), ...)
}

pdf("fig-partialdata.pdf", width=10, height=6)
sensitiv.all(mod.best, c(gofgen,gofline))
dev.off()
```
