## Supplementary material for "Unidirectional response to bidirectional selection on body size II Quantitative genetics": Data files and analysis scripts: fig-prediction.R

```
#!/usr/bin/env Rscript

library(QGglmm)
library(parallel)
library(MASS)

mat.latent.scale <- FALSE
env.correct <- FALSE
gen.trends <- TRUE

max.posteriors <- Inf

mc.cores <- detectCores()-1
line.col <- c(A="gray", G="red", P="blue")

degener <- function(gener) as.numeric(unlist(sapply(strsplit(as.character(gener),split="F"), "[", 2)))
makeOldID <- function(fam, sex, line) 
    paste0(sub("\\.", "", as.character(fam)), as.character(sex), as.character(round(runif(length(fam), 1e6, 1e7-1))), as.character(line))
     
funcdir  <- "../R_functions"
cachedir <- "./cache"

source(paste(funcdir, "Dataclean.R", sep="/"))
source(paste(funcdir, "Inbreeding.R", sep="/"))
source(paste(funcdir, "AnimalModel.R", sep="/"))

source("models.R")
grads <- read.table("tab-gradients.txt", header=TRUE)

inv.links <- function(x) rbind(QGlink.funcs("Gaussian")$inv.link(x[1,]), QGlink.funcs("binom1.probit")$inv.link(x[2,]))

makeTransparent<-function(someColor, alpha=100)
# From StackOverflow: https://stackoverflow.com/questions/8047668/transparent-equivalent-of-given-color
{
  newColor<-col2rgb(someColor)
  apply(newColor, 2, function(curcoldata){rgb(red=curcoldata[1], green=curcoldata[2],
    blue=curcoldata[3],alpha=alpha, maxColorValue=255)})
}

fit.latent <- function(mean.sdl, beta.sdl, mean.mat, beta.mat) function(l) {
		1-mean.sdl*beta.sdl-pnorm(mean.mat)*beta.mat + beta.sdl*l[1,] + beta.mat*pnorm(l[2,])
	}
d.fit.latent <- function(mean.sdl, beta.sdl, mean.mat, beta.mat) function(l) {
		rbind(beta.sdl, beta.mat*dnorm(l[2,]))
	}

response.pred <- function(mu, G, P, beta, mu.data=NULL, Ne=Inf) {
	if (is.null(mu.data))
		mu.data <- QGmvmean(mu ,vcov=P, link.inv=inv.links)
	
	fit.args <- list(mean.sdl=mu.data[1], beta.sdl=beta[1], mean.mat=mu.data[2], beta.mat=beta[2])
	fit.fun <- do.call(fit.latent, fit.args)
	d.fit.fun <- do.call(d.fit.latent, fit.args)
	
	pred <- QGmvpred(mu=mu, vcv.G=G, vcv.P=P, fit.func=fit.fun, d.fit.func=d.fit.fun, verbose=FALSE)
	newgen <- mvrnorm(1, mu=mu+pred$lat.resp, Sigma=G/Ne)
	
	list(data.scale=QGmvmean(newgen, vcov=P, link.inv=inv.links),
		latent.scale=newgen)
}

response.pred.ts <- function(mu, G, P, betas, Ne=Inf) {
	ans <- t(c(mu, QGmvmean(mu, vcov=P, link.inv=inv.links)))
	for (bb in 1:nrow(betas)) {
		rr <- response.pred(mu=ans[bb,1:2], G, P, betas[bb,], mu.data=ans[bb,3:4], Ne=Ne)
		ans <- rbind(ans, c(rr$latent.scale, rr$data.scale))
	}
	colnames(ans) <- c("Sdl.latent", "Mat.latent", "Sdl.data", "Mat.data")
	rownames(ans) <- paste0("F", 1:(nrow(betas)+1))
	ans
}

lines.response <- function(pred.matrix, trait="Sdl", scale="data", ...) {
	lines(0:(nrow(pred.matrix)-1), pred.matrix[,paste0(trait, ".", scale)], ...)
}

to.data.scale <- function(latent.Sdl, latent.Mat, P) {
	sapply(1:length(latent.Sdl), function(i) QGmvmean(mu=c(latent.Sdl[i], latent.Mat[i]), vcov=P, link.inv=inv.links))
}

mod <- as.MCMCglmm(mod.all[[grep(names(mod.all), pattern="c$")]])

tt <- c("Sdl","Mat")
mu <- colMeans(mod[["Sol"]][ , paste0("trait", tt)])
all.G <- mod[["VCV"]][, paste0("trait", rep(tt, each=2), ":trait", rep(tt,2), ".animal")]
all.R <- mod[["VCV"]][, paste0("trait", rep(tt, each=2), ":trait", rep(tt,2), ".units")]
all.A <- mod[["VCV"]][, paste0("trait", rep(tt, each=2), ":trait", rep(tt,2), ".aqua")]
G  <- matrix(colMeans(all.G), ncol=2)
R  <- matrix(colMeans(all.R), ncol=2)
A  <- matrix(colMeans(all.A), ncol=2)
P <- G + R + A

envtrends <- read.table("envBLUPs.txt", header=TRUE)
gentrends <- read.table("genBLUPs.txt", header=TRUE)

# 3 panels

Ne.G <- Ne.P <- 25 # could be Inf

pdf("fig-prediction.pdf", width=12, height=4)
layout(t(1:3))

	par(mar=c(5,4,2,1), cex=par("cex")/0.8)

	mu0 <- sapply(tt, function(ttt) mu[paste0("trait", ttt)] + gentrends["F1", paste0(ttt, ".A")])
	
	mean.Sdl.F1 <- mean(sdl$Sdl[sdl$Generation=="F1"], na.rm=TRUE)
	mean.Mat.F1 <- mean(sdl$Mat[sdl$Generation=="F1"], na.rm=TRUE)

	allresp.G <- mclapply(1:min(max.posteriors,nrow(all.G)),function(i) 
		response.pred.ts(mu0, G=matrix(all.G[i,], ncol=2), P=matrix(all.G[i,], ncol=2)+matrix(all.R[i,], ncol=2)+diag(all.A[i,],2), beta=cbind(grads$Sdl.G, grads$Mat.G), Ne=Ne.G), mc.cores=mc.cores)
	allresp.P <- mclapply(1:min(max.posteriors,nrow(all.G)),function(i) 
		response.pred.ts(mu0, G=matrix(all.G[i,], ncol=2), P=matrix(all.G[i,], ncol=2)+matrix(all.R[i,], ncol=2)+diag(all.A[i,],2), beta=cbind(grads$Sdl.P, grads$Mat.P), Ne=Ne.P), mc.cores=mc.cores)

	############### Panel 1: Sdl
	
	mean.Sdl.G <- c(F1=mean.Sdl.F1, c(by(sdl[sdl$Line=="G","Sdl"], sdl[sdl$Line=="G","Generation"], mean, na.rm=TRUE))[paste0("F", 2:7)])- if(env.correct) envtrends[paste0("F",1:7),"Sdl"] else 0
	mean.Sdl.P <- c(F1=mean.Sdl.F1, c(by(sdl[sdl$Line=="P","Sdl"], sdl[sdl$Line=="P","Generation"], mean, na.rm=TRUE))[paste0("F", 2:7)])- if(env.correct) envtrends[paste0("F",1:7),"Sdl"] else 0
	
	plot(NULL, xlim=c(0,6), ylim=c(18, 24), xaxt="n", xlab="", ylab="Body size (mm)")
	axis(1, at=0:6, paste("F",1:7))

	Sdl.data.G <- sapply(allresp.G, function(x) x[,"Sdl.data"])
	Sdl.data.P <- sapply(allresp.P, function(x) x[,"Sdl.data"])
		
	polygon(x=c(0:6, 6:0), y=c(apply(Sdl.data.G, 1, quantile, probs=0.975), rev(apply(Sdl.data.G, 1, quantile, probs=0.025))), border=NA, col=makeTransparent(line.col["G"], 100))
	polygon(x=c(0:6, 6:0), y=c(apply(Sdl.data.P, 1, quantile, probs=0.975), rev(apply(Sdl.data.P, 1, quantile, probs=0.025))), border=NA, col=makeTransparent(line.col["P"], 100))
	lines.response(response.pred.ts(mu0, G, P, beta=cbind(grads$Sdl.G, grads$Mat.G)), lwd=3, col=line.col["G"])
	lines.response(response.pred.ts(mu0, G, P, beta=cbind(grads$Sdl.P, grads$Mat.P)), lwd=3, col=line.col["P"])

	points(0:6, mean.Sdl.G, col=line.col["G"], type="b", pch=2)
	points(0:6, mean.Sdl.P, col=line.col["P"], type="b", pch=6)

	if (gen.trends) {
		gen.Sdl.G <- mu0["Sdl.traitSdl"]+gentrends[paste0("F",1:7), "Sdl.G"]
		gen.Sdl.P <- mu0["Sdl.traitSdl"]+gentrends[paste0("F",1:7), "Sdl.P"]
		
		points(0:6, gen.Sdl.G, bg=line.col["G"], type="b", pch=24, lty=2)
		points(0:6, gen.Sdl.P, bg=line.col["P"], type="b", pch=25, lty=2)
	}
	mtext("A", side=3, line=0, adj=-0.2, cex=1.6)

	########### Panel 2: Mat 
	mean.Mat.G <- c(F1=mean.Mat.F1, c(by(sdl[sdl$Line=="G","Mat"], sdl[sdl$Line=="G","Generation"], mean, na.rm=TRUE))[paste0("F", 2:7)])
	mean.Mat.P <- c(F1=mean.Mat.F1, c(by(sdl[sdl$Line=="P","Mat"], sdl[sdl$Line=="P","Generation"], mean, na.rm=TRUE))[paste0("F", 2:7)])

		# this is probably less correct than the rest (a reversed QGmvmean would be necessary?)
	latent.Mat.G.env <- qnorm(mean.Mat.G) - envtrends[paste0("F",1:7),"Mat"]
	latent.Mat.P.env <- qnorm(mean.Mat.P) - envtrends[paste0("F",1:7),"Mat"]

	plot(NULL, xlim=c(0,6), ylim=if(mat.latent.scale) c(-1,3) else c(0.6, 1), xaxt="n", xlab="", ylab="Maturity")
	axis(1, at=0:6, paste("F",1:7))

	Mat.data.G <- sapply(allresp.G, function(x) x[,if(mat.latent.scale) "Mat.latent" else "Mat.data"])
	Mat.data.P <- sapply(allresp.P, function(x) x[,if(mat.latent.scale) "Mat.latent" else "Mat.data"])

	polygon(x=c(0:6, 6:0), y=c(apply(Mat.data.G, 1, quantile, probs=0.975), rev(apply(Mat.data.G, 1, quantile, probs=0.025))), border=NA, col=makeTransparent(line.col["G"], 100))
	polygon(x=c(0:6, 6:0), y=c(apply(Mat.data.P, 1, quantile, probs=0.975), rev(apply(Mat.data.P, 1, quantile, probs=0.025))), border=NA, col=makeTransparent(line.col["P"], 100))
	lines.response(response.pred.ts(mu0, G, P, beta=cbind(grads$Sdl.G, grads$Mat.G)), trait="Mat", lwd=3, col=line.col["G"], scale=if(mat.latent.scale) "latent" else "data")
	lines.response(response.pred.ts(mu0, G, P, beta=cbind(grads$Sdl.P, grads$Mat.P)), trait="Mat", lwd=3, col=line.col["P"], scale=if(mat.latent.scale) "latent" else "data")

	if(mat.latent.scale) {
		if (env.correct) {
			points(0:6, latent.Mat.G.env, col=line.col["G"], type="b", pch=2)
			points(0:6, latent.Mat.P.env, col=line.col["P"], type="b", pch=6)			
		} else {
			# No transformation from data to latent scale in QGglmm?
			points(0:6, qnorm(mean.Mat.G.env), col=line.col["G"], type="b", pch=2)
			points(0:6, qnorm(mean.Mat.P.env), col=line.col["P"], type="b", pch=6)	
		}
	} else { # Data scale
		if (env.correct) {
			points(0:6, to.data.scale(mean.Sdl.G, latent.Mat.G.env, P)[2,], type="b", pch=2)
			points(0:6, to.data.scale(mean.Sdl.P, latent.Mat.P.env, P)[2,], col=line.col["P"], type="b", pch=6)
		} else {
			# phenotypic raw data
			points(0:6, mean.Mat.G, col=line.col["G"], type="b", pch=2)
			points(0:6, mean.Mat.P, col=line.col["P"], type="b", pch=6)			
		}
	}
	
	if (gen.trends) {
		gen.Mat.G <- mu0["Mat.traitMat"]+gentrends[paste0("F",1:7), "Mat.G"]		
		gen.Mat.P <- mu0["Mat.traitMat"]+gentrends[paste0("F",1:7), "Mat.P"]
		if (mat.latent.scale) {
			points(0:6, gen.Mat.G, bg=line.col["G"], type="b", pch=24, lty=2)
			points(0:6, gen.Mat.P, bg=line.col["P"], type="b", pch=25, lty=2)
		} else {
			points(0:6, to.data.scale(gen.Sdl.G, gen.Mat.G, P)[2,], bg=line.col["G"], type="b", pch=24, lty=2)
			points(0:6, to.data.scale(gen.Sdl.P, gen.Mat.P, P)[2,], bg=line.col["P"], type="b", pch=25, lty=2)			
			
		}
		
		
	}
	legend("bottomleft", lty=c(1,2,3), pch=c(NA, 2, 24), lwd=c(3, 1, 1), col=c("darkgray","darkgray","black"), pt.bg="darkgray", legend=c("Prediction", "Phenotype", "Breeding values"), bg="white", bty='n')
	mtext("B", side=3, line=0, adj=-0.2, cex=1.6)

	########## Panel 3: bivariate
	library(ellipse)
	library(shape)
	
	mgrad.G <- colMeans(grads)[c("Sdl.G","Mat.G")]
	mgrad.P <- colMeans(grads)[c("Sdl.P","Mat.P")]
	
	fit.args.G <- list(mean.sdl=mu0[1], beta.sdl=mgrad.G[1], mean.mat=mu0[2], beta.mat=mgrad.G[2])
	fit.args.P <- list(mean.sdl=mu0[1], beta.sdl=mgrad.P[1], mean.mat=mu0[2], beta.mat=mgrad.P[2])
	
	pred.G <- QGmvpred(mu=mu0, vcv.G=G, vcv.P=P, fit.func=do.call(fit.latent, fit.args.G), d.fit.func=do.call(d.fit.latent, fit.args.G), verbose=FALSE)
	pred.P <- QGmvpred(mu=mu0, vcv.G=G, vcv.P=P, fit.func=do.call(fit.latent, fit.args.P), d.fit.func=do.call(d.fit.latent, fit.args.P), verbose=FALSE)
	
	resp.G <- c((gen.Sdl.G[7]-gen.Sdl.G[1])/6, (gen.Mat.G[7]-gen.Mat.G[1])/6)
	resp.P <- c((gen.Sdl.P[7]-gen.Sdl.P[1])/6, (gen.Mat.P[7]-gen.Mat.P[1])/6)
	
	# Gradients for maturity on the latent scale: 2 methods
	lat.mgrad.G <- c(mgrad.G[1], mgrad.G[2]*dnorm(mu0[2]))
	lat.mgrad.P <- c(mgrad.P[1], mgrad.P[2]*dnorm(mu0[2]))
	
	lat.mgrad.G  <- pred.G$lat.grad
	lat.mgrad.P <- pred.P$lat.grad
	
	grad.reduce.fact <- 0.1
	G.reduce.fact <- 0.5
	
	plot(NULL, xlab="Body size (mm)", ylab="Maturity (probit)", xlim=c(-0.8,0.8), ylim=c(-0.6,0.6))
	
	# G matrix
	lines(ellipse(G, t=G.reduce.fact), lwd=3, col="orange")
	# gradients
	arr.grad.G.args <- list(x0=0, y0=0, x1=grad.reduce.fact*P[1,1]*lat.mgrad.G[1], y1=grad.reduce.fact*P[2,2]*lat.mgrad.G[2], col=line.col["G"], lwd=2, lty=3)
	arr.grad.P.args <- list(x0=0, y0=0, x1=grad.reduce.fact*P[1,1]*lat.mgrad.P[1], y1=grad.reduce.fact*P[2,2]*lat.mgrad.P[2], col=line.col["P"], lwd=2, lty=3)
		# when calling Arrows with lty != 1, the arrow head is ugly. Two calls are necessary to workaround the bug
	do.call(Arrows, c(arr.grad.G.args, list(arr.width=0, arr.length=0)))
	do.call(Arrows, c(arr.grad.G.args, list(lcol="transparent", arr.col=line.col["G"], arr.length=0.6)))
	do.call(Arrows, c(arr.grad.P.args, list(arr.width=0, arr.length=0)))
	do.call(Arrows, c(arr.grad.P.args, list(lcol="transparent", arr.col=line.col["P"], arr.length=0.6)))

	# predicted response
	Arrows(x0=0, y0=0, x1=pred.G$lat.resp[1], y1=pred.G$lat.resp[2], col=line.col["G"], lwd=2, lty=1)
	Arrows(x0=0, y0=0, x1=pred.P$lat.resp[1], y1=pred.P$lat.resp[2], col=line.col["P"], lwd=2, lty=1)
	
	# genetic response
	arr.resp.G.args <- list(x0=0, y0=0, x1=resp.G[1], y1=resp.G[2], col=line.col["G"], lwd=2, lty=2)
	arr.resp.P.args <- list(x0=0, y0=0, x1=resp.P[1], y1=resp.P[2], col=line.col["P"], lwd=2, lty=2)

	do.call(Arrows, c(arr.resp.G.args, list(arr.width=0, arr.length=0)))
	do.call(Arrows, c(arr.resp.G.args, list(lcol="transparent", arr.col=line.col["G"], arr.length=0.6)))
	do.call(Arrows, c(arr.resp.P.args, list(arr.width=0, arr.length=0)))
	do.call(Arrows, c(arr.resp.P.args, list(lcol="transparent", arr.col=line.col["P"], arr.length=0.6)))
	
	legend("bottomright", lty=c(1,3,1,2), lwd=c(3,2,2,2), col=c("orange",rep("black",3)), legend=c("G matrix (x 0.5)", "Gradients (x 0.1)", "Prediction", "Response"), bty='n')
	mtext("C", side=3, line=0, adj=-0.2, cex=1.6)
	
dev.off()
```
