## Supplementary material for "Unidirectional response to bidirectional selection on body size II Quantitative genetics": Data files and analysis scripts: fig-realized.R

```
#!/usr/bin/env Rscript

line.col <- c(A="darkgray", G="red", P="blue")
line.names <- c(A="Control", G="Large", P="Small")

funcdir  <- "../R_functions"

source(paste(funcdir, "Dataclean.R", sep="/"))

seldiff.algo <- c("effective", "family")[2] 
# "effective" is weighted by the breeders' reproductive success
# "family" accounts for within/among selection (from fig-seldiff.R)

h2.algo <- c("ols", "gls")[2]

sdl <- cleanSdl()

realseldiff <- function(ll, gen) {
	mm <- mean(sdl$Sdl[(if (gen != "F1") sdl$Line==ll else TRUE ) & sdl$Generation == gen])
	nboff <- sapply(sdl$ID[(if (gen != "F1") sdl$Line==ll else TRUE ) & sdl$Generation == gen], function(id) sum(c(as.character(sdl$Mother), as.character(sdl$Father)) == as.character(id), na.rm=TRUE))
	return(mean(nboff[nboff>0]/mean(nboff[nboff>0])*(sdl$Sdl[(if (gen != "F1") sdl$Line==ll else TRUE ) & sdl$Generation == gen][nboff > 0] - mm)))
}

gls.h2 <- function(R, S, Ne.sel, Ne.cont=Ne.sel, varP.sel, varP.cont=varP.sel, M.sel, M.cont=0, h2.0=0.2, verbose=TRUE, n.steps=10) {
	make.V <- function(h2) { # other parameters from the main function
		if (h2 < 0) h2 <- 0
		V <- matrix(NA, ncol=length(R), nrow=length(R))
		for (i in 1:length(R)) {
			for (j in i:length(R)) {
				V[i,j] <- V[j,i] <- h2*varP.sel*(1/M.sel[1] + i/Ne.sel) + h2*varP.cont*(1/M.cont[1]+i/Ne.cont)
				if (i==j) V[i,j] <- V[i,j] + varP.sel/M.sel[i]
			}
		}
		V
	}
	gsl.step <- function(R, S, V) solve(t(S)%*%solve(V)%*%S)%*%t(S)%*%solve(V)%*%R
		
	stopifnot(length(R) == length(S))
	h2 <- h2.0
	for (ss in seq_len(n.steps)) {
		V <- make.V(h2)
		h2 <- gsl.step(R,S,V)
		if (verbose) cat("Step", ss, ": h2=", h2, "\n")
	}
	list(h2=h2, var.h2=solve(t(S)%*%solve(make.V(h2))%*%S))
}

seldiff.table <- read.table("tab-seldiff.txt", header=TRUE)

pdf("fig-realized.pdf", width=4, height=4)
par(mar=c(5, 5, 2, 1), lwd=2)
plot(NULL, xlim=c(-6, 13), ylim=c(-0.7, 1.6), xlab="Cumulative Sdl selection differential (mm)", ylab="Sdl response to selection (mm)")
meanRef <- c(mean(sdl$Sdl[sdl$Generation=="F1"], na.rm=TRUE), tapply(sdl$Sdl[sdl$Line=="A"], sdl$Generation[sdl$Line=="A"], mean, na.rm=TRUE)[paste0("F",2:7)])
for (ll in names(line.col)) {
 	mml <- c(meanRef[1], tapply(sdl$Sdl[sdl$Line==ll], sdl$Generation[sdl$Line==ll], mean, na.rm=TRUE)[paste0("F",2:7)])
	if (seldiff.algo == "family") {
		cumdiff <- cumsum(c(0, seldiff.table[,paste0("Sdl.", ll)]))
	} else { # seldiff.algo == "effective"
		sdiffl <- sapply(paste0("F",1:6), function(gg) realseldiff(ll, gg))
		sdiffl <- tapply(sdl$Sdl[sdl$Line==ll & sdl$Breeder=="yes"], sdl$Generation[sdl$Line==ll & sdl$Breeder=="yes"], mean, na.rm=TRUE)[paste0("F",1:6)]-mml[1:6]
		cumdiff <- cumsum(c(0,sdiffl[1:6]))
	}
    resp <- mml-meanRef
    
    
    lines(cumdiff, resp, type="b", col=line.col[ll], pch=as.character(1:7))
    # arrows(x0=3:8, y0=mml-mmc-sel, y1=mml-mmc+sel,  angle=90, code=3, length=0.1, col=line.col[ll])
    # Realized h2 lines:
    if (ll != "A") {
		if (h2.algo == "ols") {
			h2ref <- lm(resp ~ 0+cumdiff)
			h2 <- list(h2=coef(h2ref), var.h2=coef(summary(h2ref))[1,"Std. Error"]^2)
		} else { #h2.algo == "gls"
			varP.sel  <- mean(tapply(sdl$Sdl[sdl$Line==ll], sdl$Gen[sdl$Line==ll], var, na.rm=TRUE)[paste0("F",1:7)])
			varP.cont <- 0 # mean(tapply(sdl$Sdl[sdl$Line=="A"], sdl$Gen[sdl$Line=="A"], var, na.rm=TRUE)[paste0("F",1:7)])
			M.sel     <- tapply(sdl$Sdl[sdl$Line==ll], sdl$Gen[sdl$Line==ll], length)[paste0("F",1:7)]
			M.cont    <- tapply(sdl$Sdl[sdl$Line=="A"], sdl$Gen[sdl$Line=="A"], length)[paste0("F",1:7)]
						
			h2 <- gls.h2(R=resp, S=cumdiff, Ne.sel=30, Ne.cont=30, varP.sel=varP.sel, varP.cont=varP.cont, M.sel=M.sel, M.cont=M.cont)
		}
        
        extrm <- cumdiff[which.max(abs(cumdiff))]
        lines(x=c(0,extrm), y=c(0, extrm*h2$h2), col=line.col[ll], lty=2)
        # print(summary(h2ref))
        if (ll == "P")
			text(x=0.4*extrm, y=0.6, col=line.col[ll], labels=paste0("h2 = ", round(h2$h2, digits=3), " +/- ", round(sqrt(h2$var.h2), digits=3)), cex=0.7)
		if (ll == "G")
			text(x=0.6*extrm, y=0.1, col=line.col[ll], labels=paste0("h2 = ", round(h2$h2, digits=3), " +/- ", round(sqrt(h2$var.h2), digits=3)), cex=0.7)		
    }
	mtext("B", side=3, line=0, adj=-0.2, cex=1.6)
}
dev.off()
```
