## Supplementary material for "Unidirectional response to bidirectional selection on body size II Quantitative genetics": Data files and analysis scripts: fig-response.R

```
#!/usr/bin/env Rscript

line.names <- c(A="Control", G="Large", P="Small")
line.col <- c(A="darkgray", G="red", P="blue")
line.names <- c(A="Control", G="Large", P="Small")
line.pch <- c(A=1, G=2, P=6)

funcdir  <- "../R_functions"

source(paste(funcdir, "Dataclean.R", sep="/"))

sdl <- cleanSdl()


pdf("fig-response.pdf", width=12, height=8)
par(mar=c(3, 5, 0.1, 2), oma=c(2, 0, 4, 0), lwd=2, xpd=NA)
layout(rbind(1:2,3:4))
par(cex=1/par("cex"))

# Sdl on the raw scale
mm <- tapply(sdl$Sdl, sdl$Generation, mean, na.rm=TRUE)[1:3]
se <- tapply(sdl$Sdl, sdl$Generation, sd, na.rm=TRUE)[1:3]/sqrt(tapply(sdl$Sdl, sdl$Generation, length)[1:3])
plot(mm, type="b", ylim=c(18, 22), xaxt="n", xlab="", ylab="Body size (mm)", xlim=c(1,8))
mtext("A", side=3, line=0, adj=-0.2, cex=1.6)
title("Original scale", line=2)
arrows(x0=1:2, y0=(mm-se)[1:2], y1=(mm+se)[1:2],  angle=90, code=3, length=0.1)
axis(1, at=1:8, paste0("F", 0:7))
for (ll in names(line.col)) {
    mml <- tapply(sdl$Sdl[sdl$Line==ll], sdl$Generation[sdl$Line==ll], mean, na.rm=TRUE)[paste0("F",2:7)]
    sel <- tapply(sdl$Sdl[sdl$Line==ll], sdl$Generation[sdl$Line==ll], sd, na.rm=TRUE)[paste0("F",2:7)]/sqrt(tapply(sdl$Sdl[sdl$Line==ll], sdl$Generation[sdl$Line==ll], length)[paste0("F",2:7)])
    lines(3:8, mml, type="b", col=line.col[ll], pch=line.pch[ll])
    arrows(x0=3:8, y0=mml-sel, y1=mml+sel,  angle=90, code=3, length=0.1, col=line.col[ll])
}

# Sdl on the control-centered scale
plot(rep(0, 3), type="b", ylim=c(-1, 1.8), xaxt="n", xlab="", ylab="Relative body size (mm)", xlim=c(1,8))
mtext("B", side=3, line=0, adj=-0.2, cex=1.6)
title("Control-centered", line=2)
#arrows(x0=1:2, y0=(se)[1:2], y1=-(se)[1:2],  angle=90, code=3, length=0.1)
axis(1, at=1:8, paste0("F", 0:7))
mmc <- tapply(sdl$Sdl[sdl$Line=="A"], sdl$Generation[sdl$Line=="A"], mean, na.rm=TRUE)[paste0("F",2:7)]
sec <- tapply(sdl$Sdl[sdl$Line=="A"], sdl$Generation[sdl$Line=="A"], sd, na.rm=TRUE)[paste0("F",2:7)]/sqrt(tapply(sdl$Sdl[sdl$Line=="A"], sdl$Generation[sdl$Line=="A"], length)[paste0("F",2:7)])
for (ll in names(line.col)) {
    mml <- tapply(sdl$Sdl[sdl$Line==ll], sdl$Generation[sdl$Line==ll], mean, na.rm=TRUE)[paste0("F",2:7)]
    sel <- tapply(sdl$Sdl[sdl$Line==ll], sdl$Generation[sdl$Line==ll], sd, na.rm=TRUE)[paste0("F",2:7)]/sqrt(tapply(sdl$Sdl[sdl$Line==ll], sdl$Generation[sdl$Line==ll], length)[paste0("F",2:7)])
    lines(3:8, mml-mmc, type="b", col=line.col[ll], pch=line.pch[ll])
    if (ll != "A")
		arrows(x0=3:8, y0=mml-mmc-sqrt(sel^2+sec^2), y1=mml-mmc+sqrt(sel^2+sec^2),  angle=90, code=3, length=0.1, col=line.col[ll])
}
legend("topleft", line.names[c("G","A","P")], lty=1, col=line.col[c("G","A","P")], pch=line.pch[c("G","A","P")])

# Mat on the raw scale
mm <- tapply(sdl$Sex != "I", sdl$Generation, mean, na.rm=TRUE)[1:3]
se <- tapply(sdl$Sex != "I", sdl$Generation, sd, na.rm=TRUE)[1:3]/sqrt(tapply(sdl$Sdl, sdl$Generation, length)[1:3])
plot(mm, type="b", ylim=c(0.5, 1), xaxt="n", xlab="Generation", ylab="Maturity", xlim=c(1,8))
mtext("C", side=3, line=0, adj=-0.2, cex=1.6)
arrows(x0=1:2, y0=(mm-se)[1:2], y1=(mm+se)[1:2],  angle=90, code=3, length=0.1)
axis(1, at=1:8, paste0("F", 0:7))
for (ll in names(line.col)) {
    mml <- tapply((sdl$Sex != "I")[sdl$Line==ll], sdl$Generation[sdl$Line==ll], mean, na.rm=TRUE)[paste0("F",2:7)]
    sel <- tapply((sdl$Sex != "I")[sdl$Line==ll], sdl$Generation[sdl$Line==ll], sd, na.rm=TRUE)[paste0("F",2:7)]/sqrt(tapply(sdl$Sdl[sdl$Line==ll], sdl$Generation[sdl$Line==ll], length)[paste0("F",2:7)])
    lines(3:8, mml, type="b", col=line.col[ll], pch=line.pch[ll])
    arrows(x0=3:8, y0=mml-sel, y1=mml+sel,  angle=90, code=3, length=0.1, col=line.col[ll])
}

# Mat on the control-centered scale
plot(rep(0, 3), type="b", ylim=c(-0.25, 0.25), xaxt="n", xlab="Generation", ylab="Relative maturity", xlim=c(1,8))
mtext("D", side=3, line=0, adj=-0.2, cex=1.6)
# arrows(x0=1:2, y0=(se)[1:2], y1=-(se)[1:2],  angle=90, code=3, length=0.1)
axis(1, at=1:8, paste0("F", 0:7))
mmc <- tapply((sdl$Sex != "I")[sdl$Line=="A"], sdl$Generation[sdl$Line=="A"], mean, na.rm=TRUE)[paste0("F",2:7)]
sec <- tapply((sdl$Sex != "I")[sdl$Line=="A"], sdl$Generation[sdl$Line=="A"], sd, na.rm=TRUE)[paste0("F",2:7)]/sqrt(tapply((sdl$Sex != "I")[sdl$Line=="A"], sdl$Generation[sdl$Line=="A"], length)[paste0("F",2:7)])
for (ll in names(line.col)) {
    mml <- tapply((sdl$Sex != "I")[sdl$Line==ll], sdl$Generation[sdl$Line==ll], mean, na.rm=TRUE)[paste0("F",2:7)]
    sel <- tapply((sdl$Sex != "I")[sdl$Line==ll], sdl$Generation[sdl$Line==ll], sd, na.rm=TRUE)[paste0("F",2:7)]/sqrt(tapply((sdl$Sex != "I")[sdl$Line==ll], sdl$Generation[sdl$Line==ll], length)[paste0("F",2:7)])
    lines(3:8, mml-mmc, type="b", col=line.col[ll], pch=line.pch[ll])
    if (ll != "A")
		arrows(x0=3:8, y0=mml-mmc-sqrt(sel^2+sec^2), y1=mml-mmc+sqrt(sel^2+sec^2),  angle=90, code=3, length=0.1, col=line.col[ll])}


dev.off()
```
