## Supplementary material for "Unidirectional response to bidirectional selection on body size II Quantitative genetics": Data files and analysis scripts: fig-seldiff.R

```
#!/usr/bin/env Rscript

# A: control, G: large, P: small
line.col <- c(A="darkgray", G="red", P="blue")
line.names <- c(A="Control", G="Large", P="Small")

lines <- setNames(names(line.col), names(line.col))

funcdir  <- "../R_functions"

makeTransparent<-function(someColor, alpha=100)
# From StackOverflow: https://stackoverflow.com/questions/8047668/transparent-equivalent-of-given-color
{
  newColor<-col2rgb(someColor)
  apply(newColor, 2, function(curcoldata){rgb(red=curcoldata[1], green=curcoldata[2],
    blue=curcoldata[3],alpha=alpha, maxColorValue=255)})
}

breedertag <- "Selected_Breeder"

source(paste(funcdir, "Dataclean.R", sep="/"))

sdl <- cleanSdl()

# Total, within-family, and among-family effective selection gradients

rdiff.total <- lapply(lines, function(ll) {
	sapply(1:6, function(gg) {
        filter <- sdl$Generation == paste0("F",gg) & if(gg==1) TRUE else sdl$Line==ll
        fam <- sdl$Family[filter]
        ss <- sdl$Sdl[filter]
        mm <- sdl$Sex[filter] != "I"
        ff <- sdl[,breedertag][filter] == "yes" & sdl$Line[filter] == ll
        ff <- ifelse(sdl$Line[filter] == ll, sapply(sdl$ID[filter], function(id) sum(c(as.character(sdl$Mother), as.character(sdl$Father)) == as.character(id), na.rm=TRUE)), 0)
        sdl.by.fam <- tapply(ss, fam, mean)
        mat.by.fam <- tapply(mm, fam, mean)
        sdlfam <- sdl.by.fam[fam]
        matfam <- mat.by.fam[fam]
        c(Sdl=mean((ss)*ff/mean(ff), na.rm=TRUE)-mean(ss, na.rm=TRUE), MatTRUE=mean((mm)*ff/mean(ff), na.rm=TRUE)-mean(mm, na.rm=TRUE))
    })}
)

rdiff.withinfam <- lapply(lines, function(ll) {
	sapply(1:6, function(gg) {
        filter <- sdl$Generation == paste0("F",gg) & if(gg==1) TRUE else sdl$Line==ll
        fam <- sdl$Family[filter]
        ss <- sdl$Sdl[filter]
        mm <- sdl$Sex[filter] != "I"
        ff <- sdl[,breedertag][filter] == "yes" & sdl$Line[filter] == ll
        ff <- ifelse(sdl$Line[filter] == ll, sapply(sdl$ID[filter], function(id) sum(c(as.character(sdl$Mother), as.character(sdl$Father)) == as.character(id), na.rm=TRUE)), 0)
        sdl.by.fam <- tapply(ss, fam, mean)
        mat.by.fam <- tapply(mm, fam, mean)
        sdlfam <- sdl.by.fam[fam]
        matfam <- mat.by.fam[fam]
        c(Sdl=mean((ss-sdlfam)*ff/mean(ff), na.rm=TRUE), MatTRUE=mean((mm-matfam)*ff/mean(ff), na.rm=TRUE))
    })}
)

rdiff.betweenfam <- lapply(lines, function(ll) {
	sapply(1:6, function(gg) {
        filter <- sdl$Generation == paste0("F",gg) & if(gg==1) TRUE else sdl$Line==ll
        fam <- sdl$Family[filter]
        ss <- sdl$Sdl[filter]
        mm <- sdl$Sex[filter] != "I"
        ff <- sdl[,breedertag][filter] == "yes" & sdl$Line[filter] == ll
        ff <- ifelse(sdl$Line[filter] == ll, sapply(sdl$ID[filter], function(id) sum(c(as.character(sdl$Mother), as.character(sdl$Father)) == as.character(id), na.rm=TRUE)), 0)
        sdl.by.fam <- tapply(ss, fam, mean)
        mat.by.fam <- tapply(mm, fam, mean)
        sdlfam <- sdl.by.fam[fam]
        matfam <- mat.by.fam[fam]
        c(Sdl=mean((sdlfam)*ff/mean(ff), na.rm=TRUE)-mean(ss, na.rm=TRUE), MatTRUE=mean((matfam)*ff/mean(ff), na.rm=TRUE)-mean(mm, na.rm=TRUE))
    })}
)


pdf("fig-seldiff.pdf", width=8, height=4)
	layout(t(1:2))
	par(lwd=2, mar=c(5, 5, 2, 1))
	
	plot(NULL, xlim=c(0,6), ylim=c(-5, 13), ylab="Selection Differential on Body size (mm)", xlab="Generation")
	for (ll in lines) {            
	    lines(0:6, cumsum(c(0,rdiff.total[[ll]]["Sdl",])), type="b", col=line.col[ll], lty=1, lwd=3)
	    lines(0:6, cumsum(c(0,rdiff.withinfam[[ll]]["Sdl",])), type="b", col=line.col[ll], lty=2)
	    lines(0:6, cumsum(c(0,rdiff.betweenfam[[ll]]["Sdl",])), type="b", col=line.col[ll], lty=1)
	}
	
	
	plot(NULL, xlim=c(0,6), ylim=c(-0.1, 1), ylab="Selection Differential on Maturity", xlab="Generation")
	for (ll in lines) {            
	    lines(0:6, cumsum(c(0,rdiff.total[[ll]]["MatTRUE",])), type="b", col=line.col[ll], lty=1, lwd=3)
	    lines(0:6, cumsum(c(0,rdiff.withinfam[[ll]]["MatTRUE",])), type="b", col=line.col[ll], lty=2)
	    lines(0:6, cumsum(c(0,rdiff.betweenfam[[ll]]["MatTRUE",])), type="b", col=line.col[ll], lty=1)
	}
	
	legend("topleft", lwd=c(3,1,1), lty=c(1,1,2), legend=c("Total","Among families", "Within families"))

dev.off()

among.ratio <- read.table("tab-variances-cor.txt", header=TRUE)

seldiff.tab <- NULL

pdf("fig-seldiff-pred.pdf", width=8, height=4)
	layout(t(1:2))
	par(lwd=2, mar=c(5, 5, 2, 1))

	tt <- "Sdl"
	plot(NULL, xlim=c(1,7), ylim=c(-6,15), xaxt="n", xlab="", ylab="Sdl effective differential (mm)")
	axis(1, at=1:7, labels=paste0("F",1:7))
	
	for (ll in lines) {
		response.within <- rdiff.withinfam[[ll]][tt,]/(2*(1-among.ratio[2:7,paste0("Sdl",ll)]))
		response.between <- rdiff.betweenfam[[ll]][tt,]/(2*among.ratio[2:7,paste0("Sdl",ll)])
		seldiff.tab <- cbind(seldiff.tab, response.within+response.between)
		colnames(seldiff.tab)[ncol(seldiff.tab)] <- paste0(tt, ".", ll)
		lines(c(0, cumsum(response.within)), col=line.col[ll], lty=2)
		lines(c(0, cumsum(response.between)), col=line.col[ll], lty=1, lwd=1)
		lines(c(0, cumsum(response.within+response.between)), col=line.col[ll], lwd=3)
		lines(c(0, cumsum(rdiff.total[[ll]][tt,])), col=makeTransparent(line.col[ll]), lwd=3, lty=3)
	}
	legend("topleft", lwd=c(1,2,3,3), lty=c(1,2,1,3), col=c("black","black","black", makeTransparent("black")), legend=c("Among families", "Within families", "Total", "Raw"), bty="n")

	tt <- "MatTRUE"
	plot(NULL, xlim=c(1,7), ylim=c(-0.2,0.9), xaxt="n", xlab="", ylab="Maturity effective differential")
	axis(1, at=1:7, labels=paste0("F",1:7))
	
	for (ll in lines) {
		response.within <- rdiff.withinfam[[ll]][tt,]/(2*(1-among.ratio[2:7,paste0("Sdl",ll)]))
		response.between <- rdiff.betweenfam[[ll]][tt,]/(2*among.ratio[2:7,paste0("Sdl",ll)])
		seldiff.tab <- cbind(seldiff.tab, response.within+response.between)
		colnames(seldiff.tab)[ncol(seldiff.tab)] <- paste0(tt, ".", ll)
		lines(c(0, cumsum(response.within)), col=line.col[ll], lty=2)
		lines(c(0, cumsum(response.between)), col=line.col[ll], lty=1, lwd=1)
		lines(c(0, cumsum(response.within+response.between)), col=line.col[ll], lwd=3)
		lines(c(0, cumsum(rdiff.total[[ll]][tt,])), col=makeTransparent(line.col[ll]), lwd=3, lty=3)
	}
	legend("topleft", lwd=c(1,2,3,3), lty=c(1,2,1,3), col=c("black","black","black", makeTransparent("black")), legend=c("Among families", "Within families", "Total", "Raw"), bty="n")

dev.off()

rownames(seldiff.tab) <- paste0("F", 1:6)
write.table(round(seldiff.tab, digits=4), file="tab-seldiff.txt", sep="\t", quote=FALSE)
```
