## Supplementary material for "Unidirectional response to bidirectional selection on body size II Quantitative genetics": Data files and analysis scripts: fig-stationarity.R

```
#!/usr/bin/env Rscript

source("./models.R")

tokeep <- c(
	"traitSdl:traitSdl.animal" = expression(V[G](Sdl)), #"$V_G(\\mathrm{Sdl})$", 
	"traitMat:traitMat.animal" = expression(V[G](Mat)), #"$V_G(\\mathrm{Mat})$", 
	"traitSdl:traitMat.animal" = expression(C[G](Sdl,Mat)),
	"traitSdl:traitSdl.units"= expression(V[E](Sdl)), #"$V_E(\\mathrm{Sdl})$", 
	"traitSdl:traitMat.units"= expression(C[E](Sdl,Mat))) #"$C_E(\\mathrm{Sdl,Mat})$",
#~ 	"traitSdl.gen" = expression(V[F](Sdl)), #"$V_F(\\mathrm{Sdl})$",
#~ 	"traitMat.gen" = expression(V[F](Mat))) #"$V_F(\\mathrm{Mat})$")
	

mod <- mod.all[[1]]

hd <- heidel.diag(mod$VCV)

hd.pv <- sapply(hd, function(x) x[,'pvalue'])

hd.pv <- hd.pv[names(tokeep),]


pdf("fig-stationarity.pdf", width=6, height=5)
plot(x=rep(1:nrow(hd.pv), each=ncol(hd.pv)), y=t(hd.pv), xaxt="n", ylab="Stationarity test p-value", ylim=0:1, xlab="")
axis(1, at=1:nrow(hd.pv), tokeep, tick=FALSE)
abline(h=0.05, col="red", lty=2)
dev.off()
```
