## Supplementary material for "Unidirectional response to bidirectional selection on body size II Quantitative genetics": Data files and analysis scripts: fig-variances.R

```
#!/usr/bin/env Rscript

line.col <- c(A="darkgray", G="red", P="blue")
line.names <- c(A="Control", G="Large", P="Small")
line.pch <- c(A=1, G=2, P=6)

vartotmethod <- c("sumWithinAmong", "phenvar")[1]

funcdir  <- "../R_functions"

lines <- setNames(names(line.col), names(line.col))

source(paste(funcdir, "Dataclean.R", sep="/"))

sdl <- cleanSdl()
fam <- cleanFam(sdl)

varByGen       <- tapply(sdl$Sdl, sdl$Gen, var, na.rm=TRUE)
varWithinByGen <- tapply(fam$VarSdl, fam$Gen, mean, na.rm=TRUE)
varAmongByGen  <- tapply(fam$MeanSdl, fam$Gen, var, na.rm=TRUE)

varByLineGen       <- lapply(lines, function(ll) c(varByGen[1:2],tapply(sdl$Sdl[sdl$Line==ll], sdl$Gen[sdl$Line==ll], var, na.rm=TRUE)[3:8]))
varWithinByLineGen <- lapply(lines, function(ll) c(varWithinByGen[1:2],tapply(fam$VarSdl[fam$Line==ll], fam$Gen[fam$Line==ll], mean, na.rm=TRUE)[3:8]))
varAmongByLineGen  <- lapply(lines, function(ll) c(varAmongByGen[1:2], tapply(fam$MeanSdl[fam$Line==ll], fam$Gen[fam$Line==ll], var, na.rm=TRUE)[3:8]))

refvarByGen <- if (vartotmethod == "phenvar") varByGen else varWithinByGen + varAmongByGen
refvarByLineGen <- if (vartotmethod == "phenvar") varByLineGen else mapply(varWithinByLineGen, varAmongByLineGen, FUN='+', SIMPLIFY=FALSE)

pdf("fig-variances.pdf", width=8, height=4)
layout(t(1:2))
par(lwd=2)

   # variance among family means
plot(varAmongByGen, type="b", xlim=c(1,8), ylim=c(0, 4.5), xaxt="n", xlab="Generation", ylab="Var Sdl (mm^2)", main="Among families")
for (ll in lines) lines(3:8, varAmongByLineGen[[ll]][paste0("F",2:7)], type="b", col=line.col[ll], pch=line.pch[ll])
axis(1, at=1:8, paste0("F", 0:7))

   # variance within families
plot(varWithinByGen, type="b", xlim=c(1,8), ylim=c(0, 9.5), xaxt="n", xlab="Generation", ylab="Var Sdl (mm^2)", main="Within families")
for (ll in lines) lines(3:8, varWithinByLineGen[[ll]][paste0("F",2:7)], type="b", col=line.col[ll], pch=line.pch[ll])
axis(1, at=1:8, paste0("F", 0:7))

dev.off()

cor.relatives <- function(sdl, trait="Sdl") {
	if (nrow(sdl) == 0) return(NA)
	allpairs <- do.call(rbind, lapply(1:nrow(sdl), function(i) {
			ff <- sdl$Family[i]
			oth <- sdl[,trait][sdl$Family == ff & -i]
			cbind(rep(sdl[,trait][i], length(oth)), oth)
	}))
	cor(allpairs[,1], allpairs[,2], use="complete")
}

cors <- list(Sdl=list(
			corByGen <- sapply(split(sdl, sdl$Gen), cor.relatives, trait="Sdl"),
			corByLineGen = lapply(lines, function(ll) c(corByGen[1:2], sapply(split(sdl[sdl$Line==ll,], sdl$Gen[sdl$Line==ll]), cor.relatives, trait="Sdl")[3:8]))
			),
			Mat=list(
			corByGen <- sapply(split(sdl, sdl$Gen), cor.relatives, trait="Mat"),
			corByLineGen = lapply(lines, function(ll) c(corByGen[1:2], sapply(split(sdl[sdl$Line==ll,], sdl$Gen[sdl$Line==ll]), cor.relatives, trait="Mat")[3:8]))
			))

pdf("fig-variances-cor.pdf", width=8, height=4)
	layout(t(1:2))
	par(lwd=2, mar=c(4,4,1,1))
	
	plot(cors$Sdl$corByGen, type="b", xlim=c(1,8), ylim=c(0, 1), xaxt="n", xlab="Generation", ylab="Within-family Sdl correlation")
	for (ll in lines) 
		lines(3:8, cors$Sdl$corByLineGen[[ll]][paste0("F",2:7)], type="b", col=line.col[ll], pch=line.pch[ll])
	axis(1, at=1:8, paste0("F", 0:7))
	
#~ 	plot(varAmongByGen/refvarByGen, type="b", xlim=c(1,8), ylim=c(0, 1), xaxt="n", xlab="Generation", ylab="Relative Among-family Var Sdl")
#~ 	for (ll in lines) 
#~ 		lines(3:8, (varAmongByLineGen[[ll]]/refvarByLineGen[[ll]])[paste0("F",2:7)], type="b", col=line.col[ll], pch=line.pch[ll])
#~ 	axis(1, at=1:8, paste0("F", 0:7))

	plot(cors$Mat$corByGen, type="b", xlim=c(1,8), ylim=c(0, 1), xaxt="n", xlab="Generation", ylab="Within-family Maturity correlation")
	for (ll in lines) 
		lines(3:8, cors$Mat$corByLineGen[[ll]][paste0("F",2:7)], type="b", col=line.col[ll], pch=line.pch[ll])
	axis(1, at=1:8, paste0("F", 0:7))	

dev.off()


tt <- cbind(SdlAll=cors$Sdl$corByGen, SdlG=cors$Sdl$corByLineGen$G, SdlA=cors$Sdl$corByLineGen$A, SdlP=cors$Sdl$corByLineGen$P, 
			MatAll=cors$Mat$corByGen, MatG=cors$Mat$corByLineGen$G, MatA=cors$Mat$corByLineGen$A, MatP=cors$Mat$corByLineGen$P)
			
write.table(round(tt, digits=2), file="tab-variances-cor.txt", sep="\t", quote=FALSE)
```
