## Supplementary material for "Unidirectional response to bidirectional selection on body size II Quantitative genetics": Data files and analysis scripts: models.R

```
#!/usr/bin/env Rscript

# Run the MCMCglmm models (if necessary) or load them from the cache
# The models are stored as lists in the mod.all global variable

degener <- function(gener) as.numeric(unlist(sapply(strsplit(as.character(gener),split="F"), "[", 2)))
makeOldID <- function(fam, sex, line) 
    paste0(sub("\\.", "", as.character(fam)), as.character(sex), as.character(round(runif(length(fam), 1e6, 1e7-1))), as.character(line))
    
    
funcdir  <- "../R_functions"
cachedir <- "./cache"

source(paste(funcdir, "Dataclean.R", sep="/"))
source(paste(funcdir, "Inbreeding.R", sep="/"))
source(paste(funcdir, "AnimalModel.R", sep="/"))

modname <- c("GRAFi", "GRAFc") #  One or several model names
# * G or g : additive genetic variances (with and without correlations, respectively)
# * R or r : residual variance (with and without correlations). 
# * A or a : aquarium effect (can be absent)
# * F or f : generation effect (can be absent)
# * i      : include an inbreeding fixed effect
# * c      : fit the model only on the control line

sdl <- cleanSdl()
sdl$ID <- ifelse(is.na(sdl$ID), makeOldID(sdl$Family, sdl$Sex, sdl$Line), as.character(sdl$ID))

ped.all <- getped.glmm(sdl)

mydata.all <- data.frame(
	Sdl=sdl$Sdl, 
	Mat=ifelse(sdl$Sex != "I", 1, 0), 
	animal=ped.all$animal, 
	dom=ped.all$animal, 
	gen=factor(sdl$Generation),
	sex=sdl$Sex, 
	aqua=sdl$Family, 
	inb=Inbreeding(sdl),
	Line=sdl$Line, 
	ID=sdl$ID, 
	Family=sdl$Family, 
	Father=sdl$Father, 
	Mother=sdl$Mother)

variance.prior <- list(V=diag(2),nu=2)
resid.prior <- list(V=diag(2), nu=2, fix=2)
burnin <- 10000
nitt   <- 1000000 #1000000

default.args <- list(
	fixed   = 'cbind(Sdl,Mat) ~trait - 1',
	family  = c("gaussian","ordinal"),
	pedigree= ped.all,
	data    = mydata.all,
	nitt    = nitt,
	burnin  = burnin,
	thin    = 100,
	pr      = TRUE)
	

mod.args <- sapply(modname, function(nn) {
		args <- default.args
		
		args$rcov <- if (grepl(nn, pattern="r")) ~idh(trait):units else if (grepl(nn, pattern="R")) ~us(trait):units else stop()
		
		randomanimal <- if (grepl(nn, pattern="g")) "idh(trait):animal" else if (grepl(nn, pattern="G")) "us(trait):animal" else ""
		randomaqua   <- if (grepl(nn, pattern="a")) "+idh(trait):aqua" else if (grepl(nn, pattern="A")) "+us(trait):aqua" else ""
		randomgen    <- if (grepl(nn, pattern="f")) "+idh(trait):gen" else if (grepl(nn, pattern="F")) "+us(trait):gen" else ""
		
		fixedinb    <- if (grepl(nn, pattern="i")) "+trait:inb" else  ""
		
		if (grepl(nn, pattern="c")) {
			sdl.controls <- sdl[sdl$Generation %in% c("F0","F1") | sdl$Line == "A",]
			ped.controls <- getped.glmm(sdl.controls)
			args$pedigree <- ped.controls
			args$data <- data.frame(
				Sdl=sdl.controls$Sdl, 
				Mat=ifelse(sdl.controls$Sex != "I", 1, 0), 
				animal=ped.controls$animal, 
				dom=ped.controls$animal, 
				gen=factor(sdl.controls$Generation),
				sex=sdl.controls$Sex, 
				aqua=sdl.controls$Family, 
				inb=Inbreeding(sdl.controls),
				Line=sdl.controls$Line, 
				ID=sdl.controls$ID, 
				Family=sdl.controls$Family, 
				Father=sdl.controls$Father, 
				Mother=sdl.controls$Mother)
		}
		
		args$random <- as.formula(paste0("~", randomanimal, randomaqua, randomgen))
		args$fixed <- as.formula(paste0(args$fixed, fixedinb))
		
		priorG <- sapply(paste0("G", 1:sum(c(randomanimal, randomaqua, randomgen)!="")), function(g) variance.prior, simplify=FALSE)
		args$prior <- list(R=resid.prior, G=priorG)
		
		args
	}, USE.NAMES=TRUE, simplify=FALSE)
	


mod.all <- sapply(modname, function(nn) loadORrun(nn, mod.args[[nn]]), USE.NAMES=TRUE, simplify=FALSE)
```
