## Supplementary material for "Unidirectional response to bidirectional selection on body size II Quantitative genetics": Data files and analysis scripts: README

*************************** README *****************************

Scripts for data analysis and figure plotting for the manuscript
"Unidirectional response to bidirectional selection on body size. II:
quantitative genetics" by A. Le Rouzic et al. 

This directory ("analyses") contains 16 R scripts. Some run MCMC models
(stored in the ./cache directory), some generate tables (in format .txt or
.tex), and some generate figures (pdf). 

Scripts rely on functions that can be found in ../R_functions/

In theory, paths are managed automatically, and scripts can be called
from anywhere. Figures and tables are stored in the current directory
(not necessarily in the analysis directory). 

Analyses were run on R 3.6 or R 4.0. on Ubuntu Linux (18.4 or 20.4). 
They need the following packages:
* MASS
* parallel
* ellipse
* QGglmm
* shape
* xtable
* abind
* kinship2
* GENLIB
* stats4

Five scripts need to be run before the rest (as they generate datasets
that are resused elsewhere) :
* model.R
* fig-genetictrends.R
* fig-gradients.R
* fig-seldiff.R
* fig-variances.R

Note that with the default parameters, models can be quite heavy to run 
and can be too demanding (CPU and RAM) for desktop computers. The size of 
the model files (which save all posteriors for thousands effets) can
be substantial.

These scripts are provided to ensure the reproducibility of our analyses. 
Making science reproducible is often confused with making software reusable;
writing generic software was not our intention here. In particular,
The code was not written to be easily run on other operating systems or applied
to other datasets. In case or reusing substantial parts of the code, 
please cite the original publication. 

Any question or suggestion can be sent to 
Arnaud Le Rouzic <>. 
