## Supplementary material for "Unidirectional response to bidirectional selection on body size II Quantitative genetics": Data files and analysis scripts: tab-convergence.R

```
#!/usr/bin/env Rscript

source("./models.R")

tokeep <- c(
	"traitSdl:traitSdl.animal" = expression(V[G](Sdl)), #"$V_G(\\mathrm{Sdl})$", 
	"traitMat:traitMat.animal" = expression(V[G](Mat)), #"$V_G(\\mathrm{Mat})$", 
	"traitSdl:traitMat.animal" = expression(C[G](Sdl,Mat)),
	"traitSdl:traitSdl.units"= expression(V[E](Sdl)), #"$V_E(\\mathrm{Sdl})$", 
	"traitSdl:traitMat.units"= expression(C[E](Sdl,Mat))) 
	

args <- mod.args[[1]]
args$thin <- 1
args$nitt <- min(args$nitt, 100000)

convmod <- loadORrun(paste0(names(mod.all)[1], "-thin1"), args)

autocorr.tab <- cbind(t(autocorr.diag(convmod$VCV, lags=c(1, 5, 10, 50, 100, 500))), "Eff. Size"=effectiveSize(mod.all[[1]]$VCV))

autocorr.tab <- autocorr.tab[names(tokeep),]
rownames(autocorr.tab) <- tokeep


library(xtable)

print(xtable(autocorr.tab, digits=2), floating=FALSE, sanitize.rownames.function=identity, file="tab-convergence.tex")
```
