## Supplementary material for "Unidirectional response to bidirectional selection on body size II Quantitative genetics": Data files and analysis scripts: tab-posteriors.R

```
#!/usr/bin/env Rscript

source("./models.R")

library(xtable)

maternal.effect <- function(motherfam, Sol, traitmother="Sdl", traitoffsp=traitmother) {
	# mother fam is a named vector of fish indexes. Names are families, and indexes are the index of the mother from each family.

	sapply(1:nrow(Sol), function(i) {
		aquaTrait <- paste0("trait", traitoffsp, ".aqua.", names(motherfam)[!is.na(motherfam)])
		animalTrait <- paste0("trait", traitmother, ".animal.", as.character(motherfam[!is.na(motherfam)]))
		aquaBLUP <- Sol[i,aquaTrait]
		motherBLUP <- Sol[i,animalTrait]
		coef(lm(aquaBLUP ~ motherBLUP))[2]
	})
}

make.posterior.table <- function(model, motherfam=NULL) {
	# If motherfam is not provided, maternal effects will not be computed
	xlong <- cbind(do.call(rbind, model$VCV), do.call(rbind, model$Sol))
	
	cov.animal <- "traitSdl:traitMat.animal"
	cov.aqua <- "traitSdl:traitMat.aqua"
	cov.gen <- "traitSdl:traitMat.gen"
	cov.units <- "traitSdl:traitMat.units"
	
	if (!cov.animal %in% colnames (xlong)) cov.animal <- NA  
	if (!cov.animal %in% colnames (xlong)) cov.animal <- NA 
	if (!cov.animal %in% colnames (xlong)) cov.animal <- NA 
	
	sdl.inb <- "traitSdl:inb"
	mat.inb <- "traitMat:inb"
	if (!sdl.inb %in% colnames (xlong)) sdl.inb <- NA
	if (!mat.inb %in% colnames (xlong)) mat.inb <- NA
	
	sdl.animal <- if (!is.na(cov.animal)) "traitSdl:traitSdl.animal" else "traitSdl.animal"
	mat.animal <- if (!is.na(cov.animal)) "traitMat:traitMat.animal" else "traitMat.animal"
	sdl.aqua <- if (!is.na(cov.aqua)) "traitSdl:traitSdl.aqua" else "traitSdl.aqua"
	mat.aqua <- if (!is.na(cov.aqua)) "traitMat:traitMat.aqua" else "traitMat.aqua"
	sdl.gen <- if (!is.na(cov.gen)) "traitSdl:traitSdl.gen" else "traitSdl.gen"
	mat.gen <- if (!is.na(cov.gen)) "traitMat:traitMat.gen" else "traitMat.gen"	
	sdl.units <- "traitSdl:traitSdl.units"
	mat.units <- "traitMat:traitMat.units"
	
	posterior.tab <- cbind(
		`$\\bm{V_G}(\\mathrm{Sdl})$`      = xlong[,sdl.animal],
		`$h^2(\\mathrm{Sdl})$`            = xlong[,sdl.animal]/(xlong[,sdl.animal]+xlong[,sdl.units]+xlong[,sdl.aqua]),
		`$\\bm{V_G}(\\mathrm{Mat})$`      = xlong[,mat.animal],
		`$h^2(\\mathrm{Mat})$`            = xlong[,mat.animal]/(xlong[,mat.animal]+xlong[,mat.units]+xlong[,mat.aqua]),
		`$\\bm{C_G}(\\mathrm{Sdl,Mat})$`  = if (is.na(cov.animal)) NA else xlong[,cov.animal],
		`$r_G(\\mathrm{Sdl,Mat})$`        = if (is.na(cov.animal)) NA else xlong[,cov.animal]/sqrt(xlong[,sdl.animal])/sqrt(xlong[,mat.animal]),
		`$\\bm{V_E}(\\mathrm{Sdl})$`      = xlong[,sdl.units],
		`$\\bm{C_E}(\\mathrm{Sdl,Mat})$`  = xlong[,cov.units],
		`$r_E(\\mathrm{Sdl,Mat})$`        = xlong[,cov.units] / sqrt(xlong[,sdl.units])/sqrt(xlong[,mat.units]),
		`$\\bm{V_A}(\\mathrm{Sdl})$`      = xlong[,sdl.aqua],
		`$\\bm{V_A}(\\mathrm{Mat})$`      = xlong[,mat.aqua],
		`$\\bm{C_A}(\\mathrm{Sdl,Mat})$` = if (is.na(cov.aqua)) NA else xlong[,cov.aqua],
		`$r_A(\\mathrm{Sdl,Mat})$`        = if (is.na(cov.aqua)) NA else xlong[,cov.aqua]/sqrt(xlong[,sdl.aqua])/sqrt(xlong[,mat.aqua]),
		`$\\bm{V_F}(\\mathrm{Sdl})$`      = xlong[,sdl.gen],
		`$\\bm{V_F}(\\mathrm{Mat})$`      = xlong[,mat.gen],
		`$\\bm{C_F}(\\mathrm{Sdl,Mat})$` = if (is.na(cov.gen)) NA else xlong[,cov.gen],
		`$r_F(\\mathrm{Sdl,Mat})$`        = if (is.na(cov.gen)) NA else xlong[,cov.gen]/sqrt(xlong[,sdl.gen])/sqrt(xlong[,mat.gen]),
		`$\\bm{f}_\\mathrm{Sdl}$`         = if (is.na(sdl.inb)) NA else xlong[,sdl.inb],
		`$\\bm{f}_\\mathrm{Mat}$`         = if (is.na(mat.inb)) NA else xlong[,mat.inb],
		`$m_\\mathrm{Sdl}$`               = if (is.null(motherfam)) NA else maternal.effect(motherfam, xlong, "Sdl","Sdl"),
		`$m_\\mathrm{Mat}$` 	          = if (is.null(motherfam)) NA else maternal.effect(motherfam, xlong, "Mat","Mat")
	)
}


# Full model
sdl.full <- mod.args[["GRAFi"]]$data
motherfam.full <- setNames(match(sapply(levels(sdl.full$aqua), function(ff) sdl.full$Mother[sdl.full$aqua==ff][1]), sdl.full$ID), levels(sdl.full$aqua))

posteriortab.full <- make.posterior.table(mod.all[["GRAFi"]], motherfam.full)
quantiletab.full <- round(apply(posteriortab.full, 2, quantile, prob=c(0.025, 0.5, 0.975)), digits=3)

# Predictive model
sdl.pred <- mod.args[["GRAFc"]]$data
motherfam.pred <- setNames(match(sapply(levels(sdl.pred$aqua), function(ff) sdl.pred$Mother[sdl.pred$aqua==ff][1]), sdl.pred$ID), levels(sdl.pred$aqua))

posteriortab.pred <- make.posterior.table(mod.all[["GRAFc"]], motherfam.pred)
quantiletab.pred <- round(apply(posteriortab.pred, 2, quantile, prob=c(0.025, 0.5, 0.975), na.rm=TRUE), digits=3)

addtorow <- list(pos=list(0,0), 
	command=c("&\\multicolumn{3}{c}{Full} &  \\multicolumn{3}{c}{Predictive} \\\\\n","& 2.5\\% & median & 97.5\\% & 2.5\\% & median & 97.5\\% \\\\\n"))

print(xtable(cbind(t(quantiletab.full), t(quantiletab.pred)), align="cllllll"), 
	floating=FALSE, 
	sanitize.rownames.function=identity, 
	add.to.row = addtorow, 
	include.colnames = FALSE,
	hline.after = c(0),
	file="tab-posteriors.tex")
```
