## Supplementary material for "Unidirectional response to bidirectional selection on body size II Quantitative genetics": Data files and analysis scripts: Absolute_fitness.R

```
Abs_fitness <- function (Data) {
# Loop through beerders
breeders <- levels(factor(Data$ID[!is.na(Data$ID)]))
X <- NULL
for(i in 1:length(breeders)){
    Sdl <- Data$Sdl[Data$ID == breeders[i] & !is.na(Data$ID)];
    mother_progeny <- dim(Data[Data$Mother == breeders[i] & Data$Phenotyping == "P3" & !is.na(Data$Mother),])[1];
    father_progeny <- dim(Data[Data$Father == breeders[i] & Data$Phenotyping == "P3" & !is.na(Data$Father),])[1];
    Gen <- levels(factor(Data$Generation[Data$ID == breeders[i] & !is.na(Data$ID)]));
    Line <- levels(factor(Data$Line[Data$ID == breeders[i] & !is.na(Data$ID)]));
    Sex <- levels(factor(Data$Sex[Data$ID == breeders[i] & !is.na(Data$ID)]));
  X <- rbind(X, data.frame(Generation = Gen, Line = Line, Breeder = breeders[i], Sdl = Sdl, Sex = Sex, P3.progeny = mother_progeny + father_progeny))	
  }
return(X)}
```
