## Supplementary material for "Unidirectional response to bidirectional selection on body size II Quantitative genetics": Data files and analysis scripts: AnimalModel.R

```
library(parallel)
mc.cores <- min(64, detectCores() -1)
mc.cores <- 20
library(MCMCglmm)

my.MCMCglmm <- function(..., nitt=13000, burnin=3000, mc.cores=min(detectCores()-1, 32), nbsplit=mc.cores) {
    stopifnot(nitt > burnin)
    nitt <- round(burnin+(nitt-burnin)/nbsplit)
    mcmcmc <- mclapply(1:nbsplit, function(i) MCMCglmm(..., burnin=burnin, nitt=max(1000, nitt), verbose=(i==1)), mc.cores=mc.cores)
    list(
        X=     mcmcmc[[1]]$X,          # All the same
        Z=     mcmcmc[[1]]$Z,          # All the same
        ZR=    mcmcmc[[1]]$ZR,         # All the same
        Fixed= mcmcmc[[1]]$Fixed,      # All the same
        Random=mcmcmc[[1]]$Random,     # All the same
        Residual=mcmcmc[[1]]$Residual, # All the same
        family=mcmcmc[[1]]$family,     # All the same
        error.term=mcmcmc[[1]]$error.term,# All the same
        meta  =mcmcmc[[1]]$meta,       # All the same
        ginverse=mcmcmc[[1]]$ginverse, # All the same
        CP=    mcmcmc[[1]]$CP,         # All the same (NULL?)
        Lambda=mcmcmc[[1]]$Lambda,     # All the same (NULL?)
        XL=    mcmcmc[[1]]$XL,         # All the same (NULL?)
        y.additional=mcmcmc[[1]]$y.additional,# All the same
        Sol=do.call(mcmc.list, lapply(mcmcmc, function(m) m$Sol)),
        VCV=do.call(mcmc.list, lapply(mcmcmc, function(m) m$VCV)),
        DIC=do.call("c", lapply(mcmcmc, function(m) m$DIC))) # This will not behave as a regular MCMCglmm object, this is a vector of DICs
}

as.MCMCglmm <- function(myMCMCglmm) {
	ans <- myMCMCglmm
	ans$Sol <- do.call(rbind, myMCMCglmm$Sol)
	ans$VCV <- do.call(rbind, myMCMCglmm$VCV)
	ans$DIC <- mean(myMCMCglmm$DIC) # ???
	ans
}

posteriorPredictiveP <- function(object, data, summaryStat = function(pred, obs) { sum(((pred-obs)/pred)^2, na.rm=TRUE) }, pred.approx="taylor2", nsim=NULL ) {
	if (!("MCMCglmm" %in% class(object))) object <- as.MCMCglmm(object)
	
	pred <- predict.MCMCglmm(object, marginal=NULL, approx=pred.approx)
	obs <- c(data$Sdl, data$Mat)
	if (is.null(nsim)) nsim <- nrow(obj$Sol)
	sim <- simulate.MCMCglmm(object, nsim=nsim)
	
	sumobs <- summaryStat(pred, obs)
	sumsim <- apply(sim, 2, function(x) summaryStat(pred, x))
	mean(sumsim < sumobs)
}

loadORrun <- function(modname, MCMCargs, force.run=FALSE, cachedir="./cache", MCMCfun=my.MCMCglmm) {
	dir.create(cachedir, showWarnings = FALSE)
	ans <- try(readRDS(paste0(cachedir, "/", modname, ".rds")), silent=TRUE)
	if (!force.run && class(ans) == "try-error") {
		cat(ans, "\n")
		cat("Unable to find the model ", modname, " in cache. Running...\n")
		force.run <- TRUE
	}
	if (!force.run) {
		nitt <- (summary(ans$VCV)$end - summary(ans$VCV)$start)*summary(ans$VCV)$nchain
		if (nitt < 0.9*(MCMCargs$nitt - MCMCargs$burnin)) {
			cat("The number of iterations in the model stored in cache (", nitt, ") is less than expected (", MCMCargs$nitt - MCMCargs$burnin, "). Re-running...\n")
			force.run <- TRUE
		}
	}
	if (force.run) {
		ans <- do.call(MCMCfun, MCMCargs)
	}
	if (class(ans) != "try-error") {
		saveRDS(ans, file=paste0(cachedir, "/", modname, ".rds"), version=2)
	} else {
		stop("The MCMCglmm call did not succeed.")
	}
	ans
}

### Inferred selection responses
extractBLUP <- function(mcmclist, effect="animal") {
	# TODO: deal with "real" mcmc objects?
	cln <- colnames(mcmclist$Sol[[1]])
	traits <- cln[!grepl(".", cln, fixed=TRUE)]
	tmp <- lapply(mcmclist$Sol, function(mm) lapply(traits, function(trait) { mm[,grepl(paste0(trait, ".", effect, "."), colnames(mm), fixed=TRUE)]}))
	ans <- do.call(Map, c(tmp, f=rbind))
	names(ans) <- traits
	ans
}

envtab <- function(dd.env) {
	# dd.env is a dataframe with two columns, Sdl and Mat
	# very little to do here, but the logic of the program is the same as for gentab
	rownames(dd.env) <- sapply(strsplit(rownames(dd.env), split='\\.'), '[', 3)
	return(dd.env)
}

envBLUP <- function(model, blupFUN=mean) {
	envblup <- extractBLUP(model, "gen")
	envtab(data.frame(Sdl= apply(envblup$traitSdl, 2, blupFUN), Mat=apply(envblup$traitMat, 2, blupFUN)))
}

envMCMC <- function(model) {
	envblup <- extractBLUP(model, "gen")
	lapply(1:nrow(envblup[[1]]), function(i) envtab(data.frame(Sdl = envblup$traitSdl[i,], Mat=envblup$traitMat[i,])))
}

gentab <- function(data, gen, line) {
	# Generates a table with Sdl/Mat means per selected line (in columns) and generations (in line). 
	# Could be called on real or simulated data
	
	stopifnot(ncol(data) == 2)
	colnames(data) <- c("Sdl","Mat") # if the order is wrong, the rest is misleading. 
	
	sdl <- tapply(data[,"Sdl"], list(gen, line), mean)
	mat <- tapply(data[,"Mat"], list(gen, line), mean)
	sdl[c("F0","F1"),] <- sdl[c("F0","F1"),"O"]
	sdl <- sdl[,colnames(sdl) != "O"]
	colnames(sdl) <- paste0("Sdl.", colnames(sdl))
	mat[c("F0","F1"),] <- mat[c("F0","F1"),"O"]
	mat <- mat[,colnames(mat) != "O"]
	colnames(mat) <- paste0("Mat.", colnames(mat))
	return(cbind(sdl, mat))
}

genBLUP <- function(model, gen=NULL, line=NULL, blupFUN=mean) {
	genblup <- extractBLUP(model, "animal")
	blup.ind <- cbind(Sdl=apply(genblup[["traitSdl"]], 2, blupFUN), Mat=apply(genblup[["traitMat"]], 2, blupFUN))
	if (is.null(gen) || is.null(line)) return (blup.ind)

	return(gentab(blup.ind, gen, line))
}

genMCMC <- function(model, gen=NULL, line=NULL) {
	genblup <- extractBLUP(model, "animal")
	return(mclapply(1:nrow(genblup[["traitSdl"]]), function(i) gentab(cbind(Sdl=genblup[[1]][i,], Mat=genblup[["traitMat"]][i,]), gen, line), mc.cores=mc.cores))
}
```
