## Supplementary material for "Unidirectional response to bidirectional selection on body size II Quantitative genetics": Data files and analysis scripts: Dataclean.R

```
######### Tools to get a clean dataset from the original data file #####

# Note: it is important to make sure that all scripts call these
# functions, so that they are all based on the same dataset

datadir  <- "../data"
funcdir  <- "../R_functions"

source(paste(funcdir, "New2Old.R", sep="/"))


cleanSdl <- function(file="Sdl_Final_v3.txt", maxgen=7, P3only=TRUE) {
	sdl <- read.table(paste(datadir, file, sep="/"), header=TRUE)
	sdl <- sdl[!grepl(sdl$Family, pattern="Bonus"),]
	if (P3only) {
		sdl <- sdl[sdl$Phenotyping == "P3",]
	} else {
		sdl <- sdl[sdl$Phenotyping %in% paste0("P", 0:5),]
	}
	sdl <- sdl[sdl$Generation %in% paste0("F", 0:maxgen),]
	sdl$Generation <- factor(as.character(sdl$Generation), levels = paste0("F", 0:maxgen))
	sdl$Aqua <- factor(Aqua2Old(as.character(sdl$Family)))
	sdl$Family <- factor(Aqua2Old(as.character(sdl$Family), "fam"))
	sdl$Line <- factor(sdl$Line)
	sdl$Mat <- sdl$Sex != "I"
	
	sdl
}

cleanFam <- function(sdl) {
	
	famMeanSdl <- tapply(sdl$Sdl, INDEX=sdl$Family, FUN=mean, na.rm=TRUE)
	famMeanMat <- tapply(sdl$Mat, INDEX=sdl$Family, FUN=mean, na.rm=TRUE)
	genMeanSdl <- tapply(sdl$Sdl, INDEX=sdl$Generation, FUN=mean, na.rm=TRUE)
	genMeanMat <- tapply(sdl$Mat, INDEX=sdl$Generation, FUN=mean, na.rm=TRUE)
	linMeanSdl <- tapply(sdl$Sdl, INDEX=list(sdl$Generation, sdl$Line), FUN=mean, na.rm=TRUE)
	linMeanMat <- tapply(sdl$Mat, INDEX=list(sdl$Generation, sdl$Line), FUN=mean, na.rm=TRUE)
	
	fam <- data.frame(
	    MeanSdl = famMeanSdl,
	    MeanMat = famMeanMat,
	    VarSdl  = tapply(sdl$Sdl, INDEX=sdl$Family, FUN=var, na.rm=TRUE), 
	    Fratry = c(table(sdl$Family)))
	
	fam$Line <- sdl$Line[match(rownames(fam), as.character(sdl$Family))]
	fam$Gen  <- sdl$Generation[match(rownames(fam), as.character(sdl$Family))]
	    
	fam$GenSdl <- genMeanSdl[fam$Gen]
	fam$LinSdl <- linMeanSdl[cbind(fam$Gen, fam$Line)]
	
	fam$GenMat <- genMeanMat[fam$Gen]
	fam$LinMat <- linMeanMat[cbind(fam$Gen, fam$Line)]

	fam
}

cleanFry <- function(file="fry.txt") {
	rawfry <- read.table(paste(datadir, file, sep="/"), header=TRUE)

	fry <- data.frame(
		Family= factor(Aqua2Old(as.character(rawfry$Family))),
		Gen   = rawfry$Generation,
		Date  = as.Date(as.character(rawfry$Date), format="%d/%m/%Y"),
		Fry   = rawfry$NbFry)
	fry <- fry[order(as.character(fry$Family)),]
	row.names(fry) <- NULL
	fry
}

cleanEggs <- function(file="eggs.txt") {
	raweggs <- read.table(paste(datadir, file, sep="/"), header=TRUE)
	
	eggs <- data.frame(
		Family= factor(Aqua2Old(as.character(raweggs$Family))),
		Gen   = raweggs$Generation,
		Line  = raweggs$Line,
		Date  = as.Date(as.character(raweggs$Date), format="%d/%m/%Y"),
		Eggs  = raweggs$NbEggs)
	
	eggs
}
```
