## Supplementary material for "Unidirectional response to bidirectional selection on body size II Quantitative genetics": Data files and analysis scripts: Inbreeding.R

```
################################# Inbreeding.R ##########################
#
# Returns the inbreeding coefficients from a sdl file
# Several methods are proposed, they are supposed to return
# identical results. 
#
# Author: Arnaud Le Rouzic <>
# Version 1.0      2017-05-12
#
# Distributed under the WTFPL v2, no warranty
#
######################################################################

this.file.name <- function () # https://stackoverflow.com/a/1816487
{
    frame_files <- lapply(sys.frames(), function(x) x$ofile)
    frame_files <- Filter(Negate(is.null), frame_files)
    normalizePath(frame_files[[length(frame_files)]])
}

this.file <- this.file.name()
this.dir <- dirname(this.file)

source(paste(this.dir, "Sdl2Ped.R", sep="/"))
source(paste(this.dir, "New2Old.R", sep="/"))

Inbreeding <- function(sdl, method=c("kin", "kinb", "genlib")) {
    # Let's assume that the sdl is correct
    # (running SanityCheck can be costly)
    stopifnot(method[1] %in% c("kin", "kinb", "genlib"))

    # All individuals need a unique name
    sdl$ID <- as.character(sdl$ID)
    noID <- which(is.na(sdl$ID))
    sdl$ID[noID] <- paste0(
        sapply(strsplit(as.character(sdl$Family)[noID],split=".",fixed=TRUE), paste, collapse=""), 
        ifelse(is.na(sdl$Sex), "I", as.character(sdl$Sex))[noID], 101:(100+length(noID)), as.character(sdl$Line)[noID])
    
    if (method[1] == "kin") {
        return(Inbreeding.kin(sdl))
    } else if (method[1] == "kinb") {
        return(Inbreeding.kinb(sdl))
    } else if (method[1] == "genlib") {
        return(Inbreeding.genlib(sdl))
    } 
}

Inbreeding.kin <- function(sdl) {
    # this method consists in getting the kinship matrix with the kinship2
    # package, and compute inbreeding coefficients from the diagonal of the matrix
    
    stopifnot(require(kinship2))
        
    ped <- getped(sdl)
    kinmat <- kinship(ped)
    # if one's kinship coefficient with onself is k, 
    # the inbreeding coefficient should be 2*(k-1/2). 
    inb <- 2*diag(kinmat-0.5)
    fam <- as.character(sdl$Family[match(ped$full.id, sdl$ID)])
    inb <- inb[!duplicated(fam)]
    names(inb) <- unique(fam)
        
    return(inb[as.character(sdl$Family)])
}
    
Inbreeding.kinb <- function(sdl) {
    # this method consists in getting the kinship matrix with the kinship2
    # package, and compute inbreeding coefficients from the kinship of both parents
    # Note: a lot of copy-paste code from the Inbreeding.kin function
    
    stopifnot(require(kinship2))
    
    ped <- getped(sdl) 
    kinmat <- kinship(ped)
    colnames(kinmat) <- rownames(kinmat) <- names(ped$id)

    inb <- unlist(mapply(as.character(sdl$Father), as.character(sdl$Mother), 
        FUN=function(fa, mo) {
            if (fa %in% rownames(kinmat) && mo %in% colnames(kinmat) && !is.na(mo) && !is.na(fa)) {
                kinmat[fa, mo] 
            } else {
                0
            }}))

    fam <- as.character(sdl$Family[match(ped$full.id, sdl$ID)])
    inb <- inb[!duplicated(fam)]
    names(inb) <- unique(fam)
    
    return(inb[as.character(sdl$Family)])
}

Inbreeding.genlib <- function(sdl) {
    # this method is based on the GENLIB package

    stopifnot(require(GENLIB))
    
    ped <- getped(sdl) 
    
    pedgenlib <- as.data.frame(ped) 
    pedgenlib$sex <- as.numeric(pedgenlib$sex)
    pedgenlib$sex <- ifelse(pedgenlib$sex == 1, 1, 2) 
    # Warning: sex is wrong, but genlib does not accept anything but 1 and 2
    colnames(pedgenlib) <- c("ind", "father", "mother", "sex")
    glgen <- gen.genealogy(pedgenlib)
    inb <- gen.f(glgen, pedgenlib$ind)
    fam <- as.character(sdl$Family[match(ped$full.id, sdl$ID)])
    inb <- inb[!duplicated(fam)]
    names(inb) <- unique(fam)
    
    return(inb[as.character(sdl$Family)])
}
```
