## Supplementary material for "Unidirectional response to bidirectional selection on body size II Quantitative genetics": Data files and analysis scripts: New2Old.R

```
################################# New2Old.R ##########################
#
# Manipulates new and old aquarium/family numbers
#
# Author: Arnaud Le Rouzic <>
# Version 1.3      2017-05-31
#
# Distributed under the WTFPL v2, no warranty
#
######################################################################


##### Summary
######### check ID validity
#
# is.ValidAqua(x)
# is.ValidID(x)
#
######### new vs old
#
# is.NewAqua(x)
# is.OldAqua(x)
# is.NewID(x)
# is.OldID(x)
#
######### change old to new and vice versa
#
# Aqua2New(x)
# Aqua2Old(x)
# ID2New(x)
# ID2Old(x)
#
########## decomposition
#
# decompAqua(x)
# decompID(x)
# 
# which provide lists with various elements, such as:
# ..$Generation
# ..$Line
# ..$Family
# etc (check the code for details)


############################ Aquarium / Families ####################

 
decompOldAqua <- function(old.aqua) {
    if (is.na(old.aqua)) return (NA)
    stopifnot(is.OldAqua(old.aqua))
	rr <- regexec(pattern="^(F\\d)[.](\\d+)([A-Z])$", old.aqua)
	res <- regmatches(old.aqua, rr)[[1]]
    
    generation <- res[2]
    oldfam     <- as.numeric(res[3])
    oldrep     <- res[4]

    line       <- "P"
    if (oldfam > 20 && oldfam <= 40) line <- "G"
    if (oldfam > 40) line <- "A"
    newfam <- oldfam
    if (line=="G") newfam <- newfam - 20
    if (line=="A") newfam <- newfam - 40
	
    ans <- list(
        Generation = generation, 
        Family     = as.character(oldfam), 
        FamilyFull = paste0(generation, ".", as.character(oldfam)),
        FamilyNew  = paste0(newfam, line),
        FamilyOld  = as.character(oldfam), 
        Aqua       = paste0(oldfam, oldrep),
        AquaFull   = paste0(generation, ".", oldfam, oldrep),
        AquaNew    = paste0(newfam, line, tolower(oldrep)), 
        AquaOld    = paste0(oldfam, oldrep), 
        Line       = line)
    class(ans) <- c("aquaID", class(ans))
    ans
}

decompNewAqua <- function(new.aqua) {
    if (is.na(new.aqua)) return (NA)
    stopifnot(is.NewAqua(new.aqua))
	rr <- regexec(pattern="^(F\\d+)[.](\\d+)([PGA0])([a-z])$", new.aqua)
	res <- regmatches(new.aqua, rr)[[1]]
    
    generation <- res[2]
    newfam     <- paste0(res[3], res[4])
    newrep     <- res[5]
    line       <- res[4]
    
    oldfam <- as.numeric(res[3])
    if (line == "G") oldfam <- oldfam + 20
    if (line == "A") oldfam <- oldfam + 40   
     
	ans <- list(
        Generation = generation, 
        Family     = newfam, 
        FamilyFull = paste0(generation, ".", newfam),
        FamilyNew  = newfam,
        FamilyOld  = as.character(oldfam), 
        Aqua       = paste0(newfam, newrep),
        AquaFull   = paste0(generation, ".", newfam, newrep),
        AquaNew    = paste0(newfam, newrep), 
        AquaOld    = paste0(oldfam, toupper(newrep)),  
        Line       = line)
    class(ans) <- c("aquaID", class(ans))
    ans
}

is.NewAqua <- function(aqua) {
    ifelse (is.na(aqua), (NA), grepl(pattern="^F\\d+.\\d+[OAGP][a-z]$", aqua))
}  

is.OldAqua <- function(aqua) {
    ifelse (is.na(aqua), NA, grepl(pattern="^F\\d+.\\d+[A-Z]$", aqua))
}

is.ValidAqua <- function(aqua) {
    is.NewAqua(aqua) | is.OldAqua(aqua)
}  

decompAqua <- function(aqua) {
    ans <- lapply(aqua, function(aqua.u) {
        if (is.na(aqua.u)) return (NA)
        if (is.NewAqua(aqua.u)) decompNewAqua(aqua.u)
        else if (is.OldAqua(aqua.u)) decompOldAqua(aqua.u)
        else NA })
    if (length(ans) == 1) {ans[[1]]} else {ans}
}

old2newAqua.u <- function(old.u, result="aqua") {
    stopifnot(result %in% c("aqua", "fam"))
    if (is.na(old.u)) return (NA)
    if(!is.ValidAqua(old.u)) stop("Aquarium index ", old.u, " is not valid.")
    if (is.NewAqua(old.u)) return(old.u)
	aquaobj <- decompOldAqua(old.u)
    if (result=="aqua") {
        makenewAqua(aquaobj)
    } else if (result == "fam") {
        makenewFam(aquaobj)
    }
}

new2oldAqua.u <- function(new.u, result="aqua") {
    stopifnot(result %in% c("aqua", "fam"))
    if (is.na(new.u)) return (NA)
    if(!is.ValidAqua(new.u)) stop("Aquarium index ", new.u, " is not valid.")
    if (is.OldAqua(new.u)) return(new.u)
	aquaobj <- decompNewAqua(new.u)
	    if (result=="aqua") {
        makeoldAqua(aquaobj)
    } else if (result == "fam") {
        makeoldFam(aquaobj)
    }
}

Aqua2New <- function(oldaq, result="aqua") {
	sapply(oldaq, old2newAqua.u, result=result)
}

Aqua2Old <- function(newaq, result="aqua") {
	sapply(newaq, new2oldAqua.u, result=result)
}

#################################### Fish IDs ##########################

decompOldID <- function(old.id) {
    if (is.na(old.id)) return (NA)
    stopifnot(is.OldID(old.id))
	rr <- regexec(pattern="^(F\\d0?)([1-9]\\d*)([A-Z])([FMI])(\\d+)([OAGP])$", old.id)
	res <- regmatches(old.id, rr)[[1]]
    
    generation <- res[2]
    oldfam     <- as.numeric(res[3])
    oldrep     <- res[4]
    sex        <- res[5]
    index      <- res[6]
    line       <- res[7]
    newfam <- oldfam
    if (line=="G") newfam <- newfam - 20
    if (line=="A") newfam <- newfam - 40
	ans <- list(
        Generation = generation, 
        Family     = as.character(oldfam), 
        FamilyFull = paste0(generation, ".", as.character(oldfam)),    
        FamilyNew  = paste0(newfam, line),
        FamilyOld  = as.character(oldfam), 
        Aqua       = paste0(oldfam, oldrep),
        AquaFull   = paste0(generation, ".", oldfam, oldrep),                
        AquaNew    = paste0(newfam, line, tolower(oldrep)), 
        AquaOld    = paste0(oldfam, oldrep), 
        Sex        = sex, 
        BreederIndex=index, 
        Line       = line)
    class(ans) <- c("fishID", class(ans))
    ans
}

decompNewID <- function(new.id) {
    if (is.na(new.id)) return (NA)
    stopifnot(is.NewID(new.id))
	rr <- regexec(pattern="^(F\\d+)\\.(\\d+)([OAGP])([a-z])\\.([FMI])(\\d+)$", new.id)
	res <- regmatches(new.id, rr)[[1]]

    generation <- res[2]
    newfam     <- paste0(res[3], res[4])
    newrep     <- res[5]
    sex        <- res[6]
    index      <- res[7]
    line       <- res[4]
    
    oldfam <- as.numeric(res[3])
    if (line == "G") oldfam <- oldfam + 20
    if (line == "A") oldfam <- oldfam + 40   
	ans <- list(
        Generation = generation, 
        Family     = newfam, 
        FamilyFull = paste0(generation, ".", newfam),
        FamilyNew  = newfam,
        FamilyOld  = as.character(oldfam), 
        Aqua       = paste0(newfam, newrep),
        AquaFull   = paste0(generation, ".", newfam, newrep),        
        AquaNew    = paste0(newfam, newrep), 
        AquaOld    = paste0(oldfam, toupper(newrep)), 
        Sex        = sex, 
        BreederIndex=index, 
        Line       = line)
    class(ans) <- c("fishID", class(ans))
    ans
}

decompSdlLine <- function(sdl.line) {
    stopifnot(
        is.data.frame(sdl.line), 
        nrow(sdl.line==1), 
        all(c("Generation", "Line", "Family", "Sex") %in% colnames(sdl.line)))
    
    generation <- as.character(sdl.line$Generation)
    family <- as.character(sdl.line$Family)
    line <- as.character(sdl.line$Line)
    sex <- as.character(sdl.line$Sex)
    
    stopifnot(
        grepl(generation, pattern="^F\\d+$"),
        is.ValidAqua(family),
        grepl(sex, pattern="^[MFI]$"),
        grepl(line, pattern="^[OAGP]$"))
    
    dec <- decompAqua(family)
    
        
}

is.NewID <- function(id) {
    ifelse (is.na(id), NA, grepl(pattern="^F\\d+.\\d+[OAGP][abcd].[MFI]\\d+$", id))
}

is.OldID <- function(id) {
    ifelse (is.na(id), NA, grepl(pattern="^F\\d+[ABCD][MFI]\\d+[OAGP]$", id))
}

is.ValidID <- function(id) {
    is.NewID(id) | is.OldID(id)
}

decompID <- function(id) {
    ans <- lapply(id, function(id.u) {
        if (is.na(id.u)) return (NA)
        if (is.NewID(id.u)) decompNewID(id.u)
        else if (is.OldID(id.u)) decompOldID(id.u)
        else NA })
    if (length(ans) == 1) {ans[[1]]} else {ans}
}

old2newID.u <- function(old.u) {
    if (is.na(old.u)) return (NA)
    stopifnot(is.ValidID(old.u) || browser())
    if (is.NewID(old.u)) return(old.u)
	idobj <- decompOldID(old.u)
	makenewID(idobj)
}

new2oldID.u <- function(new.u) {
    if (is.na(new.u)) return (NA)
    stopifnot(is.ValidID(new.u))
    if (is.OldID(new.u)) return(new.u)
	idobj <- decompNewID(new.u)
	makeoldID(idobj)
}

ID2New <- function(oldid) {
	sapply(oldid, old2newID.u)
}

ID2Old <- function(newid) {
	sapply(newid, new2oldID.u)
}

###############################" Generate Tags and IDs ##################

# Generate IDs from various objects

makenewID <- function(idobj) UseMethod("makenewID")

makenewID.default <- function(idobj) {
    stop("Calling makenewID on a non-idobj object.")
}

makenewID.fishID <- function(idobj) {
	paste0(idobj$Generation, ".", idobj$AquaNew, ".", idobj$Sex, idobj$BreederIndex)
}

makenewID.aquaID <- function(idobj) {
	warning("Generating a random individual ID from a family ID, assuming immaturity.")
    paste0(idobj$Generation, ".", idobj$AquaNew, ".", "I", round(runif(10000,99999)))
}

makeoldID <- function(idobj) UseMethod("makeoldID")

makeoldID.default <- function(idobj) {
    browser()
    stop("Calling makeoldID on a non-idobj object.")
}

makeoldID.fishID <- function(idobj) {
	paste0(idobj$Generation, idobj$AquaOld, idobj$Sex, idobj$BreederIndex, idobj$Line)
}

makeoldID.aquaID <- function(idobj) {
	paste0(idobj$Generation, idobj$AquaOld, "I", round(runif(10000, 99999)), idpbj$Line)
}

# Generate Family tags

makenewFam <- function(idobj) UseMethod("makenewFam")

makenewFam.default <- function(idobj) {
    stop("Calling makenewFam on a non-idobj object.")
}

makenewFam.fishID <- function(idobj) {
    paste0(idobj$Generation, ".", idobj$FamilyNew)
}

makenewFam.aquaID <- function(idobj) {
    paste0(idobj$Generation, ".", idobj$FamilyNew)
}

makeoldFam <- function(idobj) UseMethod("makeoldFam")

makeoldFam.default <- function(idobj) {
    stop("Calling makeoldFam on a non-idobj object.")
}

makeoldFam.fishID <- function(idobj) {
    paste0(idobj$Generation, ".", idobj$FamilyOld)
}

makeoldFam.aquaID <- function(idobj) {
    paste0(idobj$Generation, ".", idobj$FamilyOld)
}

# Generate Aquarium Tags

makenewAqua <- function(idobj) UseMethod("makenewAqua")

makenewAqua.default <- function(idobj) {
    stop("Calling makenewAqua on a non-idobj object.")
}

makenewAqua.fishID <- function(idobj) {
    paste0(idobj$Generation, ".", idobj$AquaNew)
}

makenewAqua.aquaID <- function(idobj) {
    paste0(idobj$Generation, ".", idobj$AquaNew)
}

makeoldAqua <- function(idobj) UseMethod("makeoldAqua")

makeoldAqua.default <- function(idobj) {
    stop("Calling makeoldAqua on a non-idobj object.")
}

makeoldAqua.fishID <- function(idobj) {
    paste0(idobj$Generation, ".", idobj$AquaOld)
}

makeoldAqua.aquaID <- function(idobj) {
    paste0(idobj$Generation, ".", idobj$AquaOld)
}
```
