## Supplementary material for "Unidirectional response to bidirectional selection on body size II Quantitative genetics": Data files and analysis scripts: SanityCheck.R

```
################################# SanityCheck.R ##########################
#
# Checks whether the SDL data file is correct and consistent
#
# Author: Arnaud Le Rouzic <>
# Version 1.2      2017-02-28
#
# Distributed under the WTFPL v2, no warranty
#
######################################################################

this.file.name <- function () # https://stackoverflow.com/a/1816487
{
    frame_files <- lapply(sys.frames(), function(x) x$ofile)
    frame_files <- Filter(Negate(is.null), frame_files)
    normalizePath(frame_files[[length(frame_files)]])
}

this.file <- this.file.name()
this.dir <- dirname(this.file)

source(paste(this.dir, "New2Old.R", sep="/"))

col.names <- c("Generation", "Line", "Family", "Phenotyping", "Sdl", "Date", "Sex", "Straight", "ID", "Breeder", "Father", "Mother")

generation.label <- "F"
selected.lines <- c("O","A","G","P")
phenotyping.levels <- c(paste0("P", 0:7), "P1b")
sex.levels <- c("M","F","I")
straight.levels <- c("N", "T")
breeder.levels <- c("yes","no")

max.error.display <- 10

# All-in-one
SanityCheck <- function(sdl) {
	general.summary(sdl)
	test.factor.levels(sdl)
	test.pheno(sdl)
	test.pedigree(sdl)
}

is.a.number <- function(x) {
	return(!is.na(suppressWarnings(as.numeric(x))))
}

general.summary <- function(sdl) {
	# Prints a general summary of the sdl object (number of lines, columns, 
	# and counts per generation).
	cat("General summary\n")
	cat("\t", nrow(sdl), " lines and ", ncol(sdl), " columns\n", sep="")
	if (any(!(col.names %in% colnames(sdl))) || any(!(colnames(sdl) %in% col.names))) {
		cat("\tColumn names do not match the expected pattern:\n")
		if (any(!(col.names %in% colnames(sdl))))
            cat("\t\t", col.names[!(col.names %in% colnames(sdl))], "are missing.\n")
        if (any(!(colnames(sdl) %in% col.names)))
            cat("\t\t", colnames(sdl)[!(colnames(sdl) %in% col.names)], "are not necessary.\n")
	}
	
	tt <- table(sdl$Generation)
	cat("\tObservations per generation: \n")
	cat("\t", names(tt), "\n", sep="\t")
	cat("\t", tt, "\n", sep="\t")
}

test.factor.levels <- function(sdl) {
	# Checks the consistency of factor levels and missing data (NA) when required. 
	cat("Checking factor levels...\n")
	
	# Generation
	gener <- levels(sdl$Generation)
	invisible(sapply(gener, function(gg) {
		sp <- strsplit(gg, split="")[[1]]
		if (sp[1] != generation.label || !is.a.number(paste(sp[-1],collapse="")))
			cat("\tGeneration, factor level", gg, "incorrect\n")
		}))
	
	# Line
	lin <- levels(sdl$Line)
	if (!all(lin %in% selected.lines))
		cat("\tFactor levels ", selected.lines[!selected.lines %in% lin], "is incorrect\n")
	
	# Family	
	fam <- levels(sdl$Family)
	invisible(sapply(fam, function(ff) {
		if(!is.ValidAqua(ff)) {
			cat("\tFamily label", ff, "is incorrect\n")
		}}))
	na.fam <- is.na(sdl$Family)
	if (any(na.fam)) cat("Family:", sum(na.fam), " values are NA (lines:", which(na.fam)[1:min(sum(na.fam),max.error.display)], ")\n")

    wgf <- which(sapply(as.character(sdl$Family), function(ff) if(is.ValidAqua(ff)) decompAqua(ff)$Generation) != sdl$Generation)
    if (length(wgf) > 0) cat("\tFamily labels: ", unique(as.character(sdl$Family)[wgf]), " do not match generation number\n")
    
    wlf <- which(sapply(as.character(sdl$Family), function(ff) if(is.ValidAqua(ff)) decompAqua(ff)$Line) != sdl$Line & !(sdl$Generation %in% c("F0","F1")))
    if (length(wlf) > 0) cat("\tFamily labels: ", unique(as.character(sdl$Family)[wlf]), " do not match Line indicator\n")
	
	# Phenotyping	
	phen <- levels(sdl$Phenotyping)
	if(!all(phen %in% phenotyping.levels))
		cat("\tPhenotyping, incorrect factor levels:", phen[!phen %in% phenotyping.levels], "\n")
	na.phen <- is.na(sdl$Phenotyping)
	if (any(na.phen)) cat("Phenotyping:", sum(na.phen), "values are NA \n\t\t(lines:", which(na.phen)[1:min(sum(na.phen),max.error.display)], ")\n")
		
	# Sdl
	na.sdl <- is.na(sdl$Sdl)
	if (any(na.sdl)) cat("\tSdl:", sum(na.sdl), " values are NA \n\t\t(lines:", which(na.sdl)[1:min(sum(na.sdl),max.error.display)], ")\n")
	
	# Date
	na.date <- is.na(sdl$Date)
	no.date <- sapply(sdl$Date, function(aa) {
		dd <- try(as.Date(aa), silent=TRUE)
		if (class(dd) == "try-error") return(TRUE)
		return(FALSE)
	})
	if (any(na.date)) cat("\tDate:", sum(na.date), " values are NA \n\t\t(lines:", which(na.date)[1:min(sum(na.date),max.error.display)],")\n")
	if (any(no.date)) cat("\tDate:", sum(no.date), " values cannot be interpreted as dates \n\t\t(lines:", which(no.date)[1:min(sum(no.date),max.error.display)], ")\n")
	
	# Sex
	sx <- levels(sdl$Sex)
	if (!all(sx %in% sex.levels)) cat("\tSex: one of the factor levels", sx, "is incorrect\n")
	na.sex <- is.na(sdl$Sex)
	if (any(na.sex)) cat("\tSex:", sum(na.sex), "values are NA \n\t\t(lines: ", which(na.sex)[1:max.error.display], ")\n")
	
	# Straight
	na.str <- is.na(sdl$Straight)
	st <- levels(sdl$Straight)
	if (!all(st %in% straight.levels)) cat("\tStraight: one of the factor levels", st, "is incorrect\n")
		#no details, this situation might be normal (!?)
	if(any(na.str)) cat("\tStraight:", sum(na.str), "values are NA\n") 
	
	# ID
	valid.id <- is.ValidID(levels(sdl$ID))
	if (any(!valid.id)) cat("\tID: The following IDs are incorrect \n\t\t(", levels(sdl$ID)[!valid.id][1:min(sum(!valid.id), max.error.display)], ")\n")
	
	# Breeder
	na.bre <- is.na(sdl$Breeder)
	bre <- levels(sdl$Breeder)
	if (!all(bre %in% breeder.levels)) cat("\tBreeder: one of the factor levels", bre, "is incorrect\n")
	if (any(na.bre)) cat("\tBreeder:", sum(na.bre), "values are NA\n")
	
	# Father
	validF.id <- is.ValidID(levels(sdl$Father))
	if (any(!validF.id)) cat("\tFather: The following IDs are incorrect \n\t\t(", levels(sdl$Father)[!validF.id][1:min(sum(!validF.id), max.error.display)], ")\n")
	
	# Mother
	validM.id <- is.ValidID(levels(sdl$Mother))
	if (any(!validM.id)) cat("\tMother: The following IDs are incorrect \n\t\t(", levels(sdl$Mother)[!validM.id][1:min(sum(!validM.id), max.error.display)], ")\n")
		
}


test.pheno <- function(sdl) {
	# tests whether all individually-phenotyped individuals (P3) have a label and two parents
	test.indiv.P3 <- function(ll) {
		G0 <- "F0"
		if(is.na(sdl$ID[ll]) && !is.na(sdl$Breeder[ll]) && sdl$Breeder[ll] == breeder.levels[1]) 
			cat("\tLine", ll, ": P3-Phenotyped breeder with no ID\n")
		if(is.na(sdl$Father[ll]) && sdl$Generation[ll] != G0) 
			cat("\tLine", ll, ": P3-Phenotyped individual with no Father\n")
		if(is.na(sdl$Mother[ll]) && sdl$Generation[ll] != G0) 
			cat("\tLine", ll, ": P3-Phenotyped individual with no Mother\n")
		if(is.na(sdl$Sdl[ll])) 
			cat("\tLine", ll, ": P3-Phenotyped individual with no Sdl\n")
	}
	# 
	test.indiv.nonP3 <- function(ll) {
		G0 <- "F0"
		if(!is.na(sdl$ID[ll])) 
			cat("\tLine", ll, ":", as.character(sdl$Phenotyping[ll]), "-Phenotyped individual has an ID\n")
		if(is.na(sdl$Sdl[ll]))
			cat("\tLine", ll, ":", as.character(sdl$Phenotyping[ll]), "-Phenotyped individual with no Sdl\n")
		if(is.na(sdl$Father[ll]) && sdl$Generation[ll] != G0) 
			cat("\tLine", ll, ":", as.character(sdl$Phenotyping[ll]), "-Phenotyped individual with no Father\n")
		if(is.na(sdl$Mother[ll]) && sdl$Generation[ll] != G0) 
			cat("\tLine", ll, ":", as.character(sdl$Phenotyping[ll]), "-Phenotyped individual with no Mother\n")
	}
	cat("Testing the consistency of phenotyping data...\n")
	invisible(sapply(which(sdl$Phenotyping=="P3"), test.indiv.P3))
}

test.pedigree <- function(sdl) {
	# tests if the pedigree can be reconstructed. Even goes a bit further. 
	
	cat("Testing the consistency of pedigree information\n")
	
	# IDs are unique
	tt <- table(sdl$ID)
	if (any(tt>1)) cat("\tSome individual IDs are duplicated:", names(tt)[which(tt>1)],"\n")
	
	# ID information match Generation/Family/Sex
	ll <- which(!is.na(sdl$ID))
	invisible(sapply(ll, function(l) {
        id <- as.character(sdl$ID[l])
        if (!is.ValidID(id)) {
            cat("\tLine", l, ": Individual ID", id, "is not valid. Skipping the test.\n")
            return()
        }
        gn <- as.character(sdl$Generation[l])
        fm <- as.character(sdl$Family[l])
        sx <- as.character(sdl$Sex[l])
		dec <- decompID(id)
		if (dec$Generation != gn)
			cat("\tLine", l, ": Individual ID", id,"does not match its Generation level", gn,"\n")
		if (dec$AquaFull != fm) {browser()
			cat("\tLine", l, ": Individual ID", id,"does not match its Aquarium level", fm,"\n")}
		if (dec$Sex != sx)
			cat("\tLine", l, ": Individual ID", id,"does not match its Sex level", sx,"\n")			
		}))
		
	# All breeders (except F0) have 2 parents and these parents are present in the pedigree
	ll <- which(sdl$Breeder == breeder.levels[1])
	invisible(sapply(ll, function(l) {
		fath <- as.character(sdl$Father[l])
		moth <- as.character(sdl$Mother[l])
		id <- as.character(sdl$ID[l])
		if (is.na(id)) cat("\tIndividual line", l, "is a breeder but has no ID\n")
		if (sdl$Generation[l] != "F0" && is.na(fath)) cat("\tIndividual", id, "is a breeder but has no father\n")
		if (sdl$Generation[l] != "F0" && is.na(moth)) cat("\tIndividual", id, "is a breeder but has no mother\n")
        if(!is.na(fath) && is.ValidID(fath)) {
            if(length(which(names(tt) == fath))==0) cat("\tIndividual", id, "\'s father", fath, "cannot be found\n")
            if(decompID(fath)$Sex != sex.levels[1]) cat("\tIndividual", id, "\'s father", fath, "is not a male\n")
        }
        if(!is.na(moth) && is.ValidID(moth)) {
            if(length(which(names(tt) == moth))==0) cat("\tIndividual", id, "\'s mother", moth, "cannot be found\n")
            if(decompID(moth)$Sex != sex.levels[2]) cat("\tIndividual", id, "\'s mother", moth, "is not a female\n")
        }
		if (sdl$Generation[l] != "F0") {
            numgen <- function(x) suppressWarnings(as.numeric(substr(x, 2, nchar(x))))
			num.gener <- c(numgen(decompID(id)$Generation), numgen(decompID(fath)$Generation), numgen(decompID(moth)$Generation))
			if(!any(is.na(num.gener))) { 
				if (num.gener[2] != num.gener[1]-1) cat("\tIndividual", id,"\'s father", fath, "is not from the previous generation\n")
				if (num.gener[3] != num.gener[1]-1) cat("\tIndividual", id,"\'s mother", moth, "is not from the previous generation\n")
			}
		}
	}))
	
	# Each family has two unique parents
        # remove the aquarium character (aquarium number A, B... or a, b...) from the family names
  fam.only <- sapply(as.character(sdl$Family), function(ss) { if(is.ValidAqua(ss)) {decompAqua(ss)$FamilyFull} else {NA} })

        # checks if a family has more than one father/mother
  invisible(sapply(split(sdl, fam.only), function(subsdl) {
		if(length(unique(subsdl$Father)) > 1) cat("\tFamily", as.character(unique(subsdl$Family)), "has several fathers:", unique(as.character(subsdl$Father)),"\n")
		if(length(unique(subsdl$Mother)) > 1) cat("\tFamily", as.character(unique(subsdl$Family)), "has several mothers:", unique(as.character(subsdl$Mother)),"\n")
		}))
	
	invisible(sapply(split(data.frame(gen=sdl$Generation, uf=fam.only,pa=sdl$Father), sdl$Father), function(ff) {
		if(length(unique(ff$uf)) > 1) cat("\tFamilies", as.character(ff$gen[1]), as.character(unique(ff$uf)), "share the same father", as.character(ff$pa[1]), "\n") }))
	invisible(sapply(split(data.frame(gen=sdl$Generation, uf=fam.only,pa=sdl$Mother), sdl$Mother), function(ff) {
		if(length(unique(ff$uf)) > 1) cat("\tFamilies", as.character(ff$gen[1]), as.character(unique(ff$uf)), "share the same mother", as.character(ff$pa[1]), "\n") }))	
}
```
