## Supplementary material for "Unidirectional response to bidirectional selection on body size II Quantitative genetics": Data files and analysis scripts: Sdl2Ped.R

```
# Sdl2Ped: reconstructs the pedigree from the sdl file

whichid <- function(sdl, names, colname) {
	sapply(names, function(id) {w <- which(sdl$ID==id); as.character(sdl[[colname]][w])})
}

whichline <- function(sdl, names) {
	whichid(sdl, names, "Line")
}

whichgen <- function(sdl, names) {
	whichid(sdl, names, "Generation")
}

whichsdl <- function(sdl, names) {
	whichid(sdl, names, "Sdl")
}

whichcol <- function(sdl, names, cols=c(O="black",G="blue",P="red",A="gray")) {
	cols[whichline(sdl, names)]
}

whichcolnum <- function(sdl, names, cols=c(O="black",G="blue",P="red",A="gray")) {
	tmp <- 1:length(cols)
	names(tmp) <- cols
	tmp[whichcol(sdl, names, cols)]
}

sdl2ped <- function(sdl, output="") {
	# We assume that the sdl is clean!
	
	first <- TRUE
	invisible(sapply(which(!is.na(sdl$ID)), function(ll) {
		cat(file=output, as.character(sdl$ID[ll]), "\t", as.character(sdl$Father[ll]), "\t", as.character(sdl$Mother[ll]), "\n", append=!first)
		first <<- FALSE }))
}

getped <- function(sdl, fast=TRUE) {
    numgen <- function(IDs) as.numeric(sapply(IDs, function(id) if(is.na(id)) NA else decompID(id)$Generation))
    sex    <- function(IDs) sapply(IDs, function(id) if(is.na(id)) NA else decompID(id)$Sex)
    numsex <- function(IDs) {sx <- sex(IDs); ifelse(is.na(sx), 3, ifelse(sx == "M", 1, ifelse(sx=="F", 2, 3))) }
	
	stopifnot(require("kinship2"))
    
    sdl$ID     <- as.character(sdl$ID)
    sdl$Family <- as.character(sdl$Family)
    sdl$Father <- as.character(sdl$Father)
    sdl$Mother <- as.character(sdl$Mother)
    
    sdlfast <- sdl
    if (fast) { # trims the pedigree for duplicated / unnecessary individuals
        isparent <- sdl$ID %in% c(sdl$Mother, sdl$Father)
        famparent <- sdl[isparent, "Family"]
        sdlfast <- sdl[c(which(isparent), match( unique(sdl$Family)[!(unique(sdl$Family) %in% unique(famparent))], sdl$Family)),]
    }
    
    no.id <- is.na(sdlfast$ID) | duplicated(sdlfast$ID)
    if (sum(no.id) > 0) 
        warning ("getped: some individuals have no or duplicated IDs => excluded from the pedigree")

	ID.name <- sdlfast$ID[!no.id]
        
    unknown.parents <- c(sdlfast$Father[!sdlfast$Father %in% ID.name], sdlfast$Mother[!sdlfast$Mother %in% ID.name])
    unknown.parents <- unknown.parents[!is.na(unknown.parents)]
    if (length(unknown.parents) > 0)
        warning("getped: some parents (", paste(unique(unknown.parents), collapse=", ") , ") were absent from the data set => added in the pedigree.")
    ID.name <- c(unknown.parents, ID.name)
	ID.index <- 1:length(ID.name)
    
    if (any(!(is.na(ID.name) | is.ValidID(ID.name)))) 
        stop("Invalid IDs in the pedigree: ", paste(ID.name[!is.ValidID(ID.name)][1:5], collapse=", "))

    names(ID.index) <- ID.name

	Father.index <- ID.index[sdlfast$Father[match(ID.name, sdlfast$ID)]]
	Father.index[is.na(Father.index)] <- 0
	Mother.index <-ID.index[sdlfast$Mother[match(ID.name, sdlfast$ID)]]
	Mother.index[is.na(Mother.index)] <- 0

	ped <- pedigree(id=ID.index, dadid=Father.index, momid=Mother.index, sex=numsex(ID.name))
    ped$full.id <- ID.name
    ped
}

getped.glmm <- function(sdl) {
    # Returns a pedigree in the format requested by MCMCglmm
    ped <- getped(sdl, FALSE)
    ans <- data.frame(animal=ped$id, sire=ped$mindex, dam=ped$findex)
    ans$sire[ans$sire==0] <- NA
    ans$dam[ans$dam==0]   <- NA
    ans
}

plotped <- function(sdl, cols=c(O="black",G="blue",P="red",A="gray")) {
	if(!require("kinship2")) stop("Library kinship2 is necessary")
	Allped <- getped(sdl)
	plot(Allped, col=sapply(names(Allped$id), function(id) {w <- which(sdl$ID==id); cols[as.character(sdl$Line[w])]}))
}
```
