## Supplementary material for "Unidirectional response to bidirectional selection on body size II Quantitative genetics": Data files and analysis scripts: SelectionResponse.R

```
# Functions to study the selection response from the sdl file

focal.phen <- "P3"
low.line <- "P"
control.line <- "A"
high.line <- "G"

start.generation <- c("F0","F1")
breeder.levels <- c("yes","no")

plotEvolution <- function(sdl, phen=sdl$Sdl, reference="none", breeders=TRUE, error="se", ylab="Sdl (mm)", ylim=NULL, col=c("blue","gray","red"), lty=rep(1,3), pch=rep(1,3), shift=0, breeders.pch=20, ...) {
	# Plots the response to selection from the sdl data
	# phen: vector of phenotypes to plot (default is simply Sdl)
	# reference: can be "none", "control", "grandmean", "sym"
	# breeders: if TRUE, symbols are displayed for the breeder means
	# error: can be "se" (standard errors), "sd" (standard deviations), or NULL
	# shift: shifts the time series (useful for error="sd" to avoid overlap)
	# breeders.pch: symbol for breeders
	# other parameters and ...: options for plot
	
	sdl <- sdl[sdl$Phenotyping==focal.phen,]
	
	means <- arrayFUN(sdl=sdl, phen=phen, mean, na.rm=TRUE)
	sds <-  arrayFUN(sdl=sdl, phen=phen, sd, na.rm=TRUE)
	N <- arrayFUN(sdl=sdl, phen=phen, length)
	
	if (reference=="none") correction <- rep(0,length(means$grand))
	else if (reference=="control") correction <- means$line[,control.line]
	else if (reference=="grandmean") correction <- c(means$grand)
	else if (reference=="sym") correction <- 0.5*means$line[,high.line]+0.5*means$line[,low.line]
	else stop("Unknown value for reference:", reference)

	if (is.null(ylim)) {
		ylim <- range(means$line - correction)
		ylim <- ylim + c(-1,1)*diff(ylim)*if(breeders || error=="sd") 1 else 0.5
	}
	if (error == "se") {
		sds$line <- sds$line / sqrt(N$line)
	}
	
	plot(NULL,xlab="Generations", ylab=ylab, xlim=c(1,length(means$grand)+3*shift), ylim=ylim, xaxt="n", ...)
	axis(1, at=1:length(means$grand), labels=rownames(means$line))
	for (ii in c(1,3,2)) {
		if (!is.null(error))
			arrows(x0=1:length(means$grand)+shift*(ii-1), y0=means$line[,ii]-correction-sds$line[,ii], y1=means$line[,ii]-correction+sds$line[,ii], length=0, col=col[ii], lty=lty[ii])

		lines(1:length(means$grand)+shift*(ii-1), means$line[,ii]-correction, lty=lty[ii], col=col[ii], pch=pch[ii], type="b")
		if (breeders) {
			points(1:length(means$grand)+shift*(ii-1), means$breed[,ii]-correction, pch=breeders.pch, col=col[ii])
		}
	}
}

plotResponse <- function(sdl, phen=sdl$Sdl, reference="none", xlab="Cumulative differential (mm)", ylab="Cumulative response (mm)", ylim=NULL, col=c("blue","red"), from="F1", ...) {
	# Plots the realized response vs. selection differential
	# sdl: data set
	# phen: optional, vector of phenotypes (same size as sdl) if the phenotype is not sdl$Sdl
	# reference: one among "none","control","grandmean","sym"
	# from: generation considered as the starting point of the selection response
	# other parameters and ... : parameters to plot
	
	sdl <- sdl[sdl$Phenotyping==focal.phen,]
	phen <- phen[sdl$Phenotyping==focal.phen]
	
	means <- arrayFUN(sdl=sdl, phen=phen, mean, na.rm=TRUE)
	cumdiff <- means$breed - means$line
	cumdiff <- apply(cumdiff, 2, function(x) cumsum(x) - sum(x[1:which(rownames(cumdiff)==from)]))
	
	if (reference=="none") correction <- means$line[from,]
	else if (reference=="control") correction <- means$line[,control.line]
	else if (reference=="grandmean") correction <- c(means$grand)
	else if (reference=="sym") correction <- 0.5*means$line[,high.line]+0.5*means$line[,low.line]
	else stop("Unknown value for reference:", reference)
	
	resp <- means$line - correction
	
	gen <- rownames(resp)
	gen <- gen[which(gen==from):length(gen)]
	
	
	rangeresp <- range(resp[,colnames(resp) != control.line])
	plot(NULL, xlim=range(cumdiff), ylim=rangeresp+0.2*c(-1,1)*diff(rangeresp), xlab=xlab, ylab=ylab, ...)
	lines(cumdiff[gen,low.line],resp[gen,low.line],col=col[1], type="b")
	lines(cumdiff[gen,high.line],resp[gen,high.line],col=col[2], type="b")
	
	seldiff <- c(cumdiff[gen,high.line], cumdiff[gen,low.line])
	selresp <- c(resp[gen,high.line], resp[gen,low.line])
	reg <- lm(selresp ~ 0 + seldiff)
	
	abline(reg, lty=2)
	
	legend("topleft", lty=NULL, legend=paste0("Realized h2 +- 95%CI = ", format(coef(reg)[1], digits=3), " (", format(confint(reg)[1,1], digits=3), ";", format(confint(reg)[1,2], digits=3), ")"))
	text(seldiff, selresp, rep(gen, 2), pos=ifelse(selresp > 0, 3, 1))
}

plotParentOffspring <- function(sdl, phen=sdl$Sdl, sex.off="both", sex.par="both", col=c("blue","gray","red"), gen=NULL, xlim=NULL, ylim=NULL, xlab=NULL, ylab=NULL, reg.all=TRUE, reg.line=TRUE, ...) {
	# Plots the mid-parent - offspring relationship and estimates the heritability
	# sdl: data set
	# phen: vector of phenotypes if different from sdl$Sdl
	# sex.off: can be "both", "male", or "female"
	# sex.par: can be "both", "male", or "female"
	# gen: generation of offpring to be plotted (NULL: all). 
	# col: colors for down, control, and up lines
	# reg.all: whether the parent - offspring regression should be displayed
	# ref.line: whether the parent - offspring regression should be displayed for each line separately
	# other parameters: parameters for the plot() function
	sdl <- sdl[sdl$Phenotyping==focal.phen,]
	phen <- phen[sdl$Phenotyping==focal.phen]

	# Useless except to get generation/line names
	means <- arrayFUN(sdl=sdl, phen=phen, mean, na.rm=TRUE)
	
	names(col) <- colnames(means$line)
	pch <- 1:nrow(means$line)
	names(pch) <- rownames(means$line)
	if (is.null(gen)) gen <- rownames(means$line)[-1]
	
	ans <- list()
	
	invisible(sapply(which(sdl$Generation %in% gen & !is.na(sdl$Father) & !is.na(sdl$Mother)), function(ll) {
		# ll is the line in sdl
		gn <- as.character(sdl$Generation[ll])
		lin <- as.character(sdl$Line[ll])
		if (lin=="O") lin <- control.line # does not matter as all lines are the same for F0 and F1
		offsp <- phen[ll]
		mpar <- switch(sex.par, 
			both=0.5*phenindiv(sdl, phen, as.character(sdl$Father[ll])) + 0.5*phenindiv(sdl, phen, as.character(sdl$Mother[ll])), 
			male=phenindiv(sdl, phen, as.character(sdl$Father[ll])), 
			female=phenindiv(sdl, phen, as.character(sdl$Mother[ll])))
		tag <- paste(gn,lin,sep="-")
		if (!is.na(mpar)) { # this can happen because of errors in the sdl file
			if (sex.off == "both" 
			|| (sex.off == "male" && sdl$Sex[ll]=="M") 
			|| (sex.off == "female" && sdl$Sex[ll]=="F"))
				ans[[tag]] <<- rbind(ans[[tag]], c(offsp,mpar))
		}
	}))
	
	# Normalization
	ans <- lapply(ans, function(x) t(t(x)-apply(x,2,mean)))
	
	full.dat <- do.call(rbind, ans)
	
	if (is.null(ylim)) {
		ylim <- range(full.dat[,1])
	}
	if (is.null(xlim)) {
		xlim <- range(full.dat[,2])
	}
	if (is.null(xlab)) {
		xlab <- switch(sex.par, both="Mean parent", male="Father", female="Mother")
	}
	if (is.null(ylab)) {
		ylab <- switch(sex.off, both="Offspring", male="Son", female="Daughter")
	}
		
	plot(NULL, xlim=xlim, ylim=ylim, xlab=xlab, ylab=ylab, ...)
	for (i in 1:length(ans)) {
		tag <- names(ans)[i]
		ss <- strsplit(tag, split="-")[[1]]
		gn <- ss[1]
		lin <- ss[2]
		points(ans[[tag]][,2], ans[[tag]][,1], col=col[lin], pch=pch[gn])
	}

	capt <- list()
	
	ret <- list()
	
	if(reg.all) {
		full.dat <- do.call(rbind, ans)
		reg <- lm(full.dat[,1]~full.dat[,2])
		abline(reg)
		ret[["full"]] <- reg
		capt$legend <- c(capt$legend, paste0("h2 +- 95%CI = ", displ(coef(reg)[2]), " (", displ(confint(reg)[2,1]), " ; ", displ(confint(reg)[2,2]), ")"))
		capt$col <- c(capt$col, 1)
		capt$pch <- c(capt$pch, 1)
		capt$lty <- c(capt$lty, 1)
	}
	
	for (ll in 1:length(col)) {
		capt$col <- c(capt$col, col[ll])
		capt$pch <- c(capt$pch, 16)
		capt$lty <- c(capt$lty, if(reg.line) ll else NA)
		txt <- paste0("Line ", names(col)[ll])
		if (reg.line) {
			tags <- names(ans)[sapply(names(ans), function(nn) strsplit(nn, split="-")[[1]][2]==names(col)[ll])]
			dat <- do.call(rbind, ans[tags])
			reg <- lm(dat[,1]~dat[,2])
			ret[[names(col)[ll]]] <- reg
			abline(reg, col=col[ll], lty=capt$lty[ll])
			txt <- paste0(txt, " h2 +- 95%CI = ", displ(coef(reg)[2]), " (", displ(confint(reg)[2,1]), " ; ", displ(confint(reg)[2,2]), ")")
		}
		capt$legend <- c(capt$legend, txt)
	}

	legend("topleft", legend=capt$legend, lty=capt$lty, pch=capt$pch, col=capt$col)
	invisible(ret)
}


plotSRA <- function(sdl, phen=sdl$Sdl, reference="none", exclude.gen="F0", sraFUN=sraCstvar, ...) {
	# Runs a Selection Response Analysis via the sra package
	# sdl: dataset
	# phen: phenotype if different from Sdl$Sdl
	# reference: one among "none", "control", "grandmean", or "sym"
	# exclude gen: by default, exclude the first generation
	# sraFUN: for advanced users only, change the sra function
	# ...: additional parameters to the sra function
	if(!require("sra")) stop("The package sra is necessary to run this analysis")
	
	sdl <- sdl[sdl$Phenotyping==focal.phen,]
	phen <- phen[sdl$Phenotyping==focal.phen]

	means <- arrayFUN(sdl=sdl, phen=phen, mean, na.rm=TRUE)	
	vars <- arrayFUN(sdl=sdl, phen=phen, var, na.rm=TRUE)
	N <- arrayFUN(sdl=sdl, phen=phen, length)

	if (reference=="none") correction <- rep(0,length(means$grand))
	else if (reference=="control") correction <- means$line[,control.line]
	else if (reference=="grandmean") correction <- c(means$grand)
	else if (reference=="sym") correction <- 0.5*means$line[,high.line]+0.5*means$line[,low.line]
	else stop("Unknown value for reference:", reference)

	phen.mean <- c((means$line-correction)[!rownames(means$line) %in% exclude.gen])
	phen.var <- c(vars$line[!rownames(vars$line) %in% exclude.gen])
	phen.sel <- c((means$breed-correction)[!rownames(means$breed) %in% exclude.gen])
	var.sel <- c(vars$breed[!rownames(vars$breed) %in% exclude.gen])	
	n <- c(N$line[!rownames(N$line) %in% exclude.gen])
	gen <- rep(1:(nrow(means$line)-length(exclude.gen)), ncol(means$line))
	rep <- rep(colnames(means$line), each=nrow(means$line)-length(exclude.gen))

	sraD <- sraData(phen.mean=phen.mean, phen.var=phen.var, phen.sel=phen.sel, var.sel=var.sel, N=n, gen=gen, rep=rep)
	
	sraC <- sraFUN(sraD, ...)
	layout(1:2)
	plot(sraC)
	plot(sraC, variance=TRUE, legend=FALSE)
	layout(1)
}

arrayFUN <- function(sdl, phen=sdl$Sdl, FUN=mean, ...) {
	ans <- list()
	ans$grand <- grandFUN(sdl=sdl, phen=phen, FUN=FUN, ...)
	ans$line <- lineFUN(sdl=sdl, phen=phen, FUN=FUN, ...)
	breeders <- sdl$Breeder==breeder.levels[1]
	ans$breed <- lineFUN(sdl[breeders,], phen=phen[breeders], FUN=FUN, ...)
	# Small hacks due to how data is coded
	ans$line[start.generation,] <- ans$grand[start.generation]
	ans$breed[start.generation[1],] <- 
		lineFUN(sdl[breeders,], phen=phen[breeders], lines="O", FUN=FUN, ...)[start.generation[1],]
	
	return(ans)
}

grandFUN <- function(sdl, phen=sdl$Sdl, FUN=mean, ...) {
	return(tapply(phen, sdl$Generation, FUN, ...))
}

lineFUN <- function(sdl, phen=sdl$Sdl, lines=c(low.line, control.line, high.line), FUN=mean, ...) {
	return(sapply(lines, function(ll) tapply(phen[sdl$Line==ll], sdl$Generation[sdl$Line==ll], FUN, ...)))
}

grandmean <- function(sdl, phen=sdl$Sdl) {
	grandFUN(sdl=sdl, phen=phen, FUN=mean, na.rm=TRUE)
}

linemean <- function(sdl, phen=sdl$Sdl, lines=c(low.line, control.line, high.line)) {
	lineFUN(sdl=sdl, phen=phen, lines=lines, FUN=mean, na.rm=TRUE)
}

breedmean <- function(sdl, phen=sdl$Sdl, lines=c(low.line, control.line, high.line)) {
	breeders <- sdl$Breeder==breeder.levels[1]
	lineFUN(sdl[breeders,], phen=phen[breeders], lines=lines, FUN=mean, na.rm=TRUE)
}

phenindiv <- function(sdl, phen=sdl$Sdl, indivID) {
	indivID <- as.character(indivID)
	ll <- which(sdl$ID==indivID)
	if (length(ll) == 0) {
		warning("Individual ", indivID, " cannot be found.")
		return(NA)
	}
	if (length(ll) > 1) {
		warning("Individual ", indivID, " was found several times.")
		return(NA)
	}
	return(phen[ll])
}

parentgener <- function(gener, allowneg=FALSE) {
	gener <- as.character(gener)
	if (substr(gener,1,1) != "F") stop(gener, " is not a proper generation tag.")
	gnum <- as.numeric(substr(gener, 2, nchar(gener)))
	if (is.na(gnum)) stop(gener, " is not a proper generation tag.")
	if (!allowneg && gnum < 1) return(NA)
	return(paste0(substr(gener,1,1), gnum-1))
}

displ <- function(number) {
	format(number, digits=3)
}
```
