## Supplementary material for "Unidirectional response to bidirectional selection on body size II Quantitative genetics": Data files and analysis scripts: simbidi.R

```
library(MASS)    # for mvrnorm

logit <- function(p) log(p/(1-p))
unlogit <- function(x) 1/(1+exp(-x))

global.ID <- 0
getID <- function() { global.ID <<- global.ID+1; global.ID }

assign.fitness <- function(population, sel.p) {
    stopifnot(length(population) >= 2)
    sexlim <- round(length(population)/2)
    bin.trait <- sapply(population, function(i) i$phenotype[2])
    bin.trait.males <- bin.trait[1:sexlim]
    bin.trait.females <- bin.trait[(sexlim+1):length(bin.trait)]
    quant.trait <- sapply(population, function(i) i$phenotype[1])
    quant.trait.males <- quant.trait[1:sexlim]
    quant.trait.females <- quant.trait[(sexlim+1):length(quant.trait)]
    
    # By convention, males are the first half of the population, females the other half. The
    # threshold applies independently in males and females. 
    
    stopifnot(sum(bin.trait.males) > 0, sum(bin.trait.females) > 0)
    
    thresh.males <- quantile(quant.trait.males[bin.trait.males==1], probs=if (sel.p >= 0) 1-sel.p else -sel.p)
    fitness.males <- ifelse(bin.trait.males==1 & if (sel.p > 0) quant.trait.males > thresh.males else quant.trait.males < thresh.males, 1, 0)
    
    thresh.females <- quantile(quant.trait.females[bin.trait.females==1], probs=if (sel.p >= 0) 1-sel.p else -sel.p)
    fitness.females <- ifelse(bin.trait.females==1 & if (sel.p > 0) quant.trait.females > thresh.females else quant.trait.females < thresh.females, 1, 0)    
    
    fitness <- c(fitness.males, fitness.females)
    return(lapply(seq_along(fitness), function(ni) { indiv <- population[[ni]]; indiv$fitness <- fitness[ni]; indiv }))
}

genoI2R <- function(genoI, param) {
    # Computes the correlated genotypes from uncorrelated normalized genotypes
    R <- param$G.eigvec
    #R <- t(t(param$G.eigvec)*sign(colSums(param$G.eigvec)))
    S <- diag(sqrt(param$G.eigval))
    c(R %*% S %*% genoI)
}

genoR2I <- function(genoR, param) {
    R <- param$G.eigvec
    S<- diag(sqrt(param$G.eigval))
    c(solve(R %*% S) %*% genoR)
}

genoI2R.mat <- function(genoI, Gmat) {
    egg <- eigen(Gmat)
    fake.param <- list(G.eigvec = egg$vectors, G.eigval=egg$values)
    genoI2R(genoI, fake.param)
}

genoR2I.mat <- function(genoR, Gmat) {
    egg <- eigen(Gmat)
    fake.param <- list(G.eigvec = egg$vectors, G.eigval=egg$values)
    genoR2I(genoR, fake.param)   
}

genoR2phen <- function(genoR, param) {
    # Draws a vector of phenotype from correlated genotypes
    phen <- mvrnorm(1, genoR, Sigma=param$Emat)
    phen[2] <- rbinom(1, 1, prob=unlogit(phen[2]))
    return(phen)
}

bidi.mate  <- function(father, mother, param) {
    offspring <- list(genotypeI=
        c(0.5*father$genotypeI[1]+0.5*mother$genotypeI[1] + rnorm(1, sd=sqrt(1/2)), 
          0.5*father$genotypeI[2]+0.5*mother$genotypeI[2] + rnorm(1, sd=sqrt(1/2))))
    offspring$genotypeR <- genoI2R(offspring$genotypeI, param)
    offspring$phenotype <- genoR2phen(offspring$genotypeR, param)
    offspring$ID <- getID()
    offspring$father <- father$ID
    offspring$mother <- mother$ID
    return(offspring)
}

bidi.reproduce <- function(population, param) {
    # By convention, males are the first half of the population, females the other half.
    fitness <- sapply(population, function(i) i$fitness)
    fitness.males <- fitness[1:round(length(fitness)/2)]
    fitness.females <- fitness[round(length(fitness)/2+1):length(fitness)]
    newpop <- lapply(1:length(population), function(pp) bidi.mate(population[[sample(seq_along(fitness.males), 1, prob=fitness.males)]], population[[sample(round(length(fitness)/2)+seq_along(fitness.females), 1, prob=fitness.females)]], param=param))
    newpop <- assign.fitness(newpop, param$sel.p)
    return(newpop)
}

outputpop <- function(pop, summary=TRUE, gen=NULL) {
    if (summary) 
        return (summarypop(pop, gen=gen))
    else
        return (detailedpop(pop, gen=gen))
}

summarypop <- function(pop, gen=NULL, meanphen=TRUE, fitness=TRUE) {
    ans <- if(is.null(gen)) NULL else c(gen=gen)
    phen.quant <- sapply(pop, function(i) i$phenotype[1])
    phen.bin <- sapply(pop, function(i) i$phenotype[2])    
    if (meanphen) {
        ans <- c(ans, m.phen1=mean(phen.quant), m.phen2=mean(phen.bin), cor=cor(phen.quant, phen.bin))
    }
    if (fitness) {
        fit <- sapply(pop, function(i) i$fitness)
        fit <- fit/mean(fit)
        reg <- lm(fit ~ phen.quant + phen.bin)
        ans <- c(ans, beta1=coef(reg)[2], beta2=coef(reg)[3])
    } 
    return(ans)
}

detailedpop <- function(pop, gen=gen) {
    ans <- data.frame(
        gen       = if(is.null(gen)) NA else gen,
        ID        = sapply(pop, function(i) i$ID), 
        phen.quant= sapply(pop, function(i) i$phenotype[1]), 
        phen.bin  = sapply(pop, function(i) i$phenotype[2]),
        gen.quant = sapply(pop, function(i) i$genotypeR[1]),
        gen.bin   = sapply(pop, function(i) i$genotypeR[2]),
        fitness   = sapply(pop, function(i) i$fitness),
        mother    = sapply(pop, function(i) i$mother), 
        father    = sapply(pop, function(i) i$father))
    ans
}

bidi.simrun <- function(population = NULL, g0=c(0, 2), Emat=diag(2), Gmat=diag(2), Gen=10, sel.p=0.2, N=100, summary=TRUE) {
    param <- list(Emat=Emat, Gmat=Gmat, Gen=Gen, sel.p=sel.p, N=N)
    param$G.eigvec <- eigen(param$Gmat)$vectors
    param$G.eigval <- eigen(param$Gmat)$values
    
    if (is.null(population)) {
        population <- lapply(1:param$N, function(n) {
            indiv <- list(genotypeI = rnorm(length(g0), mean=genoR2I(g0, param)))
            indiv$genotypeR <- genoI2R(indiv$genotypeI, param)
            indiv$phenotype <- genoR2phen(indiv$genotypeR, param)
            indiv$ID <- getID()
            indiv$father <- indiv$mother <- NA
            return(indiv)
        })
    }
    population <- assign.fitness(population, param$sel.p)
    ans <- NULL
    ans <- outputpop(population, summary, gen=0)
    for (gg in 1:Gen) {
        population <- bidi.reproduce(population, param)
        ans <- rbind(ans, outputpop(population, summary, gen=gg))
    }
    ans
}
```
