## Supplementary material for "Unidirectional response to bidirectional selection on body size II Quantitative genetics": Data files and analysis scripts: sra-bivar.R

```
library("stats4")

sraData.bivar <- function(
    phen.X.mean, phen.X.var, phen.Y.mean, phen.Y.var, phen.XY.cov, 
    sel.X.mean, sel.X.var, sel.Y.mean, sel.Y.var, 
    N, gen, rep) 
{
    # So far, nothing fancy: just put the data in a data.frame
  ans <- data.frame(
    phen.X.mean=phen.X.mean, phen.X.var=phen.X.var, 
    phen.Y.mean=phen.Y.mean, phen.Y.var=phen.Y.var, 
    phen.XY.cov=phen.XY.cov,
    sel.X.mean=sel.X.mean, sel.Y.mean=sel.Y.mean, 
    sel.X.var=sel.X.var, sel.Y.var=sel.Y.var,
    N=N, gen=gen, rep=rep)
    
  class(ans) <- c("sradata", class(ans))
  return(ans)
}

logit <- function(p) log(p/(1-p))
unlogit <- function(u) 1/(1+exp(-u))
logit2 <- function(r) log((1+r)/(1-r))
unlogit2 <- function(u) (exp(u)-1)/(exp(u)+1)

sraStartingvalues <- function(parameter, sradata, rand=0)
	# Provides stupid, educated, or random guess about starting values. This is a mess, should be cleaned up
	# at one point. 
{
    .sraStartingvalues.mu0 <- function(sradata, colnm="mean")
        { return(mean(sradata[sradata[,"gen"]==1,colnm]) + rand*rnorm(1,0,abs(mean(sradata[sradata[,"gen"]==1,colnm])))) }
    .sraStartingvalues.logvarA0 <- function(sradata, colnm="var")
        { return(log(0.2*mean(sradata[,colnm]*exp(rand*rnorm(1,0,0.2*mean(sradata[,colnm])))))) }
    .sraStartingvalues.logvarE0 <- function(sradata, colnm="var")
        { return(log(0.8*mean(sradata[,colnm]*exp(rand*rnorm(1,0,0.8*mean(sradata[,colnm])))))) }
    
    if (parameter == "mu0")
        { return(.sraStartingvalues.mu0(sradata, "mean")) }
    if (parameter == "mu0.A")
		{ return(.sraStartingvalues.mu0(sradata, "phen.X.mean")) }
    if (parameter == "mu0.B")
		{ return(.sraStartingvalues.mu0(sradata, "phen.Y.mean")) }		
    if (parameter == "logvarA0")
        { return(.sraStartingvalues.logvarA0(sradata)) } 
    if (parameter == "logvarA0.pos")
        { return(.sraStartingvalues.logvarA0(sradata)) } 
    if (parameter == "logvarA0.neg")
        { return(.sraStartingvalues.logvarA0(sradata)) }                 
    if (parameter == "logvarA0.A")
        { return(.sraStartingvalues.logvarA0(sradata, "phen.X.var")) }
    if (parameter == "logvarA0.A.pos")
        { return(.sraStartingvalues.logvarA0(sradata, "phen.X.var")) }
    if (parameter == "logvarA0.A.neg")
        { return(.sraStartingvalues.logvarA0(sradata, "phen.X.var")) }        
    if (parameter == "logvarA0.B")
        { return(.sraStartingvalues.logvarA0(sradata, "phen.Y.var")) }                 
    if (parameter == "logvarE0")
        { return(.sraStartingvalues.logvarE0(sradata)) }
    if (parameter == "logvarE0.A")
        { return(.sraStartingvalues.logvarE0(sradata, "phen.X.var")) }
    if (parameter == "logvarE0.B")
        { return(.sraStartingvalues.logvarE0(sradata, "phen.Y.var")) }                
    if (parameter == "logit2rA0.AB") 
		{ return(0) }
    if (parameter == "logit2rE0.AB") 
		{ return(0) }      	
    if (parameter == "logvarME")
        { return(log(exp(.sraStartingvalues.logvarE0(sradata))/2)) } 
    if (parameter == "logIA0")
        { return(log(exp(.sraStartingvalues.logvarA0(sradata)) / (.sraStartingvalues.mu0(sradata)^2)))}
    if (parameter == "logIE0")
        { return(log(exp(.sraStartingvalues.logvarE0(sradata)) / (.sraStartingvalues.mu0(sradata)^2)))}       
    if (parameter == "logith20")
        { a <- exp(.sraStartingvalues.logvarA0(sradata)); e <- exp(.sraStartingvalues.logvarE0(sradata));
          h2 <- a/(a+e); return(log(h2/(1-h2))) }       
    if (parameter == "logvarP0")
        { return(log(exp(.sraStartingvalues.logvarA0(sradata))+exp(.sraStartingvalues.logvarE0(sradata)))) }
    if (parameter == "o")
        { return(.sraStartingvalues.mu0(sradata)) }
    if (parameter == "s")
        { return(1 + rand*rnorm(1,0,1)) }
    if (parameter == "logepsilon")
        { return(-10 + rand*rnorm(1,0,10)) }
    if (parameter == "logminusepsilon")
        { return(-10 + rand*rnorm(1,0,10)) }
    if (parameter == "logvarepsilon")
        { return(0 + rand*rnorm(1,0,1)) }
    if (parameter == "kc")
        { return(0 + rand*rnorm(1,0,1)) }
    if (parameter == "kg")
        { return(0 + rand*rnorm(1,0,1)) }
    if (parameter == "logvarM")
        { return(-20 + rand*rnorm(1,0,10)) }
    if (parameter == "logNe")
        { return(log(100) + rand*rnorm(1,0,2)) }
    if (parameter == "relativekA0")
        { return(0 + rand*rnorm(1,0,0.5)) }
    if (parameter == "relativekE0")
        { return(0 + rand*rnorm(1,0,0.5)) }
    if (parameter == "kA1")
        { return(1 + rand*rnorm(1,0,0.5)) }
    if (parameter == "kE1")
        { return(1 + rand*rnorm(1,0,0.5)) }
    if (parameter == "kA2")
        { return(0 + rand*rnorm(1,0,0.5)) } 
    if (parameter == "kE2")
        { return(0 + rand*rnorm(1,0,0.5)) } 
    if (parameter == "kA3")
        { return(0 + rand*rnorm(1,0,0.5)) } 
    if (parameter == "kE3")
        { return(0 + rand*rnorm(1,0,0.5)) } 
    if (parameter == "logrelativekA0")
        { return(-1 + rand*rnorm(1,0,1)) }  
    if (parameter == "logrelativekE0")
        { return(-1 + rand*rnorm(1,0,1)) }  
    if (parameter == "logkA1")
        { return(0 + rand*rnorm(1,0,1)) }   
    if (parameter == "logkE1")
        { return(0 + rand*rnorm(1,0,1)) }   
    if (parameter == "logkA2")
        { return(-3 + rand*rnorm(1,0,2)) }
    if (parameter == "logkE2")
        { return(-3 + rand*rnorm(1,0,2)) }  
    if (parameter == "logkA3")
        { return(-3 + rand*rnorm(1,0,2)) }  
    if (parameter == "logkE3")
        { return(-3 + rand*rnorm(1,0,2)) }      
    stop("Unknown parameter ", parameter, ".")
}


sraAutoregMinuslogL.bivar <- function(
    X.mean, X.var, Y.mean, Y.var, XY.cov, data.N, 
    theor.X.mean, theor.X.var, theor.Y.mean, theor.Y.var, theor.XY.cov)
	# This is the core likelihood function. it computes the likelihood of the bivariate normal
	# distribution. Parameters are the observations (means, variances, covariance) and the
	# theoretical corresponding values
{
    minuslogL <- 0

    # TODO: check if the data provided is consistent (vectors of the same length etc). 
    #if ((length(data.mean) != length(data.var)) | (length(data.mean) != 
    #length(data.N)) | (length(data.mean) != length(theor.mean)) | 
    #(length(data.mean) != length(theor.var))) {
    #stop("vectors have different lengths")
    #}

    for (t in 1:(length(theor.X.mean))) {
        if (is.infinite(theor.X.var[t]) || is.nan(theor.X.var[t]) || (theor.X.var[t] < 0) ||
            is.infinite(theor.Y.var[t]) || is.nan(theor.Y.var[t]) || (theor.Y.var[t] < 0) ||
            is.infinite(theor.XY.cov[t]) || is.nan(theor.XY.cov[t])) {
                return(Inf)
            }
            
        rho <- theor.XY.cov[t]/(sqrt(theor.X.var[t])*sqrt(theor.Y.var[t]))
        
        if ((rho < -1) || (rho > 1))
        { 
            return(Inf)
        }
        
        K <- -data.N[t]*log(2*pi*sqrt(theor.X.var[t]*theor.Y.var[t]*(1-rho^2)))

        minuslogL <- minuslogL - K + data.N[t]*((X.var[t]+(X.mean[t]-theor.X.mean[t])^2)/theor.X.var[t] + (Y.var[t]+(Y.mean[t]-theor.Y.mean[t])^2)/theor.Y.var[t] - 2*rho*(XY.cov[t]+X.mean[t]*Y.mean[t] - theor.X.mean[t]*Y.mean[t] - theor.Y.mean[t]*X.mean[t]+theor.X.mean[t]*theor.Y.mean[t])/(sqrt(theor.X.var[t]*theor.Y.var[t])))/(2*(1-rho^2))
    }
    return(minuslogL)          
}


sraMinuslogL.bivar <- function(sradata, FUNtimeseries=sraAutoregTimeseries.bivar, Bulmer=TRUE, ...)
{
	# Computes the likelihood of a full data set (all generations, several repetitions, etc). 
	# This generic function does not need to know about the model parameters, they are passed through the
	# ... argument. 

    ss <- split(sradata, sradata$rep)
    minuslogL <- 0
    for(pop in ss)
    {
        range <- 1:(nrow(pop)-1)
        detP <- pop$phen.X.var[range] * pop$phen.Y.var[range] - (pop$phen.XY.cov[range])^2
        s.A <- pop$sel.X.mean[range] - pop$phen.X.mean[range]
        s.B <- pop$sel.Y.mean[range] - pop$phen.Y.mean[range]
        beta.A <- (s.A*pop$phen.Y.var[range] - s.B * pop$phen.XY.cov[range]) / detP
        beta.B <- (s.B*pop$phen.X.var[range] - s.A * pop$phen.XY.cov[range]) / detP
        delta.A <- (pop$sel.X.var[range] - pop$phen.X.var[range])/pop$phen.X.var[range]
        delta.B <- (pop$sel.Y.var[range] - pop$phen.Y.var[range])/pop$phen.Y.var[range]
                
        tsr <- do.call(what=FUNtimeseries, args=c(list(
            beta.A=beta.A, beta.B=beta.B, delta.A=delta.A, delta.B=delta.B,
            ...)))
        minuslogL <- minuslogL + sraAutoregMinuslogL.bivar(
			X.mean=pop$phen.X.mean, X.var=pop$phen.X.var, 
			Y.mean=pop$phen.Y.mean, Y.var=pop$phen.Y.var, 
			XY.cov=pop$phen.XY.cov, 
			data.N=pop$N, 
			theor.X.mean=tsr$mean.A, theor.Y.mean=tsr$mean.B, 
			theor.X.var=tsr$varP.A, theor.Y.var=tsr$varP.B, 
			theor.XY.cov=tsr$covarP)
    }
    return(minuslogL)       
}


sraTimeseries.bivar <- function(
    beta.A, beta.B, delta.A = rep(0, length(beta.A)), delta.B = rep(0, length(beta.B)), 
	mu0.A=0, mu0.B=0, logvarA0.A=0, logvarA0.B=0, 
	logvarE0.A=0, logvarE0.B=0, logit2rA0.AB=0, logit2rE0.AB=0, logNe=log(100))
	# Produces a theoretical time series from a set of parameters, and selection strengths 
    # (selection on the means --beta-- and the variances --delta--)
{
    ans <- list()
    
    Ne <- exp(logNe)

    mu.A <- mu0.A
    mu.B <- mu0.B
    d.A <- 0
    d.B <- 0
    vara.A <- exp(logvarA0.A) 
    vara.B <- exp(logvarA0.B)
    varA.A <- vara.A + d.A
    varA.B <- vara.B + d.B
    varE.A <- exp(logvarE0.A)
    varE.B <- exp(logvarE0.B) 
    rA <- (exp(logit2rA0.AB)-1)/(exp(logit2rA0.AB)+1)
    rE <- (exp(logit2rE0.AB)-1)/(exp(logit2rE0.AB)+1)
    covarA <- rA*sqrt(varA.A)*sqrt(varA.B)
    covarE <- rE*sqrt(varE.A)*sqrt(varE.B)
    
    for (t in 1:(length(beta.A)))
    {
        # Lande's equation
        mu.A <- c(mu.A, mu.A[t] + varA.A[t] * beta.A[t] + covarA[t]*beta.B[t])
        mu.B <- c(mu.B, mu.B[t] + varA.B[t] * beta.B[t] + covarA[t]*beta.A[t])
    
		vara.A.tp1 <- vara.A[t] * (1 - 1/(2*Ne))
		vara.B.tp1 <- vara.B[t] * (1 - 1/(2*Ne))
        d.A.tp1 <- 0.5*(1-1/Ne)*(d.A[t]+delta.A[t]*(varA.A[t]**2)/(varA.A[t]+varE.A[t]))
        d.B.tp1 <- 0.5*(1-1/Ne)*(d.B[t]+delta.B[t]*(varA.B[t]**2)/(varA.B[t]+varE.B[t]))
        
        varA.A.tp1 <- vara.A.tp1 + d.A.tp1
        varA.B.tp1 <- vara.B.tp1 + d.B.tp1
        varE.A.tp1 <- varE.A[t]
        varE.B.tp1 <- varE.B[t]
        covarA.tp1 <- covarA[t]*(1 - 1/(2*Ne))
        covarE.tp1 <- covarE[t]
                
        if (is.nan(varA.A.tp1) || is.infinite(varA.A.tp1)) 
            {varA.A.tp1 <- 0}
        if (is.nan(varA.B.tp1) || is.infinite(varA.B.tp1)) 
            {varA.B.tp1 <- 0}            
        if (varA.A.tp1 < 0)
            {varA.A.tp1 <- 0}
        if (varA.B.tp1 < 0)
            {varA.B.tp1 <- 0}            
        if (is.nan(varE.A.tp1) || is.infinite(varE.A.tp1))
            {varE.A.tp1 <- 0} 
        if (is.nan(varE.B.tp1) || is.infinite(varE.B.tp1))
            {varE.B.tp1 <- 0}             
        if (varE.A.tp1 < 0)
            {varE.A.tp1 <- 0 }    
        if (varE.B.tp1 < 0)
            {varE.B.tp1 <- 0 } 
        
        vara.A <- c(vara.A, vara.A.tp1)
        vara.B <- c(vara.B, vara.B.tp1)
        d.A <- c(d.A, d.A.tp1)
        d.B <- c(d.B, d.B.tp1)
        
        varA.A <- c(varA.A, varA.A.tp1)
        varA.B <- c(varA.B, varA.B.tp1)
        
        covarA <- c(covarA, covarA.tp1)
        covarE <- c(covarE, covarE.tp1)
        
        varE.A <- c(varE.A, varE.A.tp1)
        varE.B <- c(varE.B, varE.B.tp1)
    }
    
    ans$mean.A <- mu.A
    ans$mean.B <- mu.B
    ans$varA.A <- varA.A
    ans$varA.B <- varA.B
    ans$covarA <- covarA
    ans$covarE <- covarE
    ans$varE.A <- varE.A
    ans$varE.B <- varE.B
    
    ans$varP.A <- ans$varA.A + ans$varE.A
    ans$varP.B <- ans$varA.B + ans$varE.B
    ans$covarP <- ans$covarA + ans$covarE
    return(ans)     
}


sraCstvar.bivar <-function (sradata, start = NULL, fixed = NULL, 
          Bulmer = TRUE, ...) 
{
  if (!Bulmer) {
    sradata$sel.X.var <- sradata$phen.X.var
    sradata$sel.Y.var <- sradata$phen.Y.var
  }
  default.start <- list(mu0.A = NA, mu0.B=NA, logvarA0.A = NA, logvarA0.B = NA, logvarE0.A = NA, logvarE0.B = NA, logit2rA0.AB = NA, logit2rE0.AB = NA)
  default.fixed <- list(logNe = log(1e+10))

  default.start[names(fixed)] <- NULL
  default.start[names(start)] <- start
  default.fixed[names(start)] <- NULL
  default.fixed[names(fixed)] <- fixed
  start <- default.start
  fixed <- default.fixed
  start[is.na(start)] <- sapply(names(start[is.na(start)]), 
                                sraStartingvalues, sradata = sradata)
  mlewrapper <- function(mu0.A, mu0.B, logvarA0.A, logvarA0.B, logvarE0.A, logvarE0.B, logit2rA0.AB, logit2rE0.AB, logNe) {
    sraMinuslogL.bivar(sradata = sradata, FUNtimeseries = sraTimeseries.bivar, 
                 mu0.A = mu0.A, mu0.B = mu0.B, logvarA0.A = logvarA0.A, logvarA0.B = logvarA0.B, 
                 logvarE0.A = logvarE0.A, logvarE0.B = logvarE0.B, logit2rA0.AB = logit2rA0.AB, logit2rE0.AB = logit2rE0.AB,
                 logNe = logNe)
  }
  fit <- mle(minuslogl = mlewrapper, start = start, fixed = fixed, 
             ...)
  return(sraMakeObject.bivar(sradata = sradata, model = fit, start = start, 
                       fixed = fixed, FUNtimeseries = sraTimeseries.bivar))
}


#### AICc for SRA (bivar or univar)
AICc.SRA<-function(SRAobj){
  LL<-logLik(SRAobj)[1]
  Nob<-nrow(SRAobj$data)-2 # number of contrasts taken as N  (if two lines starting form the same P population)
  K<-length(SRAobj$coefficients)
  AIC <- -2*LL + 2*K
  AICc<- AIC + 2*K*(K+1)/(Nob-K-1) 
return(AICc)
}


sraMakeObject.bivar <- function(sradata, model, start, fixed, FUNtimeseries)
{
	# Produces a summary object containing the data and the model fit. This is enough for
	# plotting, analysis, etc. 
    ans <- list()
    ans$data <- sradata
    ans$model <- model
    ans$start <- start
    ans$coefficients <- coef(model)[names(start)]
    # Confidence intervals: approximated from standard errors
    ans$confint <- cbind(summary(ans$model)@coef[,1]-2*summary(ans$model)@coef[,2], 
            summary(ans$model)@coef[,1]+2*summary(ans$model)@coef[,2])  
    ans$pred <- list()
    ans$residuals <- list()
    ans$vresiduals <- list()
    for(pop in split(ans$data, ans$data$rep))
    {
        range <- 1:(nrow(pop)-1)
        detP <- pop$phen.X.var[range] * pop$phen.Y.var[range] - (pop$phen.XY.cov[range])^2
        s.A <- pop$sel.X.mean[range] - pop$phen.X.mean[range]
        s.B <- pop$sel.Y.mean[range] - pop$phen.Y.mean[range]
        beta.A <- (s.A*pop$phen.Y.var[range] - s.B * pop$phen.XY.cov[range]) / detP
        beta.B <- (s.B*pop$phen.X.var[range] - s.A * pop$phen.XY.cov[range]) / detP
        delta.A <- (pop$sel.X.var[range] - pop$phen.X.var[range])/pop$phen.X.var[range]
        delta.B <- (pop$sel.Y.var[range] - pop$phen.Y.var[range])/pop$phen.Y.var[range]
               
        cc <- c(list(beta.A=beta.A, beta.B=beta.B, delta.A=delta.A, delta.B=delta.B))
        cc <- c(cc, ans$coefficients, unlist(fixed))
        ts <- do.call(what=FUNtimeseries, args=cc)
            
        r <- pop$rep[1] # All should be the same 
        ans$pred[[as.character(r)]] <- list()
        ans$pred[[as.character(r)]]$Gen <- names(ts$mean.A)
        ans$pred[[as.character(r)]]$phen.A <- ts$mean.A
        ans$pred[[as.character(r)]]$beta.A <- beta.A
        ans$pred[[as.character(r)]]$phen.B <- ts$mean.B
        ans$pred[[as.character(r)]]$beta.B <- beta.B        
        ans$pred[[as.character(r)]]$varP.A <- ts$varP.A
        ans$pred[[as.character(r)]]$varP.B <- ts$varP.B
        ans$pred[[as.character(r)]]$covarP <- ts$covarP
        ans$pred[[as.character(r)]]$varA.A <- ts$varA.A
        ans$pred[[as.character(r)]]$varA.B <- ts$varA.B
        ans$pred[[as.character(r)]]$covarA <- ts$covarA
        
#~         ans$residuals[[as.character(r)]] <- ans$data[ans$data[,"rep"]==as.character(r),"mean"] - ts$mean
#~         ans$vresiduals[[as.character(r)]] <- ans$data[ans$data[,"rep"]==as.character(r),"var"] - ts$varP
    }
    class(ans) <- c("srafit", "list")
    return(ans)
}


.expandandname <- function(x, nm) {
    stopifnot(length(nm) >= length(x))
    x <- rep(x, length.out=length(nm))
    if (is.null(names(x))) {
        names(x) <- nm
    } else {
        stopifnot(all(names(x) %in% nm))
    }
    x
}

plot.srafit.bivar.mean <- function(x, y=NULL, trait1="Selected trait", trait2="Correlated trait", col="black", pch=1, lty.data=1, lty.model=2, xlab="Generations", ...) {
    stopifnot("srafit" %in% class(x))
    
    nbr.ts <- length(x$pred)
    
    range1 <- range(c(x$data$phen.X.mean, unlist(lapply(x$pred, "[[", "phen.A"))))
    range2 <- range(c(x$data$phen.Y.mean, unlist(lapply(x$pred, "[[", "phen.B"))))
    
    rangegen <- range(unique(x$data$gen))
    
    col <- .expandandname(col, names(x$pred))
    pch <- .expandandname(pch, names(x$pred))
    lty.data <- .expandandname(lty.data, names(x$pred))
    lty.model<- .expandandname(lty.model, names(x$pred))
    
    layout(1:2)
    plot(NULL, xlim=rangegen, ylim=range1, xlab=xlab, ylab=trait1, ...)
    invisible(lapply(names(x$pred), function(i) 
        lines(x$pred[[i]]$phen.A, type="l", lty=lty.model[i], col=col[i])))
    invisible(lapply(names(x$pred), function(i) 
        lines(x$data$phen.X.mean[x$data$rep==i], type="b", lty=lty.data[i], col=col[i], pch=pch[i])))
    
    plot(NULL, xlim=rangegen, ylim=range2, xlab=xlab, ylab=trait2, ...)
    invisible(lapply(names(x$pred), function(i) 
        lines(x$pred[[i]]$phen.B, type="l", lty=lty.model[i], col=col[i])))
    invisible(lapply(names(x$pred), function(i) 
        lines(x$data$phen.Y.mean[x$data$rep==i], type="b", lty=lty.data[i], col=col[i], pch=pch[i])))
    
    layout(1)
}

plot.srafit.bivar.var <- function(x, y=NULL, trait1="Selected trait", trait2="Correlated trait", col="black", pch=1, lty.data=1, lty.model=2, xlab="Generations", ...) {
    stopifnot("srafit" %in% class(x))
    
    nbr.ts <- length(x$pred)
    
    range1 <- c(0, max(c(x$data$phen.X.var, unlist(lapply(x$pred, "[[", "varP.A")))))
    range2 <- c(0, max(c(x$data$phen.Y.var, unlist(lapply(x$pred, "[[", "varP.B")))))
    range3 <- range(c(0, x$data$phen.XY.cov, unlist(lapply(x$pred, "[[", "covarP")), unlist(lapply(x$pred, "[[", "covarA"))))
    
    rangegen <- range(unique(x$data$gen))
    
    col <- .expandandname(col, names(x$pred))
    pch <- .expandandname(pch, names(x$pred))
    lty.data <- .expandandname(lty.data, names(x$pred))
    lty.model<- .expandandname(lty.model, names(x$pred))
    
    layout(1:3)
    plot(NULL, xlim=rangegen, ylim=range1, xlab=xlab, ylab=paste0("Var ", trait1), ...)
    invisible(lapply(names(x$pred), function(i) 
        lines(x$pred[[i]]$varP.A, type="l", lty=lty.model[i], col=col[i])))
    invisible(lapply(names(x$pred), function(i) 
        lines(x$pred[[i]]$varA.A, type="l", lty=3, col=col[i])))    
    invisible(lapply(names(x$pred), function(i) 
        lines(x$data$phen.X.var[x$data$rep==i], type="b", 
            lty=lty.data[i], col=col[i], pch=pch[i])))
    
    plot(NULL, xlim=rangegen, ylim=range2, xlab=xlab, ylab=paste0("Var ", trait2), ...)
    invisible(lapply(names(x$pred), function(i) 
        lines(x$pred[[i]]$varP.B, type="l", lty=lty.model[i], col=col[i])))
    invisible(lapply(names(x$pred), function(i) 
        lines(x$pred[[i]]$varA.B, type="l", lty=3, col=col[i])))      
    invisible(lapply(names(x$pred), function(i) 
        lines(x$data$phen.Y.var[x$data$rep==i], type="b", lty=lty.data[i], col=col[i], pch=pch[i])))
    
    plot(NULL, xlim=rangegen, ylim=range3, xlab=xlab, ylab=paste0("Cov (", trait1, ", ", trait2, ")"), ...)
    invisible(lapply(names(x$pred), function(i) 
        lines(x$pred[[i]]$covarP, type="l", lty=lty.model[i], col=col[i])))
    invisible(lapply(names(x$pred), function(i) 
        lines(x$pred[[i]]$covarA, type="l", lty=3, col=col[i])))
    invisible(lapply(names(x$pred), function(i) 
        lines(x$data$phen.XY.cov[x$data$rep==i], type="b", lty=lty.data[i], col=col[i], pch=pch[i])))
    
    
    
    layout(1)
}

plot.srafit.bivar.2Dgrad <- function(x, y=NULL, trait1="Selected trait", trait2="Correlated trait", col="black", lty.grad=2, lty.response=1, lty.predict=3, response.exaggerate=5, ...) {

    nbr.gen <- max(x$data$gen)    
    nbr.ts <- length(x$pred)
    
    # Average responses to selection, estimated by linear regression on the raw data
    avgrespx <- setNames(sapply(names(x$pred), function(i) {
        yy <- x$data$phen.X.mean[x$data$rep==i] - x$data$phen.X.mean[x$data$rep==i & x$data$gen==1]
        xx <- x$data$gen[x$data$rep==i]
        coef(lm(yy ~ 0 + xx))[1]}), names(x$pred))
    avgrespy <- setNames(sapply(names(x$pred), function(i) {
        yy <- x$data$phen.Y.mean[x$data$rep==i] - x$data$phen.Y.mean[x$data$rep==i & x$data$gen==1]
        xx <- x$data$gen[x$data$rep==i]
        coef(lm(yy ~ 0 + xx))[1]}), names(x$pred))
        
    # Cumulative gradients
    cumgradx <- setNames(lapply(x$pred, function(ss) cumsum(ss$beta.A*ss$varP.A[-nbr.gen])), names(x$pred))
    cumgrady <- setNames(lapply(x$pred, function(ss) cumsum(ss$beta.B*ss$varP.B[-nbr.gen])), names(x$pred))
    
    rangex <- range(c(0, unlist(cumgradx), response.exaggerate*nbr.gen*avgrespx))
    rangey <- range(c(0, unlist(cumgrady), response.exaggerate*nbr.gen*avgrespy))    

    col          <- .expandandname(col, names(x$pred))
    lty.grad     <- .expandandname(lty.grad, names(x$pred))
    lty.response <- .expandandname(lty.response, names(x$pred))
    lty.predict  <- .expandandname(lty.predict, names(x$pred))
    
    # Bivariate gradients and selection response
    plot(NULL, xlim=rangex, ylim=rangey, xlab=trait1, ylab=trait2, ...)
    invisible(lapply(names(x$pred), function(i) 
        lines(x=c(0, cumgradx[[i]]), y=c(0, cumgrady[[i]]), lty=lty.grad[i], col=col[i])))
    invisible(lapply(names(x$pred), function(i) 
        arrows(x0=0, y0=0, 
            x1=response.exaggerate*sum(diff(x$pred[[i]]$phen.A)), 
            y1=response.exaggerate*sum(diff(x$pred[[i]]$phen.B)), col=col[i], lty=lty.predict[i], length=0.1)))
    invisible(lapply(names(x$pred), function(i) 
        arrows(x0=0, y0=0, 
            x1=response.exaggerate*nbr.gen*avgrespx[i], 
            y1=response.exaggerate*nbr.gen*avgrespy[i], col=col[i], lty=lty.response[i], length=0.1)))        
}

plot.srafit.bivar.grad <- function(x, y=NULL, trait1="Selected trait", trait2="Correlated trait", col="black", pch=1, lty.data=1, lty.model=2, xlab="Generations", ...) {
    stopifnot("srafit" %in% class(x))
    
    nbr.ts <- length(x$pred)
    nbr.gen <- max(x$data$gen)
    
    range1 <- range(c(0, unlist(lapply(x$pred, "[[", "beta.A"))))
    range2 <- range(c(0, unlist(lapply(x$pred, "[[", "beta.B"))))

    
    rangegen <- range(unique(x$data$gen))
    
    col <- .expandandname(col, names(x$pred))
    pch <- .expandandname(pch, names(x$pred))
    lty.data <- .expandandname(lty.data, names(x$pred))
    lty.model<- .expandandname(lty.model, names(x$pred))
    
    layout(1:2)
    #Gradient trait 1
    plot(NULL, xlim=rangegen, ylim=range1, xlab=xlab, ylab=paste0("Gradient ", trait1), ...)
    invisible(lapply(names(x$pred), function(i) 
        lines(x$pred[[i]]$beta.A, type="b", lty=lty.data[i], pch=pch[i], col=col[i])))
    abline(h=0, lty=3)
    
    # Gradient trait 2
    plot(NULL, xlim=rangegen, ylim=range2, xlab=xlab, ylab=paste0("Gradient ", trait2), ...)
    invisible(lapply(names(x$pred), function(i) 
        lines(x$pred[[i]]$beta.B, type="b", lty=lty.data[i], pch=pch[i], col=col[i])))
    abline(h=0, lty=3)
    
    layout(1)
}


######################################################################################################
plot.sra.biv<-function(SRAbivar, SelTrait="Selected trait", CorTrait="Correlated trait", 
                        Main="Selection response",    
                        col.pred="blue", col.obs="black", Main.Size=1,
                        control=NA, col.cont="grey"
                        ) {
  
  Ngen<-max(SRAbivar$data$gen)
  ## Plot trait A (selected)
SEa <- sqrt(SRAbivar$data$phen.X.var/SRAbivar$data$N)
SEaTop<-  SRAbivar$data$phen.X.mean +SEa
SEaBot<-  SRAbivar$data$phen.X.mean -SEa
  rangeYA<-range(c(SRAbivar$data$phen.X.mean, SRAbivar$pred$D$phen.A, SRAbivar$pred$U$phen.A, SEaTop, SEaBot))
  plot(SRAbivar$data$gen, SRAbivar$data$phen.X.mean, type="p", pch=16, xaxp=c(1, Ngen, (Ngen-1)), ylim=rangeYA, xlab="Generations", ylab=SelTrait, main=Main, cex.main=Main.Size)
  arrows(SRAbivar$data$gen,  SEaBot,   SRAbivar$data$gen, SEaTop, angle=90, length=0.05, code=3)

     # Plot the control if mentioned:
if(is.data.frame(control)) {
  NgenC<-max(control$gen)
  SEaC<-sqrt(control$phen.X.var/control$N)
  SEaTopC<-  control$phen.X.mean +SEaC
  SEaBotC<-  control$phen.X.mean -SEaC
  points(control$gen,control$phen.X.mean, col=col.cont, pch=16)
  lines(control$gen[1:NgenC], control$phen.X.mean[1:NgenC], lty=2, col=col.cont)
  arrows(control$gen,  SEaBotC,   control$gen, SEaTopC, angle=90, length=0.02, code=3, col=col.cont)
}

  lines(SRAbivar$data$gen[1:Ngen], SRAbivar$data$phen.X.mean[1:Ngen], lty=2, col=col.obs)
  lines(SRAbivar$data$gen[1:Ngen], SRAbivar$data$phen.X.mean[(Ngen+1):(Ngen*2)], lty=2, col=col.obs)
  
  lines(SRAbivar$data$gen[1:Ngen], SRAbivar$pred$D$phen.A, lty=1, col=col.pred)
  lines(SRAbivar$data$gen[1:Ngen], SRAbivar$pred$U$phen.A, lty=1, col=col.pred)
    

  
  ## Plot trait B (correlated)
SEb <- sqrt(SRAbivar$data$phen.Y.var/SRAbivar$data$N)
SEbTop<-  SRAbivar$data$phen.Y.mean +SEb
SEbBot<-  SRAbivar$data$phen.Y.mean -SEb
  rangeYB<-range(c(SRAbivar$data$phen.Y.mean, SRAbivar$pred$D$phen.B, SRAbivar$pred$U$phen.B, SEbTop, SEbBot))
  plot(SRAbivar$data$gen, SRAbivar$data$phen.Y.mean, type="p", pch=16, xaxp=c(1, Ngen, (Ngen-1)), ylim=rangeYB, xlab="Generations", ylab=CorTrait, cex.main=Main.Size)
  arrows(SRAbivar$data$gen,  SEbBot,   SRAbivar$data$gen, SEbTop, angle=90, length=0.05, code=3)

     # Plot the control if mentioned:
if(is.data.frame(control)) {
  NgenC<-max(control$gen)
  SEbC<-sqrt(control$phen.Y.var/control$N)
  SEbTopC<-  control$phen.Y.mean +SEbC
  SEbBotC<-  control$phen.Y.mean -SEbC
  points(control$gen,control$phen.Y.mean, col=col.cont, pch=16)
  lines(control$gen[1:NgenC], control$phen.Y.mean[1:NgenC], lty=2, col=col.cont)
  arrows(control$gen,  SEbBotC,   control$gen, SEbTopC, angle=90, length=0.02, code=3, col=col.cont)
}

    
  Ngen<-max(SRAbivar$data$gen)
  lines(SRAbivar$data$gen[1:Ngen], SRAbivar$data$phen.Y.mean[1:Ngen], lty=2, col=col.obs)
  lines(SRAbivar$data$gen[1:Ngen], SRAbivar$data$phen.Y.mean[(Ngen+1):(Ngen*2)], lty=2, col=col.obs)
  
  lines(SRAbivar$data$gen[1:Ngen], SRAbivar$pred$D$phen.B, lty=1, col=col.pred)
  lines(SRAbivar$data$gen[1:Ngen], SRAbivar$pred$U$phen.B, lty=1, col=col.pred)  
}


###################################################################################################

plot.sra.biv.Pred<-function(SRAbivar, SelTrait="Selected trait", CorTrait="Correlated trait", 
                        Main="Selection response",    
                        col.pred="blue", col.obs="black", Main.Size=1,
                        control=NA, col.cont="grey" ,
                        G.predA=NA,  G.predB=NA,
                        col.G.pred="red", col.G.pred.ci="salmon1"
                        ) {
  
  Ngen<-max(SRAbivar$data$gen)
  
  ## Plot trait A (selected)
SEa <- sqrt(SRAbivar$data$phen.X.var/SRAbivar$data$N)
SEaTop<-  SRAbivar$data$phen.X.mean +SEa
SEaBot<-  SRAbivar$data$phen.X.mean -SEa
if(is.data.frame(G.predA)){
  rangeYA<-range(c(SRAbivar$data$phen.X.mean, SRAbivar$pred$D$phen.A, SRAbivar$pred$U$phen.A, SEaTop, SEaBot, G.predA[,2], G.predA[,3]))
  plot(SRAbivar$data$gen, SRAbivar$data$phen.X.mean, type="n", pch=16, xaxp=c(1, Ngen, (Ngen-1)), ylim=rangeYA, xlab="Generations", ylab=SelTrait, main=Main, cex.main=Main.Size)

  polygon(x=c(1:Ngen,Ngen:1), y=c(G.predA[1:Ngen,2], G.predA[Ngen:1,3]), col=col.G.pred.ci, border=col.G.pred.ci)
  polygon(x=c(1:Ngen,Ngen:1), y=c(G.predA[(Ngen+1):(2*Ngen),2], G.predA[(2*Ngen):(Ngen+1),3]), col=col.G.pred.ci, border=col.G.pred.ci)
  segments(1:(Ngen-1), G.predA[1:(Ngen-1),1], 2:Ngen, G.predA[2:Ngen,1], col="red", lwd=1.5)
  segments(1:(Ngen-1), G.predA[(Ngen+1):(2*Ngen-1),1], 2:Ngen, G.predA[(Ngen+2):(2*Ngen),1], col=col.G.pred, lwd=1.5)


  points(SRAbivar$data$gen, SRAbivar$data$phen.X.mean, pch=16)
  arrows(SRAbivar$data$gen,  SEaBot,   SRAbivar$data$gen, SEaTop, angle=90, length=0.05, code=3)

  }
  
  
     # Plot the control if mentioned:
if(is.data.frame(control)) {
  NgenC<-max(control$gen)
  SEaC<-sqrt(control$phen.X.var/control$N)
  SEaTopC<-  control$phen.X.mean +SEaC
  SEaBotC<-  control$phen.X.mean -SEaC
  points(control$gen,control$phen.X.mean, col=col.cont, pch=16)
  lines(control$gen[1:NgenC], control$phen.X.mean[1:NgenC], lty=2, col=col.cont)
  arrows(control$gen,  SEaBotC,   control$gen, SEaTopC, angle=90, length=0.02, code=3, col=col.cont)
}

  lines(SRAbivar$data$gen[1:Ngen], SRAbivar$data$phen.X.mean[1:Ngen], lty=2, col=col.obs)
  lines(SRAbivar$data$gen[1:Ngen], SRAbivar$data$phen.X.mean[(Ngen+1):(Ngen*2)], lty=2, col=col.obs)
  
  lines(SRAbivar$data$gen[1:Ngen], SRAbivar$pred$D$phen.A, lty=1, col=col.pred, lwd=1.5)
  lines(SRAbivar$data$gen[1:Ngen], SRAbivar$pred$U$phen.A, lty=1, col=col.pred, lwd=1.5)
    

  
  ## Plot trait B (correlated)
SEb <- sqrt(SRAbivar$data$phen.Y.var/SRAbivar$data$N)
SEbTop<-  SRAbivar$data$phen.Y.mean +SEb
SEbBot<-  SRAbivar$data$phen.Y.mean -SEb
if(is.data.frame(G.predB)){
  rangeYB<-range(c(SRAbivar$data$phen.Y.mean, SRAbivar$pred$D$phen.B, SRAbivar$pred$U$phen.B, SEbTop, SEbBot, G.predB[,2], G.predB[,3]))
  plot(SRAbivar$data$gen, SRAbivar$data$phen.Y.mean, type="n", pch=16, xaxp=c(1, Ngen, (Ngen-1)), ylim=rangeYB, xlab="Generations", ylab=CorTrait, cex.main=Main.Size)
  polygon(x=c(1:Ngen,Ngen:1), y=c(G.predB[1:Ngen,2], G.predB[Ngen:1,3]), col=col.G.pred.ci, border=col.G.pred.ci)
  polygon(x=c(1:Ngen,Ngen:1), y=c(G.predB[(Ngen+1):(2*Ngen),2], G.predB[(2*Ngen):(Ngen+1),3]), col=col.G.pred.ci, border=col.G.pred.ci)
  segments(1:(Ngen-1), G.predB[1:(Ngen-1),1], 2:Ngen, G.predB[2:Ngen,1], col=col.G.pred, lwd=1.5)
  segments(1:(Ngen-1), G.predB[(Ngen+1):(2*Ngen-1),1], 2:Ngen, G.predB[(Ngen+2):(2*Ngen),1], col=col.G.pred, lwd=1.5)


  points(SRAbivar$data$gen, SRAbivar$data$phen.Y.mean, pch=16)

  arrows(SRAbivar$data$gen,  SEbBot,   SRAbivar$data$gen, SEbTop, angle=90, length=0.05, code=3)
  }
     # Plot the control if mentioned:
if(is.data.frame(control)) {
  NgenC<-max(control$gen)
  SEbC<-sqrt(control$phen.Y.var/control$N)
  SEbTopC<-  control$phen.Y.mean +SEbC
  SEbBotC<-  control$phen.Y.mean -SEbC
  points(control$gen,control$phen.Y.mean, col=col.cont, pch=16)
  lines(control$gen[1:NgenC], control$phen.Y.mean[1:NgenC], lty=2, col=col.cont)
  arrows(control$gen,  SEbBotC,   control$gen, SEbTopC, angle=90, length=0.02, code=3, col=col.cont)
}

    
  Ngen<-max(SRAbivar$data$gen)
  lines(SRAbivar$data$gen[1:Ngen], SRAbivar$data$phen.Y.mean[1:Ngen], lty=2, col=col.obs)
  lines(SRAbivar$data$gen[1:Ngen], SRAbivar$data$phen.Y.mean[(Ngen+1):(Ngen*2)], lty=2, col=col.obs)
  
  lines(SRAbivar$data$gen[1:Ngen], SRAbivar$pred$D$phen.B, lty=1, col=col.pred, lwd=1.5)
  lines(SRAbivar$data$gen[1:Ngen], SRAbivar$pred$U$phen.B, lty=1, col=col.pred, lwd=1.5)  
}


# Arnaud's functions for CI estimates:
do.profile.things <- function(param, minuslogL, p=c(0.05,0.001), ylim=range(-minuslogL), ylab="Log lik", xlab="Parameter", ...) { 
  # Param: the values of the parameter that have been tested 
  # minuslogL: minus log likelihood 
  # p: p-values associated to the desired CI (p=0.05 corresponds to a 95% CI, etc). 
  # the rest: options for plot() 
  
  plot(param, -minuslogL, type="l", ylim=ylim, ylab=ylab, xlab=xlab, ...) 
  
  # If the user wants CIs:  
  if (length(p) > 0) { 
    thresh <- qchisq(1-p, df=1)/2     
    # According to the standard likelihood analysis (p-values based on a likelihood ratio test) 
    
    # horizontal lines 
    abline(h=c(-min(minuslogL), -min(minuslogL)-thresh/2), lty=seq(1,length(p)+1), col="red") 
    
    for (tt in seq_along(thresh)) { 
      # Find the index of intersection points 
      intersec.index <- which(diff(sign(minuslogL - min(minuslogL) - thresh[tt])) != 0) 
      # Take the mean (very sloppy, at least, a linear regression would be better) 
      intersec <- sapply(intersec.index, function(i) {mean(param[c(i,i+1)])}) 
      # Draws the stuff on the plot 
      arrows(x0=intersec, y0=min(ylim), y1=-min(minuslogL)-thresh[tt], col="blue", lty=tt+1, code=0) 
      text(x=intersec, y=min(ylim)+0.2*diff(ylim), labels=format(intersec, digits=5), pos=2, srt=90) 
    } 
  } 
} 

do.profile.thingsCI <- function(param, minuslogL, p=0.05) { 
  # Param: the values of the parameter that have been tested 
  # minuslogL: minus log likelihood 
  # p: p-values associated to the desired CI (p=0.05 corresponds to a 95% CI, etc). 
  # the rest: options for plot() 
  
  # If the user wants CIs:  
  if (length(p) > 0) { 
    thresh <- qchisq(1-p, df=1)/2     
    # According to the standard likelihood analysis (p-values based on a likelihood ratio test) 
    
    # horizontal lines 
    
    for (tt in seq_along(thresh)) { 
      # Find the index of intersection points 
      intersec.index <- which(diff(sign(minuslogL - min(minuslogL) - thresh[tt])) != 0) 
      # Take the mean (very sloppy, at least, a linear regression would be better) 
      intersec <- sapply(intersec.index, function(i) {mean(param[c(i,i+1)])}) 
      
      if(length(intersec)==2){return(intersec)}
      if(length(intersec)==1){return(c(0,intersec))}      
    } 
  } 
} 
```
