## Supplementary material for "Unidirectional response to bidirectional selection on body size II Quantitative genetics": Data files and analysis scripts: Synthetic_maturity_data.R

```
#================================ DEFINITION OF PARAMETERS ==================================================================================================
# N = average number of fish per family
# ngen = number of generations
# nfam = number of families per line per generation
# maxage = maximum age of the fish (in days)
# nphen = number of phenotyping events in a fish life
# Line.levels = vector of selected lines 
# LF = asymptotic length in control treatment and high line
# g = growth rate parameter
# I = age at inflexion point
# LF_betas = line effects on LF growth parameter 
# g_betas = line effects on g growth parameter
# I_betas = line effects on I growth parameter
# Mat_beta0 = intercept for A line of maturity GLM
# Mat_beta1 = slope of age effect in maturity GLM
# Mat_beta2 = slope of length effect in maturity GLM
# Mat_beta3 = G line effect on intercept maturity GLM
# Mat_beta4 = P line effect on intercept maturity GLM
#=============================================================================================================================================================

Synthetic_maturity_data <- function(N, nfam, ngen, maxage, nphen, Line.levels, LF_beta0, LF_beta1, LF_beta2, LF_sigma, g_beta0, g_beta1, g_beta2, I_beta0, I_beta1, I_beta2, Mat_beta0, Mat_beta1, Mat_beta2, Mat_beta3,  Mat_beta4, Mat_sigma) {
#=====================================================================================================================================
#  1. BUILD DATASET
#=====================================================================================================================================
age <- seq(0, maxage, length.out = nphen);
Data <- expand.grid(Generation = c(1:ngen), Line = Line.levels, Family = c(1:nfam), ID = c(1:N), Age = age);
#=====================================================================================================================================
#  2. BUILD GROWTH CURVES WITH NOMALLY-DISTIBUTED RANDOM INDIVIDUAL AND GENERATION EFFECTS
#=====================================================================================================================================
### Design matrix for growth parameters
Xmat_growth <- model.matrix(~ Data$Line);	#Design matrix for the ANOVA, applied to each parameter
### Parameter vector for LF
LF.parm.vec <- c(LF_beta0, LF_beta1, LF_beta2); # Vector of paramaters on the response scale for "A", "G" and "P" lines
### Vector of residuals for LF
# Number of unique individuals in the experiment
Data$Unique <- paste(Data$Generation, Data$Line, Data$Family, Data$ID, sep = ".");
nbind <- length(levels(as.factor(Data$Unique)));
# Vector of individual-level residuals (half of the source of variance in residuals)
Data <- Data[order(Data$Unique),];
Data$LF.res.ind <- rep(rnorm(n = nbind, mean = 0, sd = sqrt(LF_sigma/2)), each = length(age)); #residuals, individual-wise
# Vector of generation-level residuals (half of the source of variance in residuals)
Data <- Data[order(Data$Generation),];
Data$LF.res.gen <- rep(rnorm(n = ngen, mean = 0, sd = sqrt(LF_sigma/2)), each = dim(Data[Data$Generation == 1,])[1]) #residuals, generation-wise
### Parameter vector for g
g.parm.vec <- c(g_beta0, g_beta1, g_beta2)
## Parameter vector for I
I.parm.vec <- c(I_beta0, I_beta1, I_beta2)
## Predicted length with the logistic growth model, using errors only on LF
Data$Length <- (Xmat_growth %*% LF.parm.vec + Data$LF.res.ind + Data$LF.res.gen)/(1 + exp((Xmat_growth %*% g.parm.vec) * (Xmat_growth %*% I.parm.vec - Data$Age)))
#=====================================================================================================================================
#  3. MATURITY STATUS WITH NOMALLY-DISTIBUTED RANDOM INDIVIDUAL AND GENERATION EFFECTS ON THE LOGIT SCALE
#=====================================================================================================================================
Xmat_mat <- model.matrix(~ Data$Age + Data$Length + Data$Line);	# Design matrix for the logistic regression
Mat.parm.vec <- c(Mat_beta0, Mat_beta1, Mat_beta2, Mat_beta3, Mat_beta4);	# Vector of parameters on a logit scale (logit link)
### Vector of residuals for maturity probability
# Vector of individual-level residuals (half of the source of variance in residuals)
Data <- Data[order(Data$Unique),];
Data$Mat.res.ind <- rep(rnorm(n = nbind, mean = 0, sd = sqrt(Mat_sigma/2)), each = length(age)); # Residuals, individual-wise
# Vector of generation-level (family-level) residuals (half of the source of variance in residuals)
#Data$fam <- paste(Data$Generation, Data$Line, Data$Family, sep = ".")
#Data <- Data[order(Data$fam),];
#Data$Mat.res.fam <- rep(rnorm(n = nfam, mean = 0, sd = sqrt(Mat_sigma/2)), each = dim(Data[Data$Family == 1,])[1]); # Residuals, family-wise
Data <- Data[order(Data$Generation),];
Data$Mat.res.gen <- rep(rnorm(n = ngen, mean = 0, sd = sqrt(Mat_sigma/2)), each = dim(Data[Data$Generation == 1,])[1]); # Residuals, generation-wise

invlogit <- function(x) {1/(1+exp(-x))}		# Inverse logit for back transform to a probability scale
Data$Maturity <- rbinom(n = dim(Data)[1], 1, p = invlogit(Xmat_mat %*% Mat.parm.vec + Data$Mat.res.ind + Data$Mat.res.gen));
# Loop through individuals to make maturity consistent (only 1s afer maturation)
unique <- levels(as.factor(Data$Unique));
for(i in 1:length(unique)){
  position <- match(1, Data$Maturity[Data$Unique == unique[i]])
  if (!is.na(position)) {Data$Maturity[Data$Unique == unique[i]][position:nphen] <- 1}
  }
return(Data[,-c(7,8,10:12)])
}
```
